# A Na^+^-selective receptor enables state-dependent plasticity in high-salt taste

**DOI:** 10.1101/2025.10.07.680871

**Authors:** Guangnan Qu, Bo Wang, Yan Chen

## Abstract

Sodium is an essential ion, but harmful in excess. Animals must suppress sodium aversion during deficiency, yet a cation-nonselective override would expose them to other harmful salts. Whether and how the sensory periphery regulates high-salt taste to achieve Na^+^-specific tuning remains unknown. Here we identify a Na^+^-selective, state-tunable high-salt detection mechanism in *Drosophila*. IR11a and IR25a function in bitter gustatory receptor neurons (GRNs) and are required for aversion specifically to high Na^+^. Loss of IR11a abolishes responses to Na^+^ and Li^+^ while sparing detection of K^+^, Ca^2+^, and bitter compounds. Heterologous expression confirms that IR11a and IR25a together confer Na^+^ responsiveness. During sodium deprivation, Na^+^ sensitivity in IR11a-expressing neurons is decreased and restored upon satiety, allowing flies to attenuate Na^+^ aversion specifically. The IR7c-dependent nonselective mechanism remains insensitive to sodium deprivation, thereby preserving aversion to other noxious cations even during sodium deficiency. The IR11a-mediated response is amiloride-sensitive, a functional signature reminiscent of mammalian ENaC despite the absence of sequence homology. This work reveals that peripheral high-salt taste combines a fixed, cation-nonselective defense with a sodium-selective, physiologically tunable mechanisms, offering a candidate logic for aversive high-salt taste in vertebrates, where the molecular basis remains poorly defined.

## Introduction

Na^+^ is an indispensable extracellular cation required for action potential generation, fluid homeostasis, and nutrient transport. However, excessive Na^+^ intake incurs substantial physiological costs, driving hyperosmotic stress and disrupting ionic homeostasis across species ^[1]^. To balance these conflicting constraints, animals have evolved a conserved bimodal taste system: low concentrations of Na^+^ (<100 mM) elicit appetitive behaviors to promote dietary intake, whereas high concentrations (>200 mM) trigger potent aversion to prevent toxicity ^[2, 3]^. This bimodal system serves well under stable conditions, but sodium availability fluctuates, and a sodium-deprived animal must ingest sodium even at concentrations that normally provoke aversion ^[4]^. State-dependent modulation of the appetitive low-salt mechanism has been reported during sodium deficiency, with enhanced attraction to low Na^+^ to drive intake, although the underlying mechanisms remain incompletely understood ^[5, 6]^. The aversive high-salt response, however, has been viewed as a rigid, cation-nonselective defense ^[4]^. This creates a physiological conflict: if suppression of high-salt aversion during deprivation were cation-nonselective, it would inevitably blunt aversion to other toxic cations as well. How the sensory periphery selectively attenuates sodium aversion without compromising defense against other noxious ions remains unknown.

In mammals, the appetitive mechanism for low Na^+^ is well-defined, relying on the amiloride-sensitive epithelial sodium channel (ENaC) in specialized taste receptor cells (TRCs) to provide Na^+^-selective attraction ^[7, 8]^. Conversely, high-salt aversion is distributed across bitter- and sour-sensing TRCs ^[9]^. This aversive response is broadly tuned to various cations and osmolarities via amiloride-insensitive mechanisms ^[9]^. Although several molecular candidates, including T2Rs, Car4, TMC4, and TRPV1, have been implicated in high-salt taste, they appear broadly tuned and their precise functional roles remain debated; none confers Na^+^-selective aversive taste ^[9–11]^. In line with this absence of a peripheral Na^+^-selective detector, rodents can increase their tolerance to aversive Na^+^ concentrations during sodium deprivation, but only through a central nervous system mechanism that is itself cation-nonselective^[4]^. Whether any animal has evolved a peripheral, Na^+^-selective high-salt receptor whose sensitivity is tuned by dietary sodium state thus remains an open question.

The fruit fly *Drosophila melanogaster* provides an exceptional model for decoding the molecular and cellular principles of salt taste ^[2]^. Flies deploy distinct sets of gustatory receptor neurons (GRNs) to mediate salt sensing ^[12]^. Ionotropic Receptors (IRs) expressed in these neurons play a major role in encoding taste quality ^[13]^. Specifically, low-salt taste is driven by IR56b in sweet GRNs and by IR20a in a separate GRN population; both receptors function with co-receptors IR25a and IR76b to provide Na^+^-selective attraction ^[14–16]^. In contrast, high-salt avoidance relies on parallel, molecularly distinct sensory mechanisms. Bitter GRNs and a specific population of glutamatergic GRNs expressing pickpocket 23 (*Ppk23^glut^*) both drive behavioral avoidance to concentrated salts ^[12]^. In the *Ppk23^glut^*GRNs, the tuning receptor IR7c works with IR25a and IR76b to non-selectively detect monovalent cations ^[17]^. Meanwhile, bitter GRNs execute high-salt responses through an IR76b-independent mechanism whose molecular mechanism has remained unidentified ^[12]^. Whether bitter GRNs mediate high-salt taste through IRs, and if so, whether this receptor confers Na^+^-selective aversion, remain unclear.

Here, we identify IR11a as the tuning subunit of an amiloride-sensitive, Na^+^-selective high-salt receptor. IR11a, alongside the obligate coreceptor IR25a, is expressed in a discrete subset of bitter GRNs and is selectively required for detecting high concentrations of Na^+^, but not K^+^ or bitter tastants. Heterologous expression confirms that IR11a and IR25a together are sufficient to confer Na^+^ responsiveness independent of IR76b. Importantly, Na^+^ sensitivity within *IR11a*-expressing GRNs exhibits state-dependent plasticity. Neuronal and behavioral responses to high Na^+^ are significantly dampened under salt-deprived conditions and re-established upon salt feeding, operating without transcriptional changes in the underlying IRs. This tunable, Na^+^-selective IR11a/IR25a mechanism operates alongside the statically tuned, cation-nonselective IR7c-dependent mechanism. Together, they enable selective attenuation of sodium aversion while preserving robust defense against other noxious salts.

## Results

### A behavioral screen identifies IR11a as a candidate high-salt taste receptor

To identify receptors contributing to high-salt detection, we conducted an RNAi based behavioral screen targeting ionotropic receptors (IRs), pickpocket (Ppk) family members, gustatory receptors (GRs), and transient receptor potential (TRP) channels, which are gene families previously implicated in *Drosophila* gustation ^[13, 18–20]^. We conditionally knocked down 64 candidate genes in adult flies using the pan-neuronal driver *nSyb-GAL4* combined with temperature-sensitive tubulin-GAL80^ts^ to avoid the potential effect of these genes in development, and we assessed high-salt avoidance with a two-choice feeding assay.

Our screen identified three genes whose pan-neuronal knockdown strongly reduced avoidance of 400 mM NaCl, including the previously characterized co-receptor IR25a and the high-salt tuning receptor IR7c ^[17]^, as well as the novel candidate IR11a (Figure 1A). Additionally, IR10a produced a weaker but statistically significant reduction (Figure 1A). However, knockdown of the remaining 60 candidates did not significantly affect avoidance (Figure S1). We focused our investigation on IR11a for further study, while IR10a was not pursued further in this study due to its weak phenotype.

**Fig. 1.**
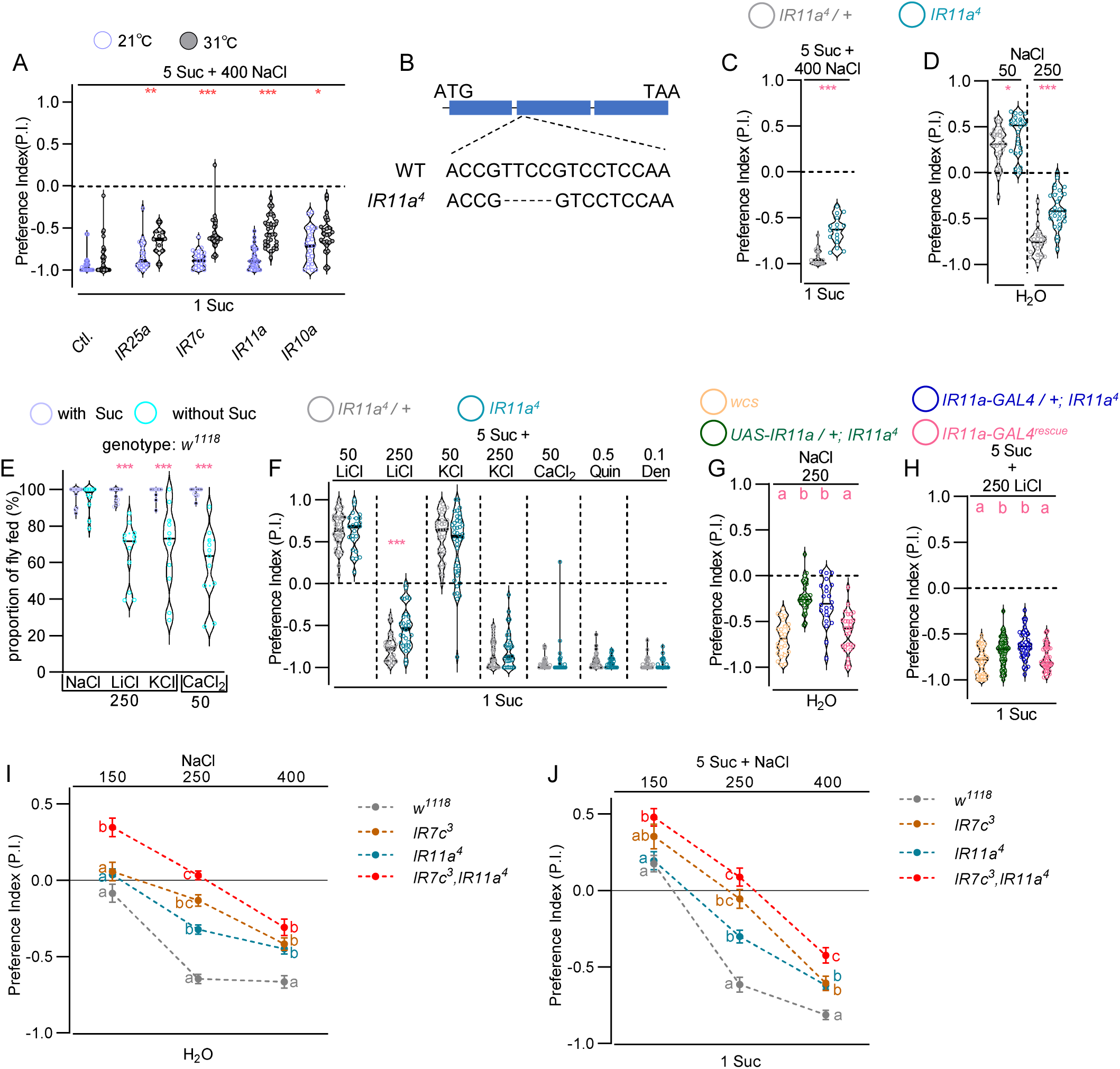
IR11a is selectively required for high Na^+^ and Li^+^ avoidance. **(A)** Pan-neuronal knockdown (*nSyb-GAL4/+; Tub-GAL80^ts^/+* > *UAS-RNAi*) of *IR25a*, *IR7c*, *IR11a*, and *IR10a* reduces high-Na^+^ avoidance. **(B)** Schematic of the *IR11a^4^* allele. **(C)** *IR11a^4^* mutants show reduced avoidance of 400 mM NaCl compared to controls. **(D)** *IR11a^4^* mutants show increased attraction to 50 mM NaCl and reduced avoidance of 250 mM NaCl compared to controls. **(E)** Proportion of flies fed on sucrose-supplemented versus unsupplemented salts across genotypes. **(F)** *IR11a^4^* mutants show reduced avoidance of high LiCl, while avoidance of other tested tastants remains intact. **(G - H)** Avoidance of 250 mM NaCl and LiCl is restored by genetic rescue with IR11a. **(I - J)** Epistasis analysis of *IR7c* and *IR11a* in Na^+^ and Li^+^ avoidance. Concentrations are in mM. Data are mean ± SEM (n = 10 - 45). Mann-Whitney test for **(A - F)**; *p<0.05, **p<0.01, ***p<0.001. Kruskal-Wallis with Dunn’s multiple comparisons test for **(G - J)**; different letters indicate p<0.05. Genotypes and statistical details are provided in Data S1. See also Fig. S1.

### IR11a is selectively required for behavioral avoidance of Na^+^ and Li^+^

To validate the RNAi screening results, we generated an *IR11a* loss-of-function mutant (*IR11a^4^*) via CRISPR/Cas9-mediated deletion of four nucleotides (TTCC at positions 173 to 176 of exon 2), which results in a coding frameshift (Figure 1B). In two-choice feeding assays, *IR11a* mutants displayed reduced avoidance of 400 mM NaCl plus 5 mM sucrose versus 1 mM sucrose (Figure 1C). To exclude potential confounding effects of sucrose on salt perception, we tested NaCl versus plain agarose. Under these conditions, *IR11a* mutants showed reduced aversion to 250 mM NaCl and a weak but statistically significant increase in attraction to 50 mM NaCl (Figure 1D). This observation suggests that sodium attraction pathways might be unmasked or compensatorily enhanced when the IR11a-dependent aversive pathway is compromised.

A significantly higher proportion of flies fed on tastants supplemented with sucrose than on tastants alone. We observed this effect for KCl, LiCl, CaCl_2_, quinine, and denatonium, but not for NaCl (Figure 1E). Therefore, we used tastant with 5 mM sucrose versus 5 mM sucrose alone to assess aversion to all other tastants. *IR11a* mutants exhibited selectively reduced avoidance of 250 mM LiCl, whereas responses to KCl, CaCl_2_, the bitter alkaloid quinine, and the synthetic bitter compound denatonium remained indistinguishable from controls (Figure 1F). Targeted expression of IR11a using *IR11a-GAL4* ^[21]^ restored avoidance of 250 mM NaCl and LiCl to wild-type levels (Figures 1G and 1H), confirming that the behavioral deficits are caused by loss of IR11a function.

To determine whether IR11a acts independently of IR7c, which has been implicated in high-salt detection^[17]^, we evaluated the feeding behavior of single and double mutants at intermediate (150 mM) and high (250 and 400 mM) NaCl concentrations. At 150 mM NaCl, wild-type controls and single mutants displayed roughly neutral responses, whereas *IR7c;IR11a* double mutants exhibited an anomalous salt preference (Figure 1I and 1J). As the concentration increased to 250 and 400 mM, both *IR11a* and *IR7c* single mutants showed significantly reduced salt avoidance compared to wild-type flies, a deficit that was profoundly exacerbated in the double mutants (Figure 1I and 1J). This additive phenotype confirms that IR11a and IR7c function in genetically separable, parallel mechanisms for high-salt detection.

### IR11a is expressed in a discrete subset of bitter GRNs

Next, we analyzed the expression pattern of *IR11a-GAL4* to identify the anatomical sites of IR11a function. In the labellum, which is the primary external taste organ of *Drosophila*, *IR11a-GAL4* consistently labeled GRNs in sensilla S4, S6, and S10 (Figure 2A1 and S2A). Additional labeling in S8 sensilla was observed in approximately 40% of preparations (Figure 2D1 and S2A). In the legs, expression was restricted to the f5s sensillum of the foreleg tarsi (Figure 2A2 and S2B). We also observed two labeled neurons in the ventral cibarial sense organ (VCSO) of the pharynx (Figures 2A3). No expression was detected in the brain, ventral nerve cord, gut, or Malpighian tubules (Figures S2C and S2D), indicating that IR11a is confined to peripheral gustatory tissues.

**Fig. 2.**
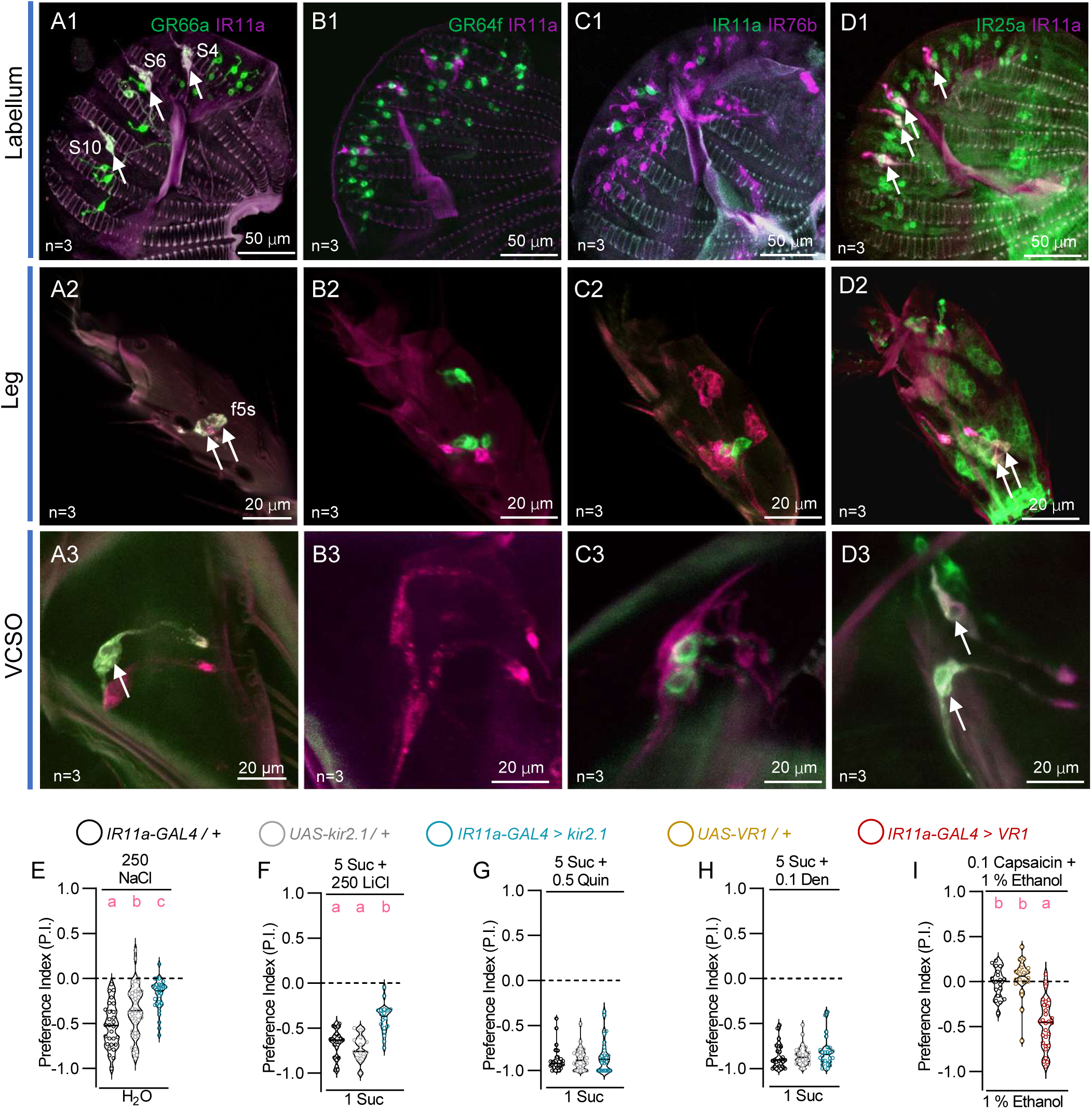
IR11a is expressed in a subset of bitter GRNs that mediate high NaCl and LiCl avoidance. **(A - D)** Co-localization analysis of *IR11a-GAL4* with *GR66a-LexA* (bitter GRNs), *GR64f-LexA* (sweet GRNs), *IR76b-QF* (co-receptor), and IR25a (co-receptor; using IR25a antibody) in labellum, legs, and ventral cibarial sense organ (VCSO). Pseudo-coloring was applied to facilitate identification of labeled cells. Arrowheads indicate co-expression. **(E - H)** Silencing *IR11a*-expressing GRNs reduces avoidance of high NaCl **(E)** and LiCl **(F)**, but does not affect avoidance of quinine **(G)** or denatonium **(H)**. **(I)** Ectopic expression of the capsaicin receptor VR1 in *IR11a*-expressing GRNs induces capsaicin avoidance. Concentrations are in mM. Data are mean ± SEM (**E - I**, n = 13-39). Kruskal-Wallis with Dunn’s multiple comparisons test; different letters indicate p<0.05. Genotypes and statistical details are provided in Data S2. See also Fig. S2.

To determine the cellular identity of *IR11a*-expressing GRNs, we performed co-expression analyses with cell-type specific markers. *IR11a-GAL4* showed complete co-localization with *GR66a-LexA* ^[22]^, a marker for bitter GRNs, in labellar and tarsal sensilla (Figure 2A). Partial co-localization was observed in the VCSO, where one of the two *IR11a*-expressing neurons co-expressed *GR66a-LexA* (Figure 2A). All *IR11a*-expressing GRNs co-expressed the broad IR co-receptor IR25a (Figure 2D). However, these neurons did not co-express *IR76b-QF* ^[23]^ or the sweet GRN marker *GR64f-LexA* ^[24]^ (Figures 2B and 2C). These data indicate that IR11a defines a distinct subset of IR25a and GR66a positive bitter GRNs. Bitter GRNs were previously shown to respond to high salt through IR76b independent mechanisms [6], and our findings identify IR11a as the likely tuning receptor in this context.

To assess the functional role of *IR11a*-expressing GRNs, we silenced them by expressing the inward rectifying K^+^ channel Kir2.1 ^[25]^ under the control of *IR11a-GAL4* and conducted two-choice feeding assays. Silencing these GRNs reduced avoidance of 250 mM NaCl and LiCl, but it had no effect on avoidance of quinine or denatonium (Figures 2E to H). Conversely, artificial activation of these GRNs via ectopic expression of the capsaicin receptor VR1 ^[26]^ induced robust avoidance of capsaicin laced food (Figure 2I). This phenotype indicates that *IR11a*-expressing GRNs are necessary for sodium and lithium avoidance and sufficient to drive aversive behavior. The lack of effect on bitter avoidance does not rule out a role for the *IR11a*-expressing GRNs in bitter sensing, as other bitter GRNs likely provide sufficient function to sustain avoidance of quinine and denatonium.

### IR11a and IR25a are required for Na**^+^**-selective responses in *IR11a*-expressing GRNs

We next characterized the physiological response properties of *IR11a*-expressing GRNs and the specific contributions of IR11a and its co-receptors. We performed single-cell Ca^2+^ imaging using GCaMP6s in tarsal *IR11a*-expressing GRNs, labeled by *GR33a^GAL4^*^[27]^, which is expressed in the f5s sensillum. These neurons exhibited dose-dependent Ca^2+^ responses to NaCl concentrations ranging from 150 to 500 mM, reaching a plateau at 250 mM (Figures 3A and 3B). Application of a broader panel of stimuli revealed that these neurons broadly responded to 250 mM LiCl, NaBr, and KCl, as well as 50 mM CaCl_2_ and 1 mM denatonium. Howeveer, they did not respond to 250 mM NMDG-Cl, confirming that salt responses are cation-dependent (Figure 3C).

**Fig. 3.**
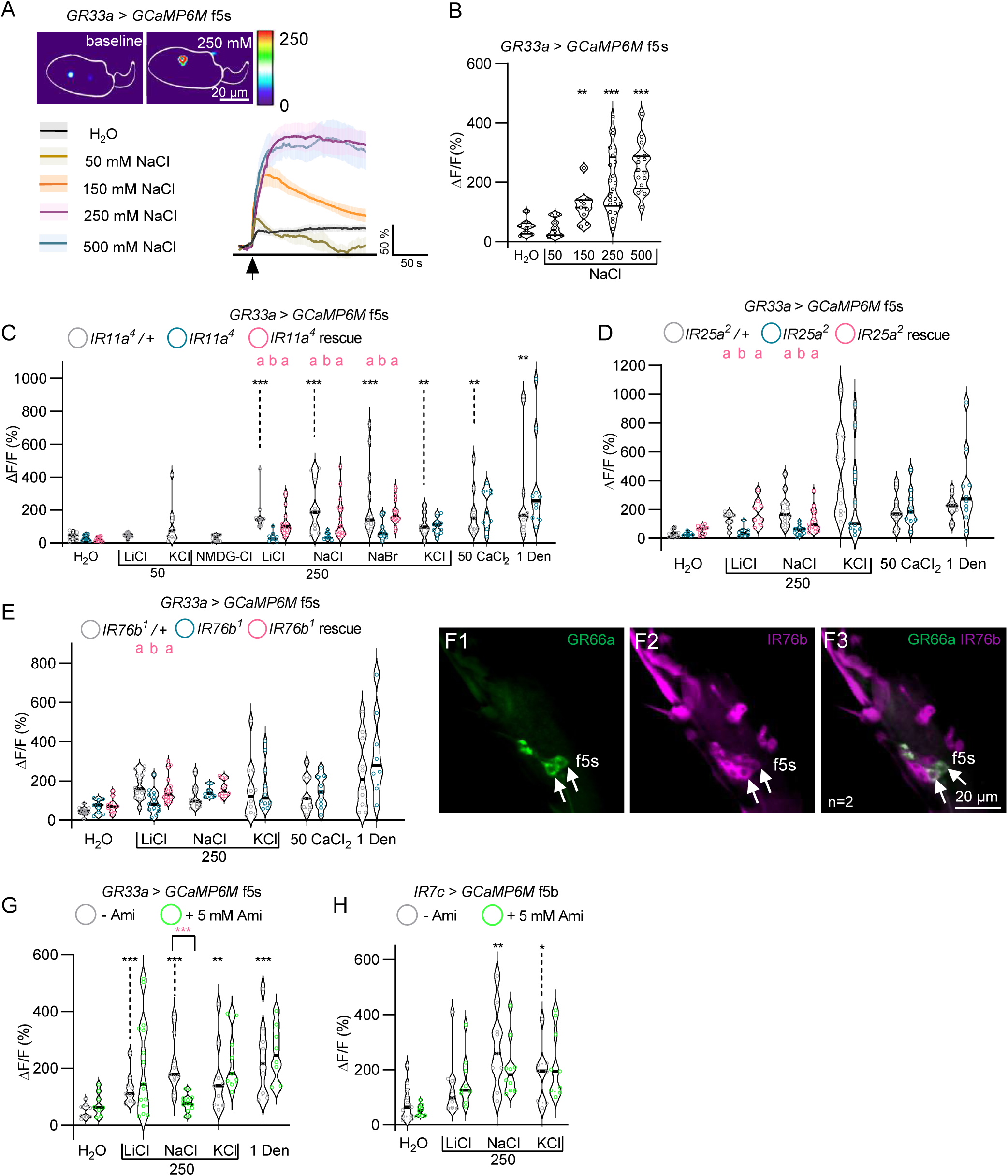
IR11a and IR25a mediate high Na^+^ and Li^+^ detection through distinct pharmacological and molecular requirements. **(A - B)** Calcium responses of *IR11a*-expressing tarsal GRNs to 50 - 500 mM NaCl. **(A)** Heatmap and response traces. **(B)** Quantification. **(C - E)** Calcium responses of *IR11a*-expressing tarsal GRNs (labeled by *GR33a^GAL4^*, f5s) to LiCl, NaCl, NMDG-Cl, NaBr, KCl, CaCl_2_, and denatonium in *IR11a* (C), *IR25a* (D), and *IR76b* (E) mutants and rescue flies. **(F)** Co-expression analysis of *GR66a-LexA* and *IR76b-GAL4* in the tarsal 5s sensillum; arrowheads indicate co-expression. **(G - H)** Effect of amiloride on LiCl, NaCl, KCl, and denatonium responses in *IR11a*-expressing GRN. Black asterisks indicate responses differing from H_2_O in control flies (p<0.05). Red asterisks denote significant differences between designated groups. Concentrations are in mM. Data are mean ± SEM (n = 8 - 24). Mann-Whitney test for (**B, C, G, H**) comparing tastant responses with H_2_O in control flies; *p<0.05, **p<0.01, ***p<0.001. Dunn’s multiple comparisons test for **(C - E)** among control, mutant, and rescue groups; different letters indicate p<0.05. Genotypes and statistical details are provided in Data S3. See also Fig. S3.

While the *IR11a*-expressing GRNs are broadly tuned, *IR11a* mutants displayed a selective physiological deficit. Ca^2+^ responses to Na^+^ and Li^+^ salts were severely reduced, whereas responses to KCl, CaCl_2_, and denatonium remained intact (Figure 3C). This specific deficit was fully rescued by the reintroduction of IR11a (Figure 3C). Axonal Ca^2+^ imaging in the subesophageal zone (SEZ) confirmed that labellar *IR11a*-expressing GRNs exhibited a similar Na^+^ selective dependence on IR11a (Figures S3A to S3C). However, these labellar neurons lacked LiCl sensitivity (Figures S3A to S3C), pointing toward tissue-specific differences in receptor subunit composition.

We then examined the requirement for the co-receptors IR25a and IR76b. In *IR25a* mutants ^[28]^, responses to both NaCl and LiCl were selectively reduced without affecting KCl, CaCl_2_, or denatonium responses (Figure 3D). Neuron-specific expression of IR25a fully rescued these deficits (Figure 3D). This mirrors the *IR11a* mutant phenotype and indicates that IR25a functions as a required co-receptor for the IR11a dependent salt detection complex.

In contrast, *IR76b* mutants ^[23]^ showed a more restricted phenotype where only LiCl responses were reduced, while NaCl, KCl, CaCl_2_, and denatonium responses remained normal (Figure 3E). Because our initial expression analysis did not detect *IR76b-QF* in *IR11a*-expressing GRNs (Figure 2C), we re-examined IR76b expression using *IR76b-GAL4* ^[28]^ driver combined with *GR66a-LexA,* which labels *IR11a*-expressing GRNs in tarsal f5s sensilla. Co-expression of *IR76b-GAL4* and *GR66a-LexA* was confirmed in these sensilla, suggesting that IR76b is co-expressed with IR11a (Figure 3F). These findings reveal that in tarsal GRNs, where IR76b is co-expressed, both Na^+^ and Li^+^ are detected (Figure 3C). In labellar GRNs, where IR76b is absent, only Na^+^ detection requires IR11a (Figure S3C). We therefore propose that IR11a and IR25a form the core Na^+^ detection complex, while IR76b serves as an accessory subunit required for Li^+^ sensitivity in specific tissues.

We also tested whether IR11a dependent Na^+^ responses are sensitive to amiloride, an inhibitor of the vertebrate Na^+^-selective low-salt receptor ENaC ^[7]^. Application of 100 μM amiloride selectively reduced NaCl-evoked Ca^2+^ responses in tarsal *IR11a*-expressing GRNs, with no effect on LiCl or KCl responses (Figure 3G). In contrast, amiloride had no effect on NaCl, LiCl, or KCl responses in *IR7c*-expressing GRNs (Figure 3H). This pharmacological distinction provides further evidence that the IR11a and IR7c represent molecularly distinct Na^+^ detection mechanisms.

### Ectopic expression in pheromone**-**sensing neurons confer Na^+^-selective sensitivity

To determine whether IR11a is sufficient to confer Na^+^ sensitivity to otherwise salt-insensitive neurons, we ectopically expressed IR11a in pheromone-sensing tarsal f5a GRNs using *Ppk23-GAL4* ^[22]^. *Ppk23-GAL4* is expressed in four GRNs in the f5a sensillum; all of them endogenously express IR25a, but only two of them express IR76b (Figures 4A and 4B) ^[16]^. Previous studies established that the *Ppk23-GAL4* f5a neurons do not respond to sugars, bitter compounds, or acids ^[16, 22, 29]^, a result we confirmed for NaCl, LiCl, and KCl (Figure 4C). Misexpression of IR11a in *Ppk23*-expressing GRNs selectively conferred Ca^2+^ responses to 250 mM NaCl, without inducing sensitivity to LiCl or KCl (Figure 4C). This acquired NaCl response was abolished in *IR25a* mutants but remained functional in *IR76b* mutants (Figures 4D and 4E). These data demonstrate that IR11a and IR25a are sufficient to confer Na^+^ sensitivity to heterologous neurons independent of IR76b.

**Fig. 4.**
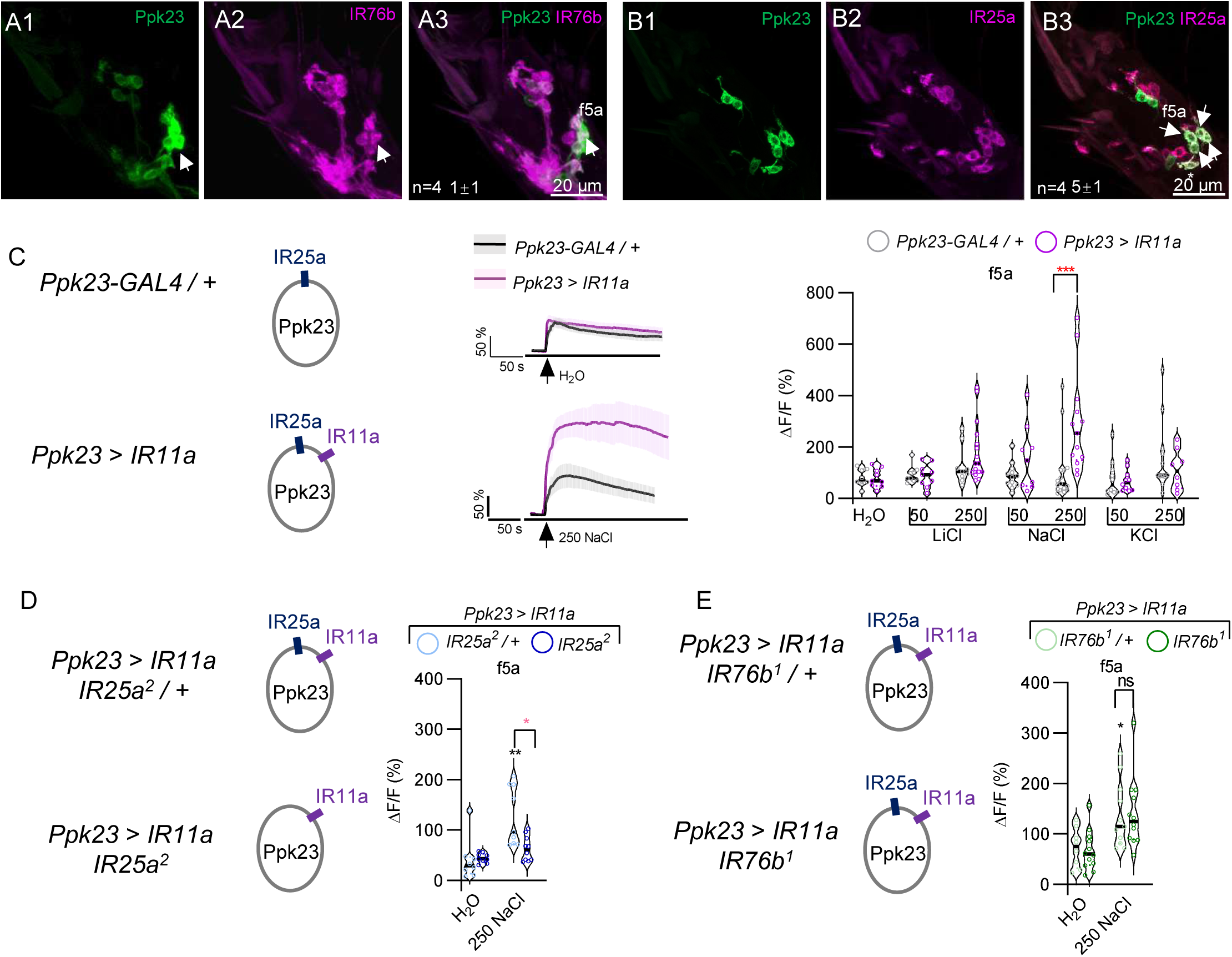
Ectopic expression of IR11a confers IR25a-dependent, IR76b-independent NaCl sensitivity to pheromone-sensing GRNs. **(A - B)** Co-expression analysis of *Ppk23-GAL4* with *IR76b-QF* and with IR25a (using IR25a antibody) in f5a sensilla. Arrowheads indicate co-expression. **(C)** Calcium responses to LiCl, NaCl, and KCl in *Ppk23-GAL4*-labeled f5a GRNs that ectopically express IR11a. **(D - E)** Calcium responses to LiCl, NaCl, and KCl of *Ppk23-GAL4*-labeled f5a GRNs that ectopically express IR11a in *IR25a* control and mutant backgrounds **(D)**, or in *IR76b* control and mutant backgrounds **(E)**. Concentrations are in mM. Data are mean ± SEM (n = 8 - 16). Mann-Whitney test; *p<0.05, **p<0.01, ***p<0.001. Red asterisks denote significant differences between designated groups in (**C, D**); black asterisks indicate responses differing from H_2_O in control flies (p<0.05) in (**D, E**). Genotypes and statistical details are provided in Data S4.

### Heterologous expression in S2 cells reveals subunit-dependent ionic tuning

Next, we expressed IR11a and IR25a, either alone or together with IR76b, in *Drosophila* S2 cells and measured Ca^2+^ responses to salt stimulation to directly assess whether they are sufficient to confer Na^+^-selective sensitivity, and to examine the contribution of IR76b. Cells expressing either IR11a or IR25a alone showed no responses to 250 mM NaCl, LiCl, or KCl (Figures 5A to 5C). Co-expression of IR11a and IR25a conferred Na^+^-selective Ca^2+^ responses, with cells responding to 250 mM NaCl but not to equimolar LiCl or KCl (Figures 5A, 5D, and 5E).

**Fig. 5.**
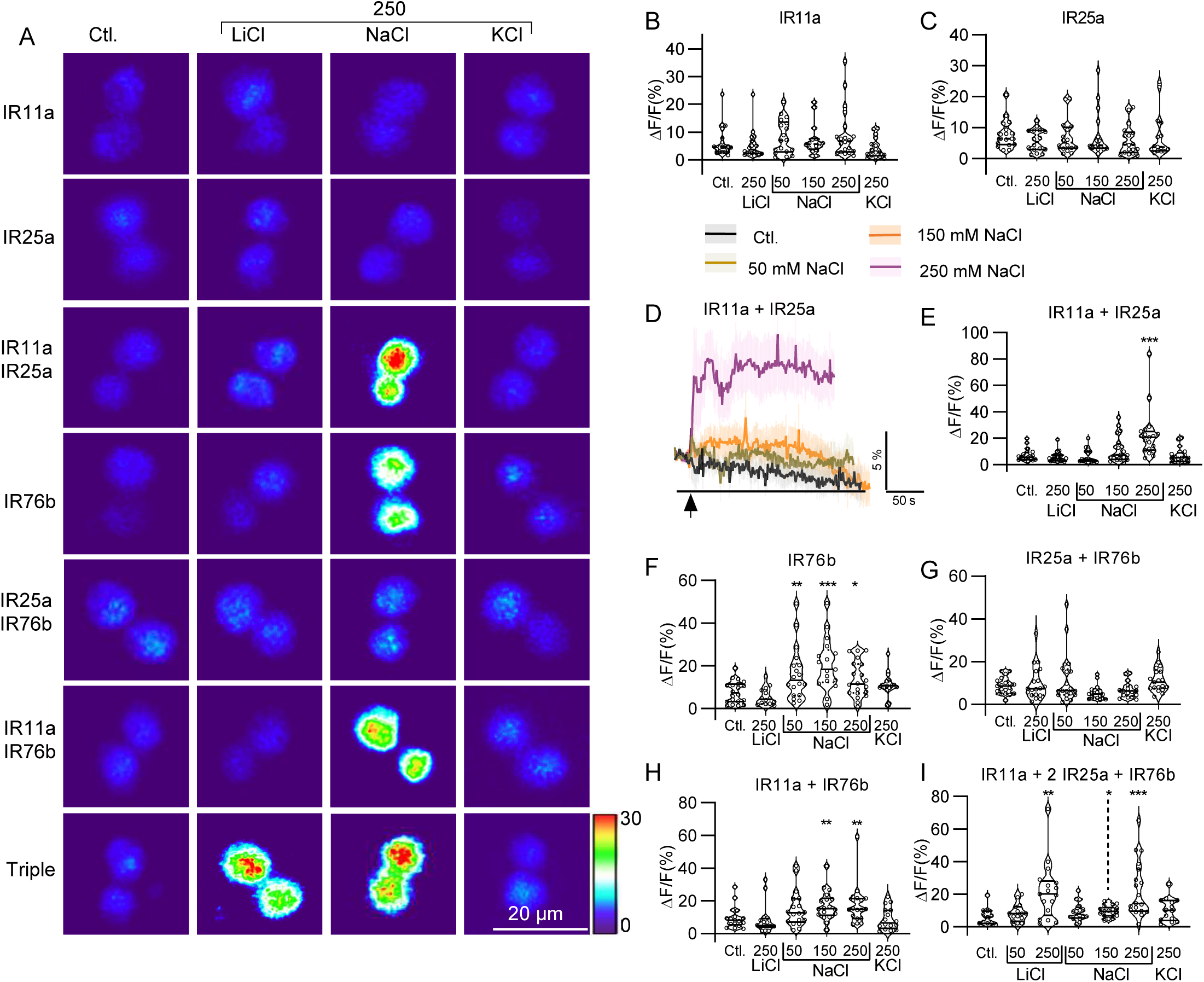
IR11a and IR25a together confer Na^+^-selective high-salt sensitivity, with cation and concentration range further shaped by IR76b. (A) Heat maps of calcium responses in S2 cells expressing different IR combinations. **(B - E)** Calcium responses to LiCl, NaCl, and KCl in S2 cells that express IR11a, IR25a, or both. **(F - I)** Calcium responses to LiCl, NaCl, and KCl in S2 cells expressing IR76b alone, IR76b with IR25a, IR76b with IR11a, or the triple (IR11a, IR76b, and IR25a) combination. Concentrations are in mM. Data are mean ± SEM (n = 11 - 20). Mann-Whitney test; *p<0.05, **p<0.01, ***p<0.001. Genotypes and statistical details are provided in Data S5.

Consistent with previous reports ^[16, 23]^, expression of IR76b alone conferred NaCl sensitivity in S2 cells (Figures 5A and 5F). Co-expression of IR25a with IR76b abolished IR76b mediated Na^+^ responses at all tested concentrations (Figure 5G). Co-expression of IR11a with IR76b abolished IR76b mediated responses to 50 mM NaCl while preserving responses to 150 mM and 250 mM NaCl (Figure 5H) ^[16]^. In contrast, triple expression of IR11a, IR25a, and IR76b enabled responsiveness to 150 mM NaCl, 250 mM NaCl and LiCl, but not to KCl (Figure 5I). This pattern suggests that IR76b expands the ionic sensitivity of the IR11a/IR25a complex to include Li^+^ and to lower NaCl concentrations.

Taken together, our heterologous expression experiments show that IR11a and IR25a are sufficient to confer Na^+^-selective responses, while IR76b serves as a modulatory subunit that broadens ionic sensitivity.

### IR11a and IR25a are required for state-dependent modulation of Na^+^ sensitivity

The independent functions of IR11a and IR7c prompted us to investigate their distinct physiological roles in high-salt taste. We hypothesized that the Na^+^ selectivity of IR11a allows flies to selectively adjust Na^+^ consumption dependent on dietary sodium state. To test this hypothesis, we assessed behavioral avoidance of 250 mM NaCl in flies subjected to either a sodium-deprived or a sodium-fed diet. Wild-type flies exhibited significantly reduced avoidance of NaCl when salt-deprived (Figures 6A and 6B). Such state-dependent modulation was completely abolished in *poxn* mutants (Figure 6A), which lack functional peripheral taste sensilla but retain intact pharyngeal taste neurons ^[30]^. The absence of modulation in *poxn* mutants implies that peripheral GRNs play important role in state-dependent high-salt avoidance.

**Fig. 6.**
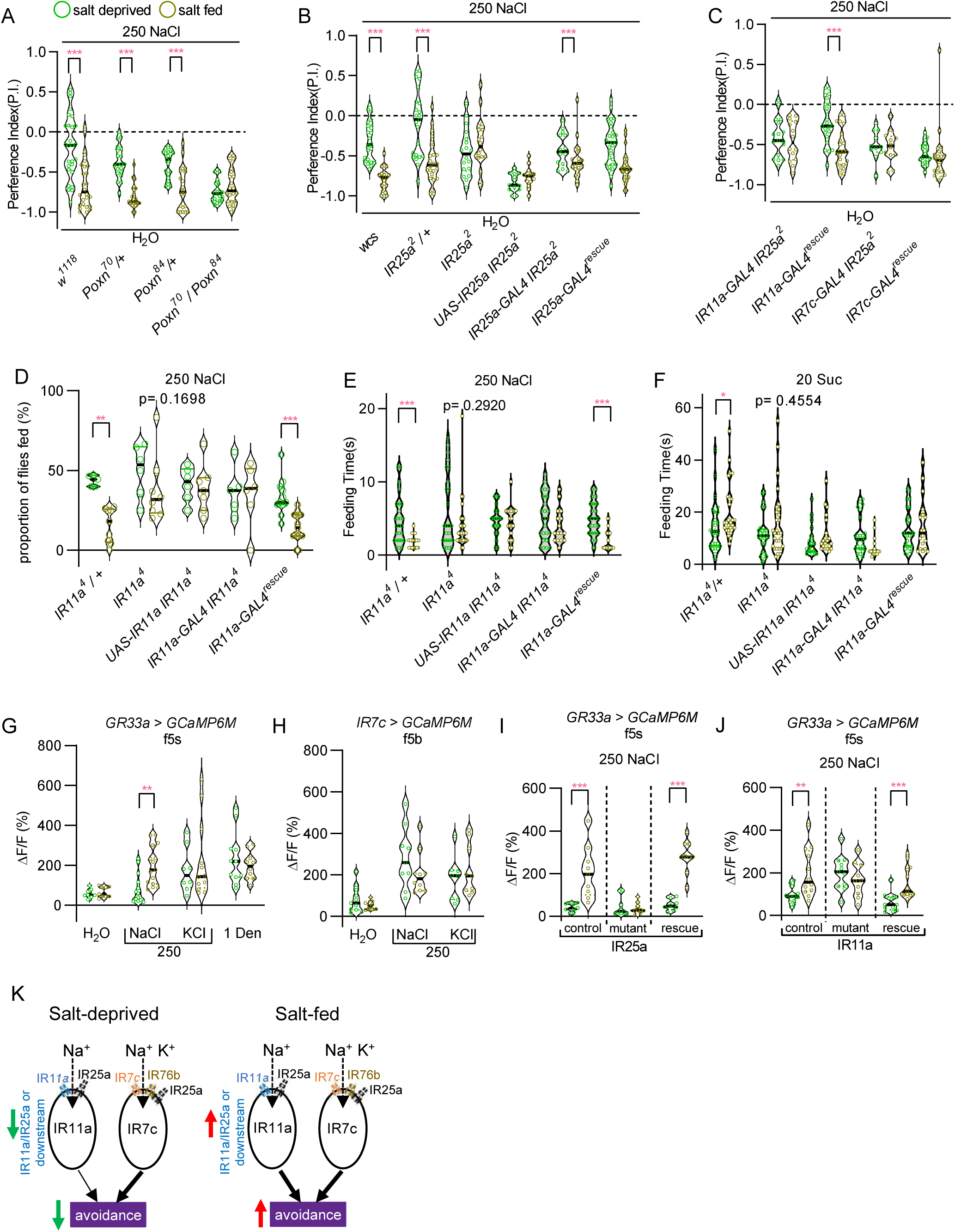
IR11a and IR25a are required for state-dependent plasticity of high-salt feeding. **(A)** Avoidance of 250 mM NaCl in controls and *poxn* mutants under salt-fed and salt-deprived conditions. **(B - C)** Avoidance of 250 mM NaCl in controls, *IR25a* mutants, and cell-type-specific rescue genotypes under salt-fed and salt-deprived conditions. **(D)** Proportion of *IR11a* controls, mutants, and rescue flies feeding on 250 mM NaCl under salt-fed and salt-deprived conditions. **(E - F)** Feeding duration on 250 mM NaCl and 20 mM sucrose in *IR11a* controls, mutants, and rescue flies under salt-fed and salt-deprived conditions. **(G - H)** Calcium responses to NaCl in *IR11a*-expressing GRNs (labeled by *GR33a^GAL4^*, 5s) **(G)** and *IR7c*-expressing GRNs **(H)** under salt-deprived and salt-fed conditions. **(I - J)** Calcium responses of *IR11a*-expressing GRNs to NaCl in *IR25a* and *IR11a* controls, mutants, and rescue flies under salt-deprived and salt-fed conditions. **(J)** Working model: the IR11a-dependent Na^+^-selective receptor and the IR7c-dependent cation-nonselective receptor operate in parallel to enable state-dependent modulation of sodium aversion. Concentrations are in mM. Data are mean ± SEM (n = 8 - 15 for calcium imaging; n = 5 - 29 for two-choice feeding assays and MAFE assays). Mann-Whitney test; *p<0.05, **p<0.01, ***p<0.001. Genotypes and statistical details are provided in Data S6. See also Fig. S4.

To identify the molecular components involved, we examined the requirement for IR25a, which serves as an essential co-receptor for both the IR11a and IR7c tuning receptors. *IR25a* mutants failed to show state-dependent modulation of high-salt avoidance (Figure 6B). The behavioral defect was successfully rescued by reintroducing IR25a under the control of *IR25a-GAL4* (Figures 6B). To identify the specific GRN population in which IR25a mediates behavioral plasticity, we performed cell-type-specific rescue experiments. Expression of IR25a in *IR11a*-expressing GRNs fully restored state-dependent modulation in *IR25a* mutants, whereas expression in *IR7c*-expressing GRNs had no effect (Figure 6C). The selective rescue establishes that *IR11a*-expressing GRNs, rather than *IR7c*-expressing GRNs, are required for state-dependent high-salt avoidance.

We further employed the manual feeding (MAFE) assay ^[31]^ to directly assess the impact of IR11a on feeding behavior at single-fly resolution. Wild-type flies displayed a lower feeding proportion and reduced feeding duration on 250 mM NaCl under salt-fed conditions compared to salt-deprived conditions, but showed no state-dependent difference when offered 20 mM sucrose (Figures 6D to 6F). *IR11a* mutants exhibited similar feeding behavior for NaCl under both conditions, failing to adjust intake according to internal state (Figures 6D to 6E). The feeding adjustment failure in mutant flies was fully rescued by IR11a re-expression (Figures 6D to 6E). The MAFE assay results demonstrate that IR11a is required for adjusting sodium consumption in response to dietary sodium state.

To determine the cellular basis of behavioral plasticity, we performed Ca^2+^ imaging to examine whether the Na^+^ sensitivity of *IR11a*-expressing GRNs is modulated by dietary sodium state. *IR11a*-expressing GRNs, labeled by *GR33a^GAL4^*, exhibited reduced NaCl-evoked activity in salt-deprived flies compared to salt-fed controls, whereas responses to KCl and denatonium remained unchanged (Figure 6G). In contrast, *IR7c*-expressing GRNs maintained stable NaCl responses regardless of salt status (Figure 6H), consistent with the inability of *IR7c-GAL4* driven IR25a expression to rescue state-dependent behavior (Figure 6C). Mutant analysis revealed that the state-dependent Na^+^ sensitivity change in *IR11a*-expressing GRNs was lost in both *IR11a* and *IR25a* mutants, and normal physiological modulation was restored by re-expressing the respective receptor genes (Figures 6I and 6J).

To determine whether state-dependent modulation involves changes in receptor expression, we measured transcript and protein levels of key IR subunits. Quantitative PCR analysis revealed no significant differences in *IR11a*, *IR25a*, *IR76b*, or *IR7c* transcript levels between salt-deprived and salt-fed conditions (Figure S4A). Immunostaining with anti-IR25a antibodies ^[28]^ showed no detectable changes in IR25a protein levels in *IR11a*-expressing GRN cell bodies (Figures S4B to S4E). Because of a lack of specific antibodies for IR11a, we were unable to test whether IR11a protein levels are modulated by salt state. Nevertheless, the combined expression data suggest that state-dependent modulation likely occurs at a post-transcriptional level.

## Discussion

High-salt taste has traditionally been viewed as a broadly tuned, cation-nonselective defensive mechanism rigidly hardwired against homeostatic demand ^[4, 9, 17]^. Here, we show that *Drosophila* IR11a and IR25a form a functional Na^+^-selective receptor that drives high-salt aversion and is subject to modulation by dietary sodium status. This finding reveals that taste neurons possess a previously unappreciated capacity for selective, state-dependent tuning and provides the first evidence in any animal that high-salt avoidance can be dynamically adjusted by the sodium homeostatic state via the high-salt taste receptor.

### Molecular determinants of IR11a-mediated Na^+^ selectivity and sensitivity

We demonstrate that IR11a and IR25a are necessary for Na^+^-selective high-salt responses in bitter GRNs, and sufficient when ectopically expressed in either pheromone-sensing neurons or cultured cells. Furthermore, incorporation of IR76b expands the dynamic range to include Li^+^ and lower concentrations of NaCl (150 mM), potentially enabling animals to adjust the strength of sodium avoidance according to their dietary state. Given that the ionic radius of Li^+^ is smaller than that of Na^+^, the Li^+^ permeability of the IR11a/IR25a/IR76b complex likely arises as a biophysical byproduct of a pore architecture strictly optimized to discriminate Na^+^ from larger cations such as K^+^; the negligible environmental concentration of Li^+^ renders this leak functionally inconsequential.

### Adaptive and static peripheral coding of high salt taste

Salt feeding regulation relies on a multi-tiered hierarchy that integrates physiological need. In *Drosophila*, state-dependent plasticity has been reported in appetitive salt taste GRNs, where sodium-enriched diets attenuate sweet GRN activity by modulating IR56b-dependent low-salt detection or by driving internalization of gustatory receptor components^[6, 32]^. However, the dynamic modulation of aversive high-salt taste has largely been attributed to post-peripheral checkpoints, such as downstream circuits of *Ppk23^glut^*GRNs, pharyngeal LSO neurons, and central nutrient-sensing networks ^[12, 33, 34]^. The identification of the IR11a-mediated complex expands this paradigm, establishing that external aversive GRNs themselves serve as active substrates for adaptive modulation. By selectively dampening the IR11a-mediated aversive signal during Na^+^ depletion, the fly can transiently attenuate its first line of behavioral defense specifically for sodium to satisfy physiological demand. Meanwhile, the statically tuned IR7c-dependent receptor ensures continuous, non-selective protection against other noxious environmental salts. Thus, the IR11a-dependent Na^+^-selective receptor and the IR7c-dependent cation-nonselective receptor operate in parallel: the former delivers a sodium-specific, adaptive tuning that aligns intake with physiological demand, while the latter mediate a generalist aversion to diverse salts (Figure 6K).

### Ecological and evolutionary implications

Ecologically, although rotting fruit, the main *Drosophila* food source in nature, is typically Na^+^-poor and K^+^-rich, microenvironmental factors such as extreme desiccation, local microbial fermentation, or anthropogenic interference can abruptly elevate Na^+^ to harmful levels in specific niches. Under such fluctuating conditions, maintaining a Na^+^-specific aversion pathway that can be selectively attenuated during nutrient deficit is likely favored by natural selection.

Salt taste constitutes an ancient sensory modality, constrained by the dual physiological imperative across Metazoa to secure sodium while preventing hyperosmotic toxicity ^[3]^.The IR11a-dependent receptor complex in *Drosophila* displays classical biophysical hallmarks of sodium sensing, such as amiloride sensitivity and Li^+^ permeability, which parallel the functional properties of mammalian low salt receptor ENaC ^[7]^. Although ionotropic receptors represent an invertebrate-expanded lineage derived from ionotropic glutamate receptors (iGluRs) that lacks direct vertebrate sequence orthologs, core co-receptors such as IR76b retain conserved sequence motifs in their pore-lining TM3 domains that are essential for Na^+^ permeation^[23]^. Importantly, whereas mammalian ENaCs drive low-salt attraction, the IR11a-dependent complex mediates high-salt aversion. This distinction illustrates a modular evolutionary logic. Despite primary sequence divergence, selective pressures may converge on similar three-dimensional pore architecture to recognize Na^+^, while the ultimate behavioral output of attraction or aversion is dictated by the cellular identity of the expressing gustatory cells. Given that the identity of mammalian high-salt receptors remains elusive, these findings suggest that vertebrate taste systems may similarly utilize sequentially distinct yet structurally and functionally analogous receptors to drive high-salt aversion.

### Limitations of the study

Our study presents several limitations that warrant future investigation. First, although IR11a is required for high-salt responses in a subset of bitter GRNs, its expression is not detected across all bitter GRNs, leaving open the question of whether IR11a-negative bitter neurons also contribute to salt sensation. Additionally, while our RNAi screen identified a mild phenotype for IR10a, its precise site of action and relevance to salt detection remain to be mapped. Second, residual NaCl avoidance persists in *IR7c;IR11a* double mutants, indicating that additional sensors contribute to high-salt aversion. Candidates include pharyngeal IR60b- and GR2a-expressing neurons, which have been implicated in salt avoidance ^[15, 34–36]^; however, the receptor composition of these pathways remains incompletely understood, and whether they undergo state-dependent modulation has not been tested. Third, the molecular mechanism underlying the state-dependent modulation of IR11a remains unresolved. Because qPCR detected no transcriptional changes and the lack of a functional antibody precluded examination at the protein level, post-translational modifications or receptor trafficking remain plausible hypotheses. Fourth, although heterologous co-expression of IR11a and IR25a in S2 cells elicited Na^+^-selective Ca^2+^ responses, whole-cell patch-clamp recordings could not be obtained because high-salt solutions disrupted osmotic balance and prevented gigaseal formation, leaving the channel’s intrinsic biophysical properties uncharacterized. Finally, high-resolution structural determination of the IR11a-containing complex and subsequent 3D motif-based screening may offer a powerful strategy to identify structurally or functionally analogous high-salt receptors in mammals.

## Materials and methods

### Flies husbandry and strains

Flies were reared on standard food at 25℃ and 70% humidity with 12 hr light/dark cycles. The standard food comprises 72.53 g of agar, 520 g of corn meal, 1100 g of malt extract, 275 g of yeast, 31.27 g of propionic acid, and 14.07 g of tegosept in 10L of water. Key Resouces Table provides the full list of source reagents for standard food and fly strains used in this study.

### Molecular biology and transgenic flies

#### pUAST-IR11a

Genomic DNA sequence was used to generate *UAS-IR11a* transgenic flies. DNA was extracted from *w^1118^* flies, and the genomic sequences of *IR11a* were PCR amplified and cloned into the pUAST vector (See Key Resouces Table for primers). The sequences were confirmed by sequencing before injection.

#### *IR11a-*gRNA, *IR7c-*gRNA and *poxn*-gRNA

The guide RNAs were designed from the website: www.e-crisp.org/E-CRISP/ ^[37]^. Two gRNAs of each gene were chosen and directly synthesized into the pCFD6 vector (See Key Resouces Table for primers).

All plasmids were injected into embryos of our control *w^1118^* line by Fungene Biotechnology Co., Ltd. to obtain the transgenic flies. Transgenic gRNA flies were crossed with flies that express CAS9 in germ cells to obtain candidate mutants ^[38]^.

The crossing scheme to generate the *IR11a* mutant is as:

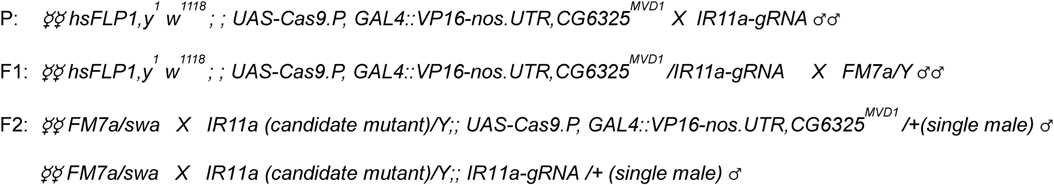

The crossing scheme to generate the *IR7c* mutant is as:

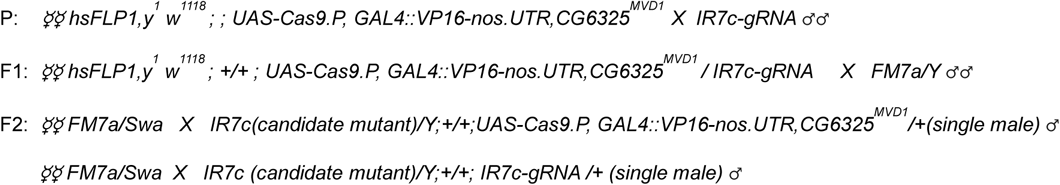

The crossing scheme to generate the *poxn* mutant is as:

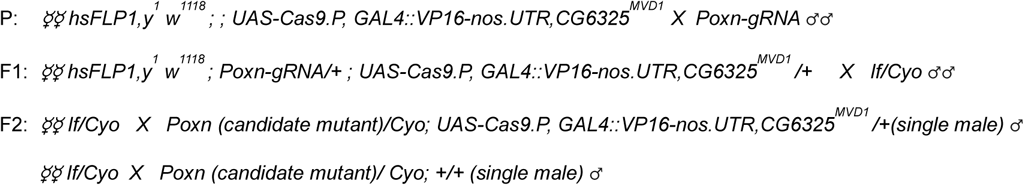

Candidate mutants were sequenced. *IR11a^4^* was identified as having a four-nucleotide deletion (TTCC at positions 173-176 of exon 2), and *IR7c^3^* was identified as having a 23-nucleotide deletion (TATCTGGTTCCTGGACAGTTTGC at positions 273 - 301 of exon 1), and *Poxn^84^* possessed a single-nucleotide deletion (C at position 18 of exon 4), all causing frameshift mutations. The mutant lines were backcrossed to *w^1118^* for six generations before calcium imaging and behavioral analyses.

### Behavioral Analysis

#### Two-choice feeding assays

two-choice feeding assays were performed as described previously ^[39]^, with minor modifications. Flies aged 8-10 days were used for the assays. Newly eclosed flies were maintained on standard food except when specified otherwise. For salt-deprived and salt-fed flies, newly eclosed flies were maintained on standard food for 2-4 days, then transferred to either salt-deficient food containing 0.7% agar and 563 mM sucrose, or salt-excess food containing 0.7% agar, 563 mM sucrose, and 50 mM NaCl, for 5 days. The 563 mM sucrose was used because it provides equivalent energy to our standard food. Each vial contained around 30 flies, starved for 18 to 20 hours in vials containing wet filter paper before the assays. Petri dishes (60 mm x 60 mm) were prepared immediately before each assay. Each dish was filled with 0.5% agarose mixed with tastants (including or excluding sucrose as specified), with one half containing blue dye (brilliant blue FCF at 0.125 mg/ml) and the other half containing red dye (sulforhodamine B at 0.2 mg/ml). Blue and red dyes were interchanged to eliminate bias due to dye color. Given that a higher proportion of flies fed on a mixture of tastant and sucrose compared to the tastant alone except for NaCl, we used a tastant-sucrose mix versus sucrose alone to assess the feeding preference or avoidance for tastants other than NaCl. The flies were chilled on ice and subsequently transferred to the dishes, where they were allowed to feed for 2 hours in dark conditions before being frozen for counting. Abdominal colors of flies in each dish were recorded under a stereomicroscope to assess feeding preferences. The abdomen of fed flies could exhibit blue, red, purple, or no coloration (indicating no feeding). The preference index (PI) was calculated using the formula: PI = [(number of flies with color 1) - (number of flies with color 2)] / (total number of fed flies). Data sets were discarded if the number of unfed flies exceeded half of the total fly count.

#### MAFE assay

Manual feeding (MAFE) assays were performed as described previously ^[31]^, with minor modifications. Flies used for MAFE assays received the same treatment as in the two-choice feeding assays. Flies were anesthetized on ice and affixed dorsally to glass slides using double-sided tape, with 20-30 flies per slide. The slides were then placed in an incubator at 25°C with 70% relative humidity for 1 hour to allow recovery. During the MAFE assay, each fly was first presented with water until it ceased drinking, followed by 250 mM NaCl solution until it stopped consuming the solution. This procedure was repeated for 3-4 rounds. Subsequently, 20 mM sucrose was applied using the same protocol for 3-4 rounds. If a fly did not consume 250 mM NaCl but drank 20 mM sucrose, it was scored as not feeding on 250 mM NaCl. Two datasets were collected: the proportion of flies that fed on 250 mM NaCl, and for those that did feed, the feeding duration was calculated. Both datasets were used in the Fig. 6H-J.

### Screening candidates for the high salt receptor

*UAS-RNAi* lines targeting candidate genes were crossed with the pan-neuronal driver *nSyb-GAL4*, in combination with the temperature-sensitive GAL4 inhibitor, *tubulin-GAL80^ts^* (*nSyb-GAL4, Tub-GAL80^ts^*). Flies were raised and maintained at 21°C to allow GAL80 to inhibit GAL4 activity, then shifted to 31°C for two days to restrict GAL80 activity before undergoing two-choice feeding assays. For the control, a subset of flies was raised and maintained at 21°C and was not shifted to 31°C before the assays. All two-choice feeding assays were conducted at 25°C. The two-choice assays presented flies with a choice between 400 mM NaCl + 5 mM sucrose and 1 mM sucrose to screen for potential high-salt receptor candidates. Flies with the same genotypes treated at 31°C that showed significantly decreased avoidance of high salt compared to those at 21°C were considered to carry candidate genes encoding high salt receptors.

### Immunohistochemistry

Fly dissection and immunostaining were conducted as described previously ^[40]^. Flies aged 6-10 days were used for dissection. Flies were anesthetized with CO_2_ until immobile, and the labellum and legs were excised with a razor blade. The legs were cut between the third and fourth segments of the tarsi. The preparations were placed in 4% paraformaldehyde immediately after dissection, and the collected preparations were briefly centrifuged at 1000 rpm for 30 seconds to ensure complete immersion of the tissues in the fixation solution. The preparations were fixed on ice for 1 hour, then washed three times with PBST (1x PBS + 0.3% Triton X-100) at room temperature for 15 minutes each. For antibody staining, preparations were incubated in PBST + 5% horse serum solution for 2 hours at room temperature. Following this, they were rinsed twice and subsequently incubated with primary antibodies overnight at 4°C. Afterward, the preparations were washed three times with PBST for 15 minutes each. Subsequently, the samples were incubated in PBST + 5% horse serum for 30 minutes at room temperature, rinsed twice again, and then incubated with secondary antibodies for 3 hours at room temperature. Secondary antibodies were removed and preparations were washed three times with PBST for 15 minutes each. For mounting, preparations were placed on slides with a drop of antifade mounting medium, then covered with a coverslip and sealed with nail polish. Antibodies were diluted in PBST. Primary antibodies used in this study: chicken anti-GFP (1:2000), rabbit anti-RFP (1:2000), mouse anti-HA (1:500), mouse anti-nc82 (1:5000), and rabbit anti-IR25a (1:200). The IR25a antibodies were produced utilizing the same synthetic peptides described in Benton et al ^[28]^ as the antigen: SKAALRPRFNQYPATFKPRF and DVAEANAERSNAADHPGKLVDGV. Secondary antibodies: goat anti-chicken Alexa 488 (1:2000), goat anti-rabbit Alexa 488 (1:200), goat anti-mouse Alexa 568 (1:2000), goat anti-rabbit Alexa 568 (1:2000), and goat anti-mouse Alexa 647 (1:2000). Images were taken using a Zeiss LSM 800 confocal microscope with a 20X or 40X objective.

### Calcium imaging

*UAS-GCaMP6m* was expressed under the control of different GAL4 drivers as specified in the experiments. Female flies aged 7-10 days were used for calcium imaging.

#### Calcium imaging in the SEZ (subesophageal zone)

calcium imaging in the SEZ was conducted as described by Marella et al ^[26]^. Flies were anesthetized on ice. The back of the neck and thorax were fixed in a custom chamber using glue. The proboscis was extended and immobilized with a needle. Antennae were removed, and a small window was cut in the cuticle around the SEZ to expose the brain. The exposed SEZ region was bathed in AHL solution (108 mM NaCl, 8.2 mM MgCl_2_, 4 mM NaHCO_3_, 1 mM NaH_2_PO_4_, 2 mM CaCl_2_, 5 mM KCl, 5 mM HEPES, pH 7.5). The sample was examined under a Nikon Ni-U upright microscope with a 40X water immersion objective. For labellum stimulation, a tastant solution in an injector mounted on a micromanipulator was delivered to the labellum for 5 seconds. Fluorescence recording of GCaMP started 5 seconds before stimulation and continued for 10 seconds after stimulation. The labellum was rinsed with water at the end of each recording and allowed to recover for 1 minute before the next recording. Each preparation was stimulated with different tastants in ascending concentrations.

#### Calcium imaging of single GRNs in the legs

calcium imaging in the legs was performed as described previously ^[41]^. A piece of double-sided tape was affixed to a glass-bottom petri dish (35 x 35 mm). The legs were cut between the tibia and femur, with the cut end coated with grease and affixed to the tape, leaving the 4^th^ and 5^th^ tarsal segments hanging off the tape. A drop of melted agarose (50-60°C) was placed on the tape to secure the preparation. Imaging was conducted using a Nikon Ni-U upright microscope with a 40X water immersion objective. The fluorescence of the cell body was monitored, using the adjacent area as an internal control. Tastants were applied with a pipette, with recording starting 10 seconds before tastant application and lasting for 2-3 minutes.

The fluorescence changes in SEZ and legs were calculated using the same formula: ΔF/F (%) = (F - F0) / F0, where F is the fluorescence value, and F0 is the mean baseline value (the average fluorescence of the 5 seconds before stimulation). The maximum ΔF/F (%) value was used for statistical analysis.

### S2 cell culture, transfection, and calcium imaging

S2 cell experiments were adapted from Asefa et al with modifications ^[15]^*. Drosophila* S2 cells were cultured in Sf-900^TM^ II SFM medium supplemented with 10% fetal bovine serum (FBS) and 0.5% penicillin/streptomycin at 25°C without CO_2_ supplementation. Cells were passaged every 3 days at a 1:2 to 1:3 dilution ratio. The genomic DNA sequence of IR11a and the cDNA sequences of IR25a, IR76b, and GCaMP6m were cloned into the pAc5.1 vector; pAc5.1-GFP was provided by Dr. Guoqiang Zhang. Calcium imaging was performed using a Nikon Ni-U upright microscope equipped with a 10× objective. Immediately before imaging, the culture medium containing transfection reagent was replaced with an isotonic bath buffer (10 mM HEPES, 10 mM glucose, 250 mM NMDG-Cl; pH 7.4; 520 mOsm). Approximately 2 minutes after buffer exchange, the bath buffer was gently removed, leaving only a minimal volume sufficient to immerse the cells. Baseline fluorescence (F_0_) was recorded for 10 seconds. Stimulation was then initiated by adding ligand solutions containing x mM salt (NaCl, KCl, or LiCl), 10 mM HEPES, 10 mM glucose, and y mM NMDG-Cl (pH 7.4); the salt concentration (x) and NMDG-Cl concentration (y) were adjusted to achieve 520 mOsm. Fluorescence (F) was monitored continuously for 120 seconds. Fluorescence changes were calculated as: ΔF/F (%) = [(F – F_0_) / F_0_] × 100. Peak ΔF/F responses, following background subtraction (using adjacent non-fluorescent regions of interest), were quantified for statistical analysis.

### Quantification of IR25a expression in *IR11a-expressing* GRNs

To quantify IR25a expression in *IR11a-expressing* GRNs, we dissected the fifth tarsal segments of *GR33a^Gal4^*>*UAS-mCD8-RFP* flies maintained under salt-deprived or salt-fed conditions. Tissues were fixed and incubated overnight at 4 °C with rabbit anti-IR25a (1:200) and Rat anti-mCD8 (1:2000), followed by goat anti-rabbit Alexa 488 and goat anti-rat Alexa 568 (1:2000). All samples were processed in parallel and imaged on a Zeiss LSM 800 confocal microscope with identical laser power and acquisition settings. Maximum-intensity projections were generated in Zeiss Zen, and mean fluorescence intensities were measured in the green (488 nm) and red (568 nm) channels. A signal-free region served as background. Relative IR25a signal was calculated as (F_green_ – B_green_)/(F_red_ – B_red_) for each leg segment.

### Quantitative RT-PCR

Quantitative RT-PCR was performed to determine ionotropic receptor (IR) gene expression under salt-deprived and salt-fed conditions. Labella were dissected from female flies in triplicate biological replicates. Total RNA was extracted with the Quick-RNA Microprep Kit (Beijing Tianmo Biotech), and 500 ng RNA was reverse-transcribed using HiScript II Q RT SuperMix (+gDNA wiper) (Vazyme). Reactions (15 µl) contained 2× M5 HiPer SYBR Premix EsTaq (7.5 µl), 1:10-diluted cDNA (4 µl), 10 µM gene-specific primers (0.3 µl each), ROX reference dye (0.3 µl), and nuclease-free water (2.6 µl). Cycling conditions were 95 °C for 5 min, followed by 40 cycles of 95 °C for 15 s and 60 °C for 30 s. Relative expression was calculated by the 2^−ΔΔCT^ method ^[42]^ with rp49 as the endogenous control. Primer sequences are provided in Key Resouces Table.

### Quntification and statistical analysis

Normal distribution was evaluated, and nonparametric tests were used for non-normal data. The Mann-Whitney test was used to compare two groups, while the Kruskal-Wallis test with Dunn’s multiple comparisons was used for multiple group comparisons. Detailed quantification and statistical analysis methods are listed in Data S1-S9.

### Key Resouces Table

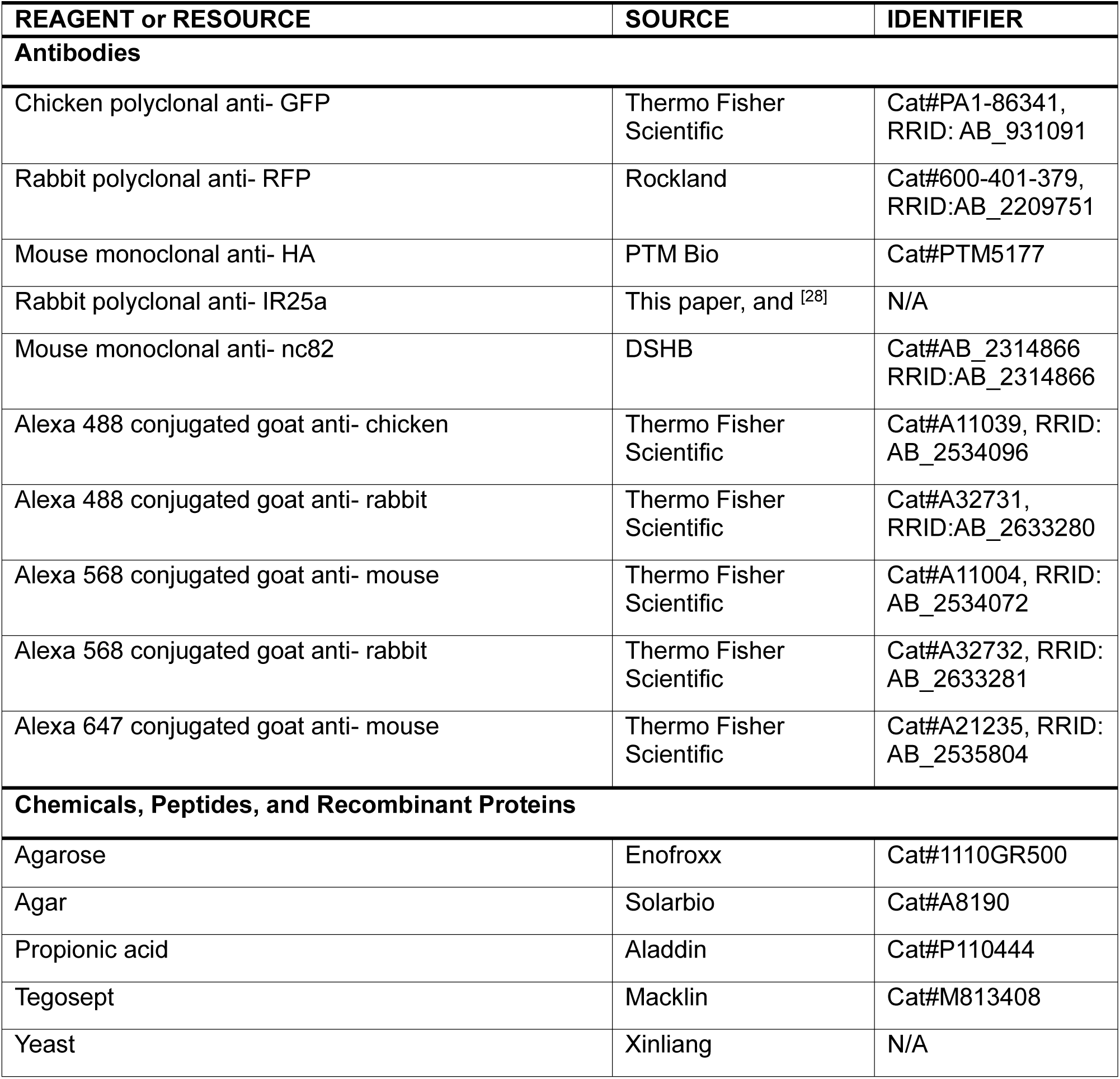
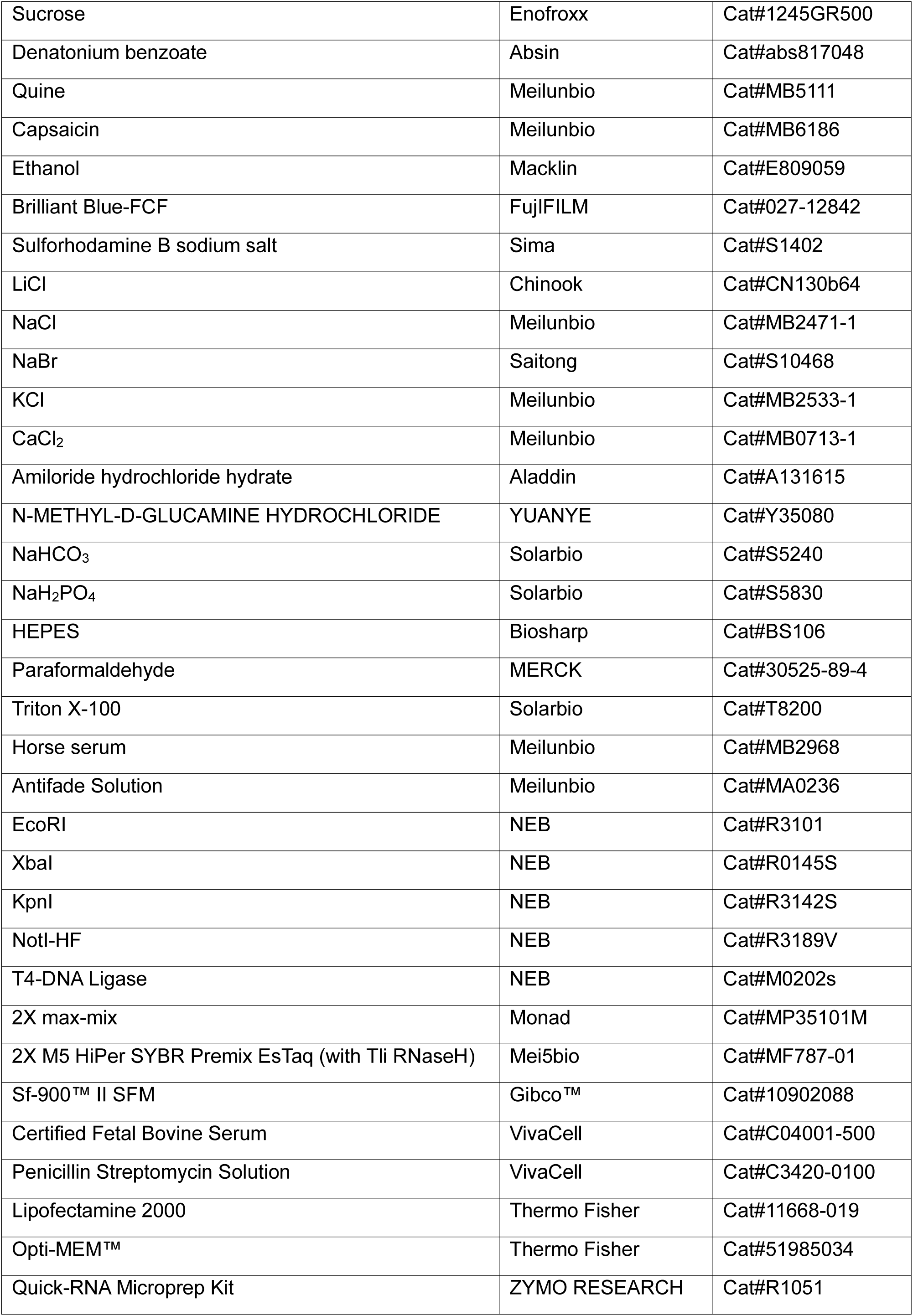
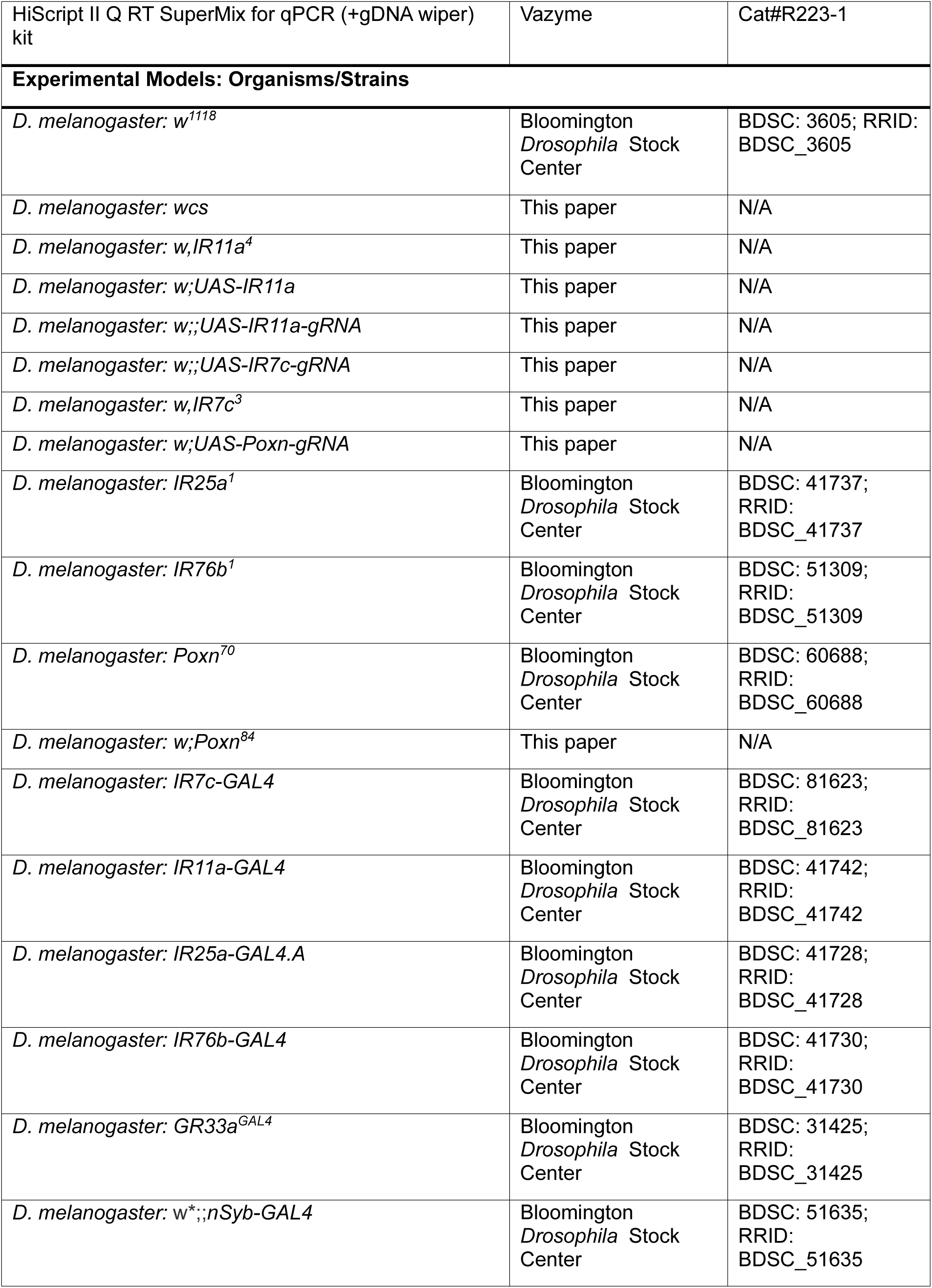
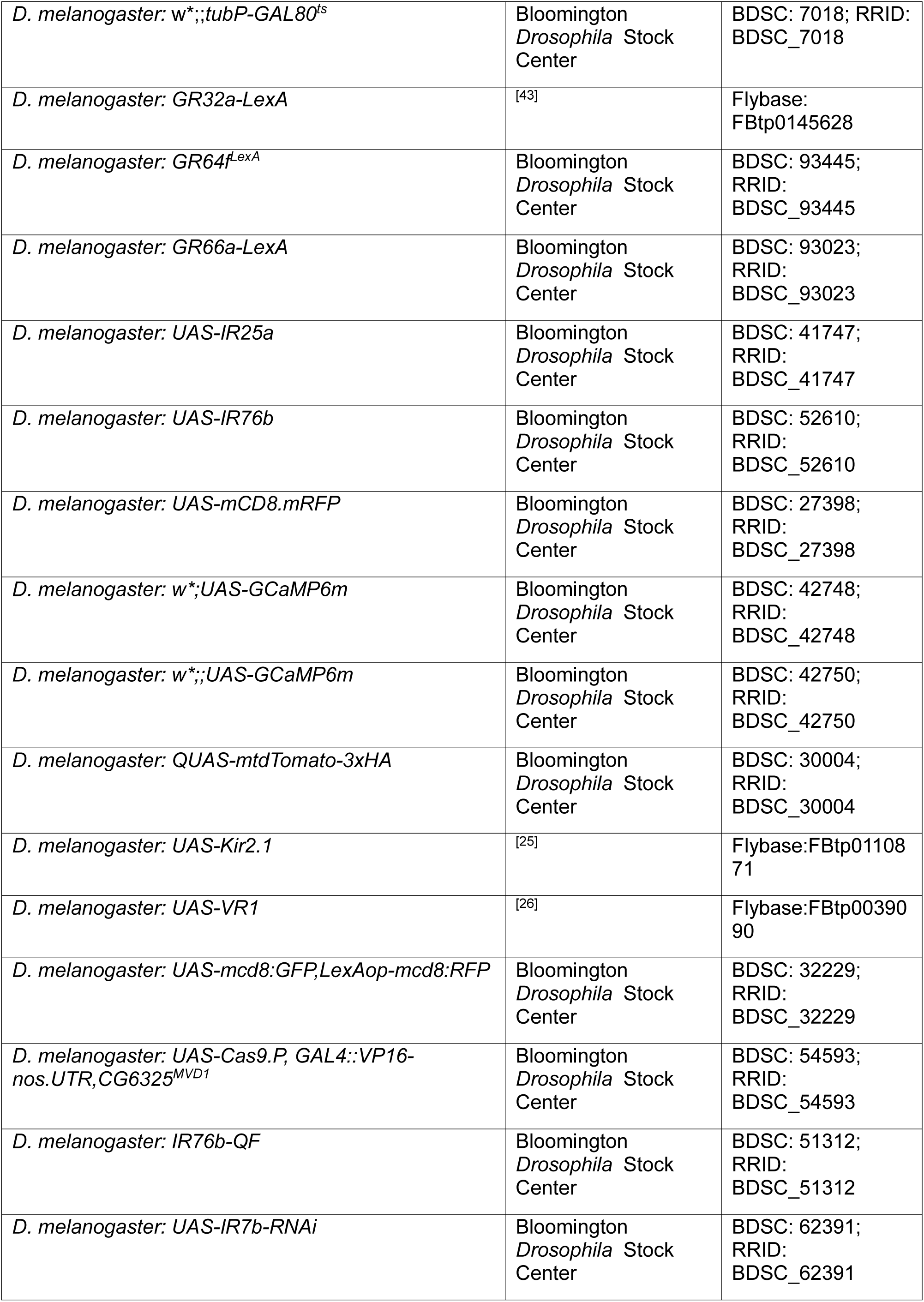
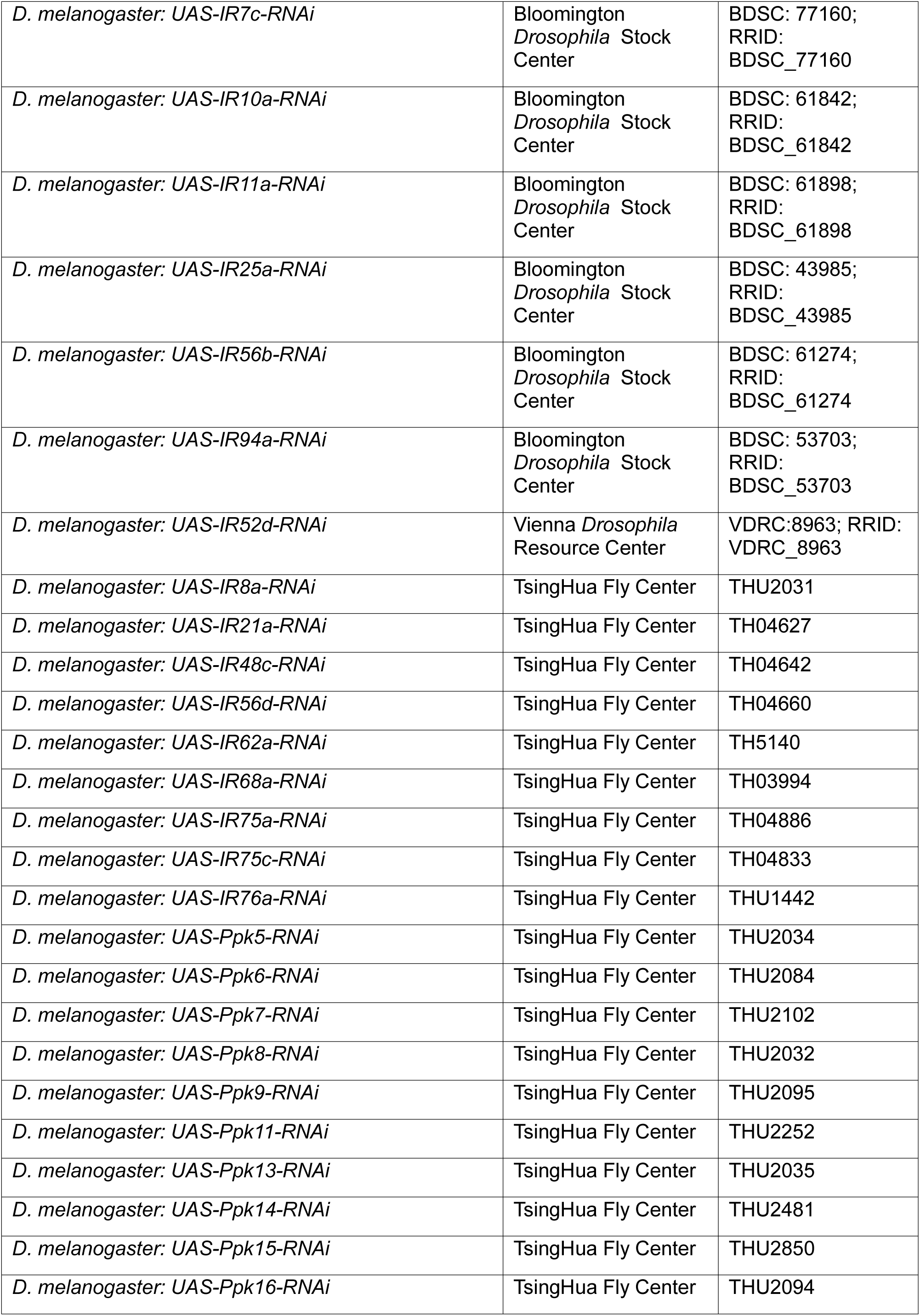
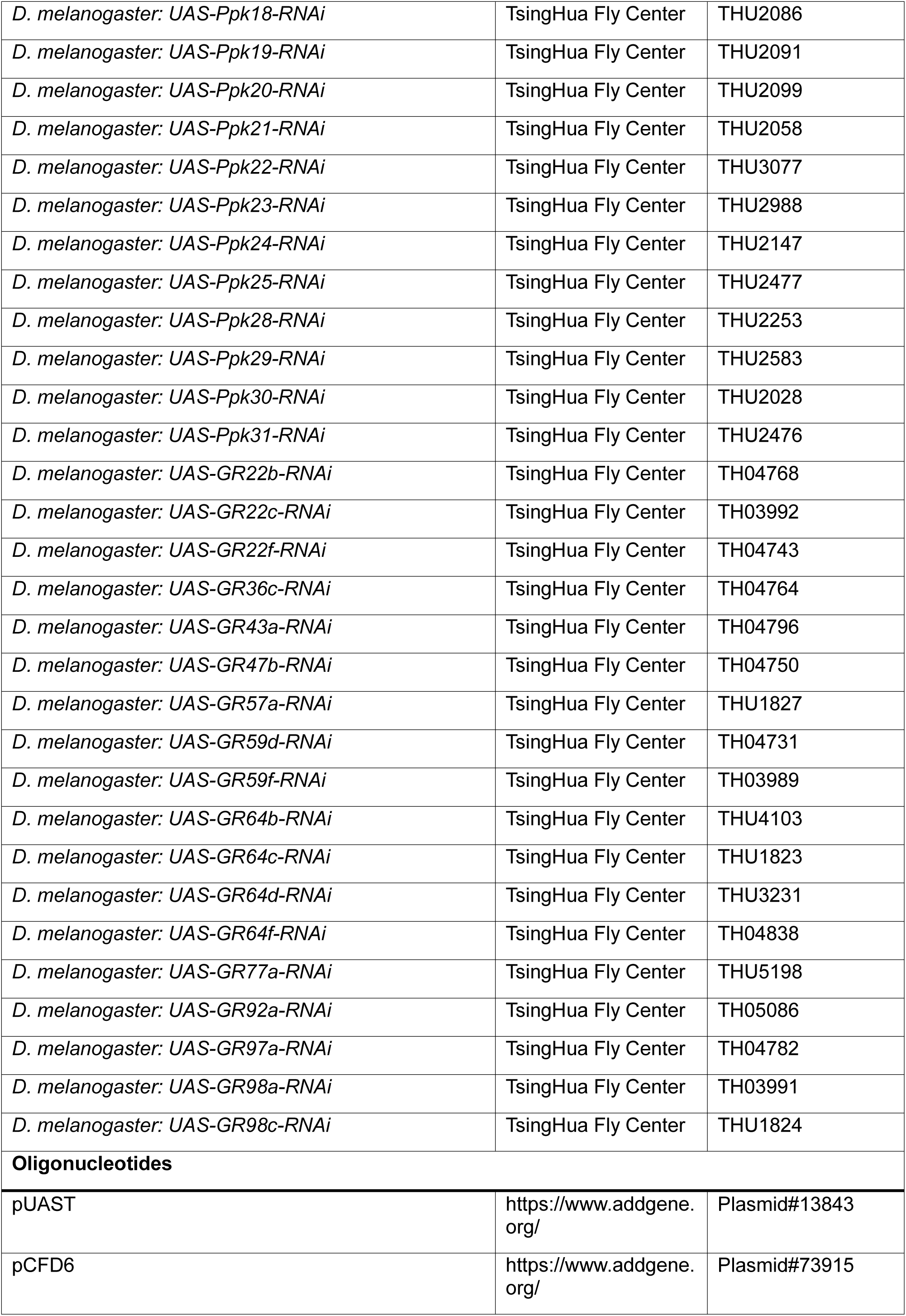
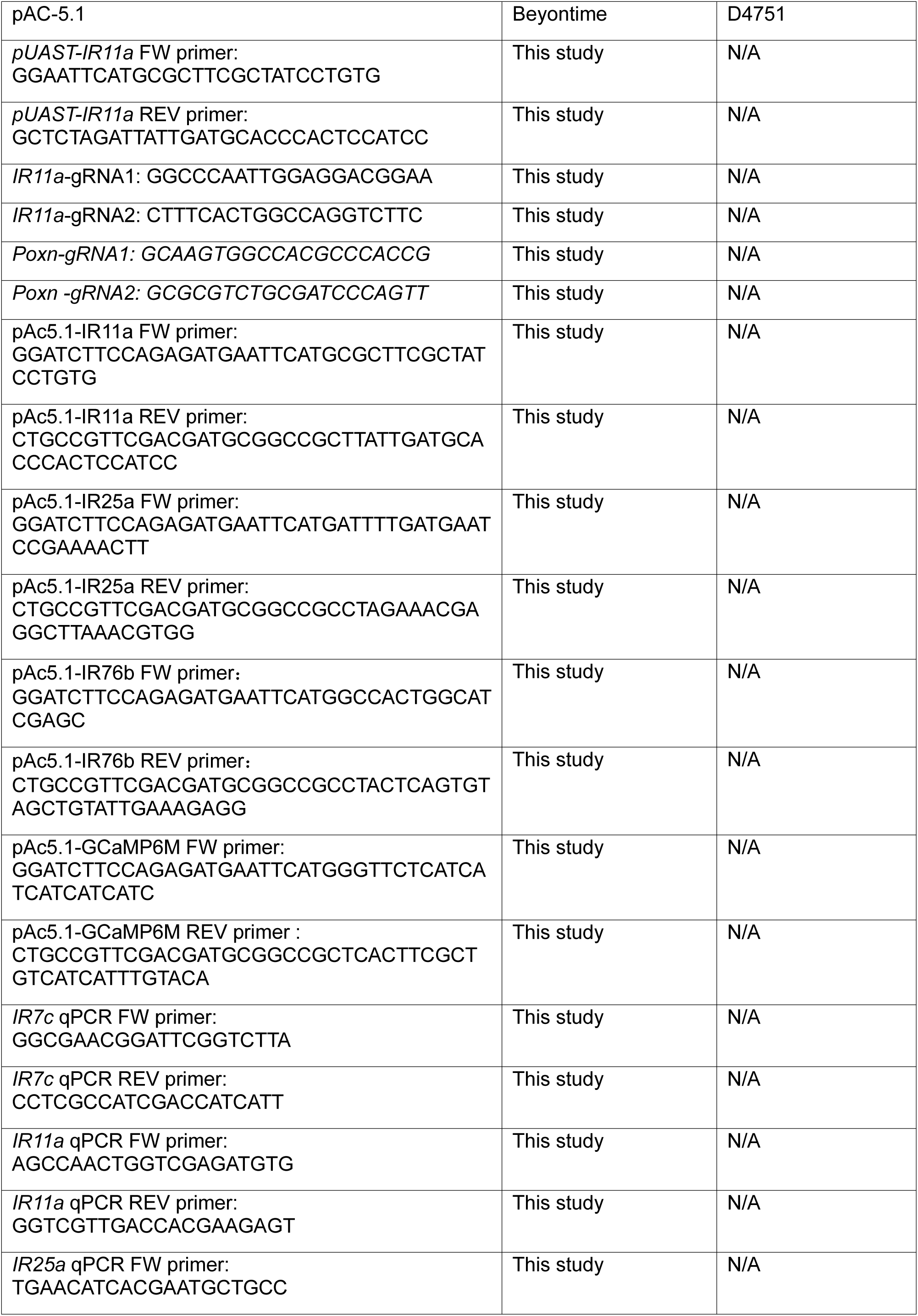
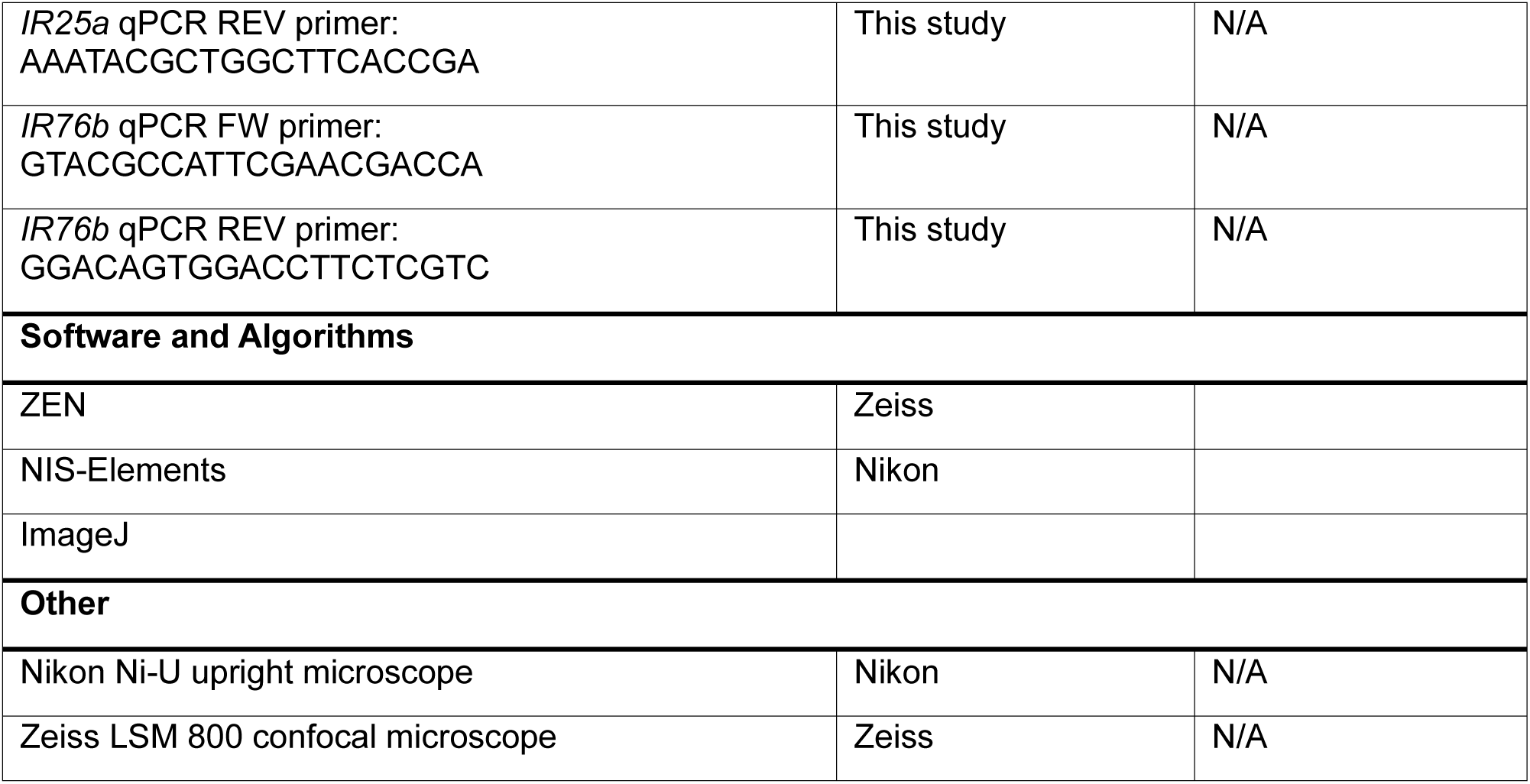

## Supporting information

Supplemental dataset for statistics and genotype detail

## Acknowledgments

We thank Drs. Hubert Amrein, Wei Zhang, Zhihua Liu, Shan Jin, Hui Xiao, Guoqiang Zhang, the Bloomington *Drosophila* Stock Center, the Tsinghua Fly Center, and VDRC for fly strains and plasmids, and the Animal Center of Yunnan University for antibodies. We thank Yan Pan for assistance with the MAFE assay and Drs. Wei Zhang, Zhihua Liu, Bin Qi, and Qingfeng Chen for critical comments on the manuscript.

## Funding

This work was supported by grants from the National Natural Science Foundation of China (https://www.nsfc.gov.cn/, No. to Y.C), Applied Basic Research Foundation of Yunnan Province (https://kjgl.kjt.yn.gov.cn/egrantweb/, No. 2019FY003018 to Y.C.), Research Startup Funding of Yunnan University (https://www.ynu.edu.cn/, to Y.C.). The funders had no role in study design, data collection and analysis, decision to publish, or preparation of the manuscript.

## Authors contribution

G.Q. performed the screen, molecular biology, immunostaining, leg calcium imaging, and two-choice feeding assays, B.W. performed SEZ calcium imaging and two-choice feeding assays, G.Q., B.W., and Y.C. analyzed data, Y.C. conceived and supervised the project, designed experiments, and wrote the paper.

## Declaration of interests

The authors declare no competing interests.

---

**Fig. S1.**
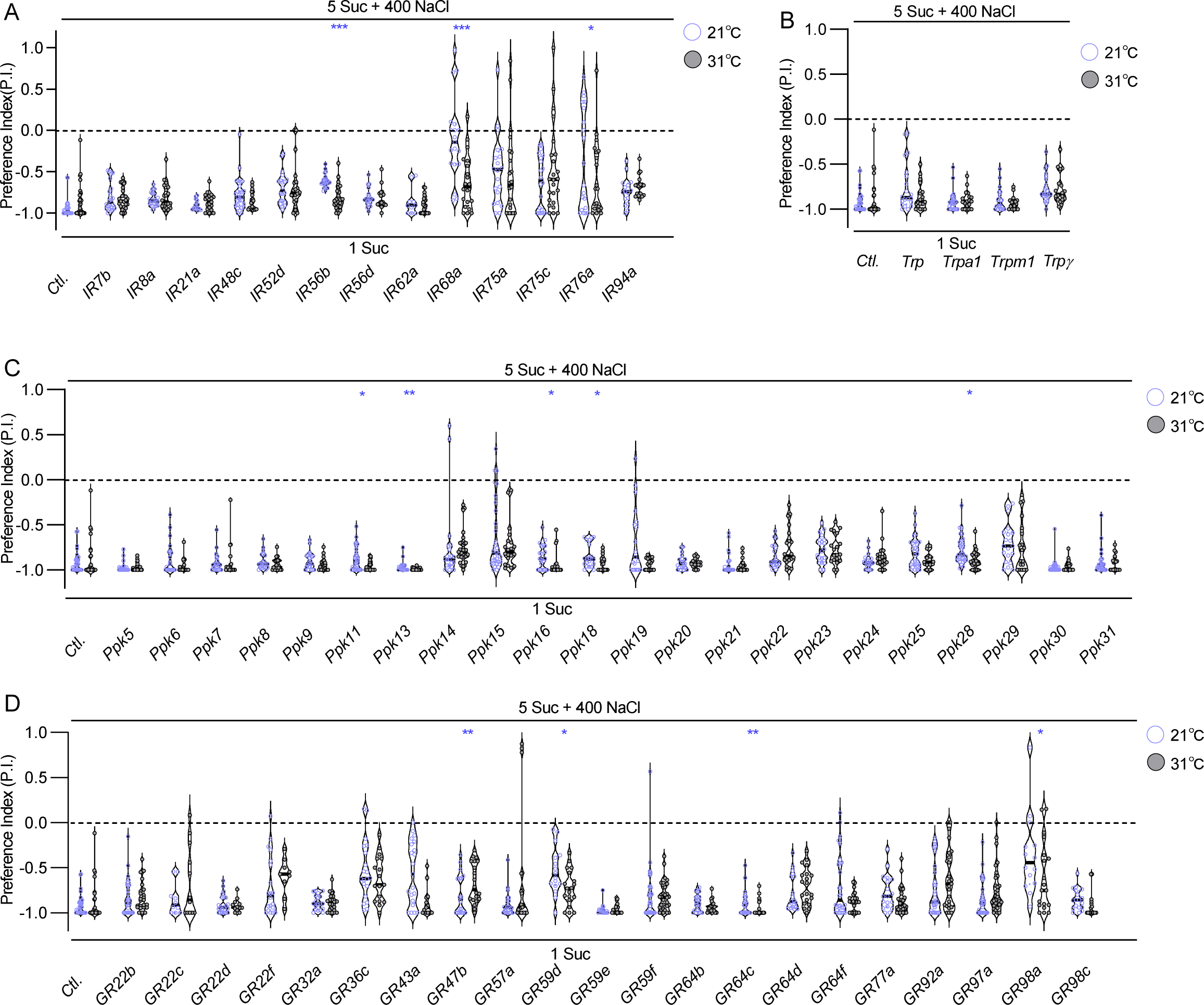
A pan-neuronal RNAi screen for receptor genes mediating high-salt avoidance (Related to Fig. 1). (A - D) Avoidance of 400 mM NaCl following pan-neuronal knockdown (*nSyb-GAL4/+; Tub-GAL80^ts^/+* > *UAS-RNAi*) of target genes under restrictive (21°C) and permissive (31°C) conditions. Blue asterisks indicate significant increases in avoidance compared to controls. Genes whose knockdown increased avoidance were excluded as candidates, as high-salt receptor knockdown is expected to reduce avoidance. Concentrations are in mM. Data are mean ± SEM (n = 10 - 33). Mann-Whitney test; *p<0.05, **p<0.01, ***p<0.001. Genotypes and statistical details are provided in Data S7.

**Fig. S2.**
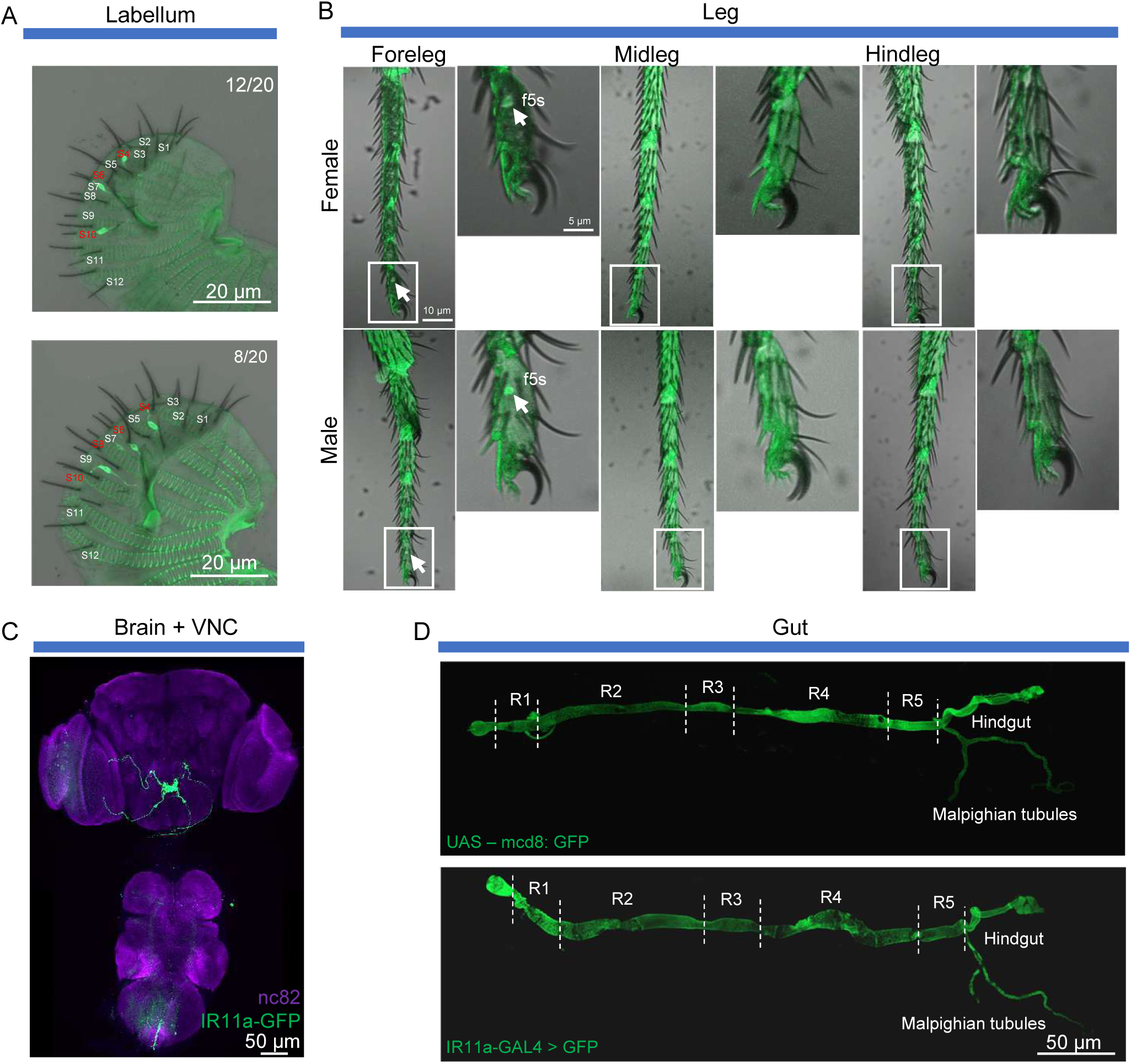
Expression analysis of IR11a-GAL4 (Related to Fig. 2). **(A)** *IR11a-GAL4* labels S4, S6, and S10 sensilla consistently in the labellum, with additional expression in S8 sensilla in 40% of preparations. **(B)** *IR11a-GAL4* is exclusively expressed in the f5s sensillum of the forelegs; arrowheads indicate cell bodies in the 5s sensillum. **(C)** Axonal projections of *IR11a-GAL4*-expressing neurons in the brain and ventral nerve cord. **(D)** No expression of IR11a is observed in the gut or Malpighian tubules. Genotypes details are provided in Data S8.

**Fig. S3.**
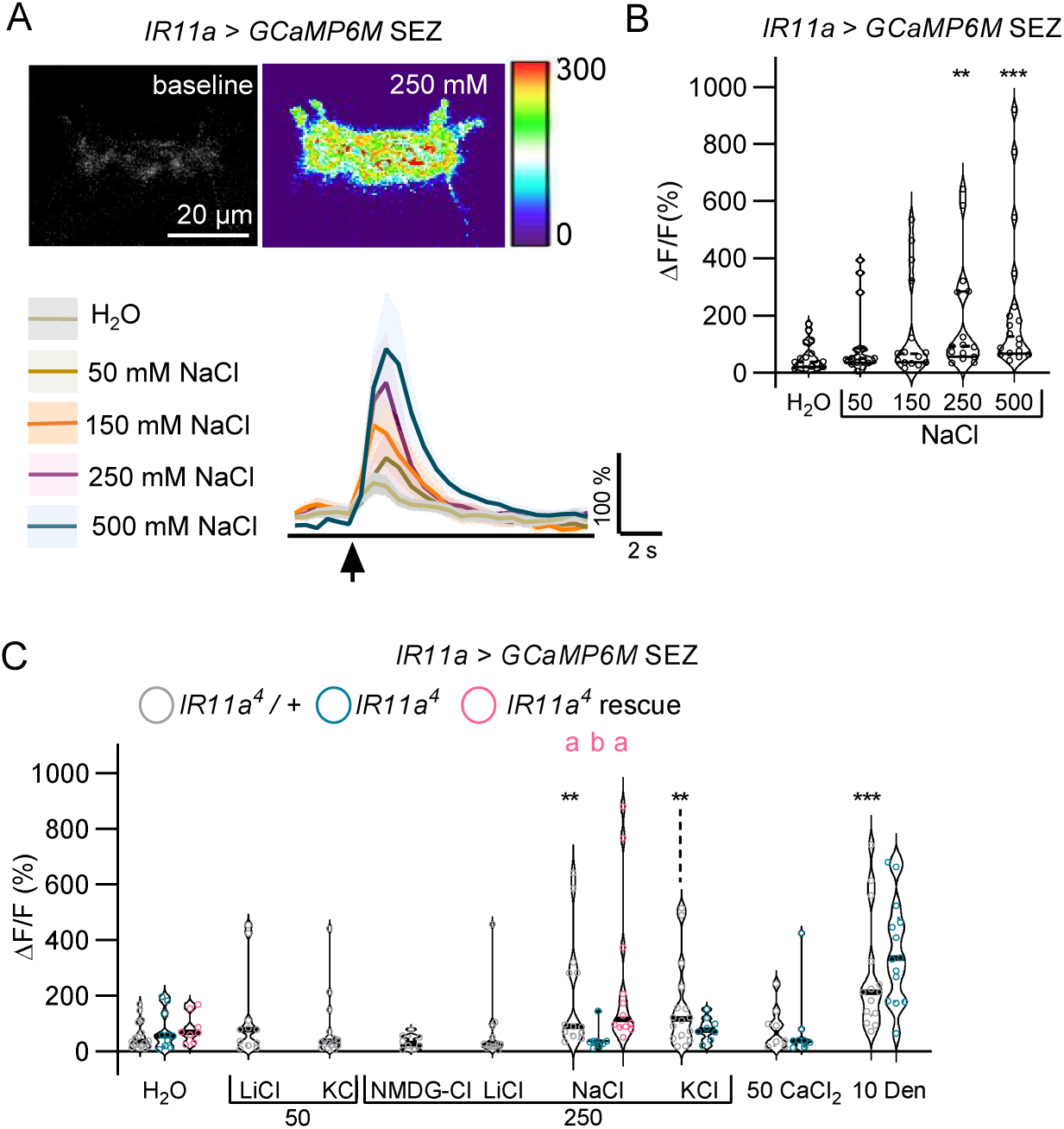
IR11a is selectively required for high Na^+^ responses in labellum GRNs (Related to Fig. 3). **(A - B)** Calcium responses at axonal terminals in the SEZ of *IR11a*-expressing labellum GRNs to 50 - 500 mM NaCl. **(C)** Calcium responses at axonal terminals in the SEZ of *IR11a*-expressing labellum GRNs to NaCl, LiCl, NMDG-Cl, NaBr, KCl, CaCl_2_, and denatonium in *IR11a* controls, mutants, and rescue flies. Concentrations are in mM. Data are mean ± SEM (n = 8-16). Mann-Whitney test for **(B, C)** comparing tastant responses with H_2_O; *p<0.05, **p<0.01, ***p<0.001. Kruskal-Wallis with Dunn’s multiple comparisons test for **(C)** comparing genotypes stimulated with the same tastant; different letters indicate p<0.05. Genotypes and statistical details are provided in Data S9.

**Fig. S4.**
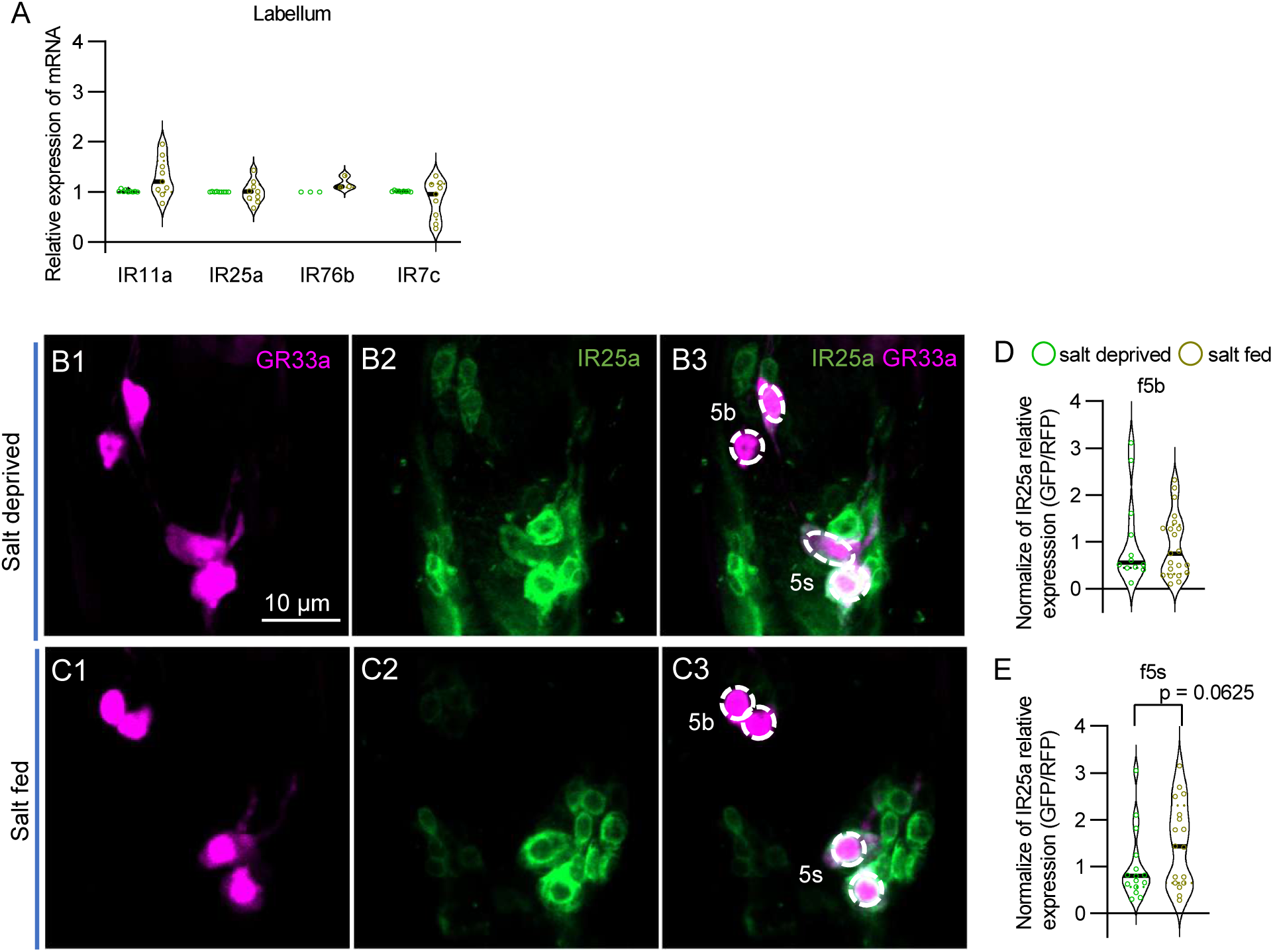
IR expression under salt-deprived and salt-fed conditions (Related to Fig. 6). **(A)** qPCR analysis of *IR25a*, *IR76b*, *IR11a*, and *IR7c* transcript levels in salt-deprived and salt-fed flies. **(B - E)** Anti-IR25a immunostaining of *IR11a*-expressing tarsal GRNs (labeled by *GR33a^GAL4^*) under salt-deprived and salt-fed conditions. Data are mean ± SEM (n = 3 - 21). Mann-Whitney test for (A, D - E).

## Notes

### Competing Interest Statement

The authors have declared no competing interest.

### Summary of Updates

Reorganize the text to better align it with the data.

## References

[1] Collaborators GBDD. Health effects of dietary risks in 195 countries, 1990-2017: A systematic analysis for the global burden of disease study 2017. Lancet, 2019, 393: 1958–1972

[2] Liman Emily R, Zhang Yali V, Montell C. Peripheral coding of taste. Neuron, 2014, 81: 984–1000

[3] Taruno A, Gordon MD. Molecular and cellular mechanisms of salt taste. Annual Review of Physiology, 2023, 85: 25–45

[4] Zhang Y, Pool A-H, Wang T, et al. Parallel neural pathways control sodium consumption and taste valence. Cell, 2023,

[5] Shigemura N, Iwata S, Yasumatsu K, et al. Angiotensin ii modulates salty and sweet taste sensitivities. J Neurosci, 2013, 33: 6267–6277

[6] McDowell SAT, Li J, Gordon MD. Modulation of attractive salt taste in drosophila. iScience, 2026, 29: 115128

[7] Chandrashekar J, Kuhn C, Oka Y, et al. The cells and peripheral representation of sodium taste in mice. Nature, 2010, 464: 297–301

[8] Lindemann B. Receptors and transduction in taste. Nature, 2001, 413: 219–225

[9] Oka Y, Butnaru M, von Buchholtz L, et al. High salt recruits aversive taste pathways. Nature, 2013, 494: 472–475

[10] Kasahara Y, Narukawa M, Ishimaru Y, et al. Tmc4 is a novel chloride channel involved in high-concentration salt taste sensation. The Journal of Physiological Sciences, 2021, 71:

[11] Lyall V, Heck GL, Vinnikova AK, et al. The mammalian amiloride-insensitive non-specific salt taste receptor is a vanilloid receptor-1 variant. The Journal of Physiology, 2004, 558: 147–159

[12] Jaeger AH, Stanley M, Weiss ZF, et al. A complex peripheral code for salt taste in drosophila. eLife, 2018, 7:

[13] Montell C, Truman J. Drosophilasensory receptors—a set of molecular swiss army knives. Genetics, 2021, 217: 1–34

[14] Dweck HKM, Talross GJS, Luo Y, et al. Ir56b is an atypical ionotropic receptor that underlies appetitive salt response in drosophila. Curr Biol, 2022,

[15] Asefa WR, Woo JN, Kim SY, et al. Molecular and cellular basis of sodium sensing in drosophila labellum. iScience, 2024, 27: 110248

[16] Wang B, Qu G, Wang X, et al. The drosophila ir20a integrates l-arginine and salt signals via distinct subunit assemblies. 2026, 10.7554/eLife.112388.1

[17] McDowell SAT, Stanley M, Gordon MD. A molecular mechanism for high salt taste in drosophila. Curr Biol, 2022, 32: 3070–3081 e3075

[18] Freeman EG, Dahanukar A. Molecular neurobiology of drosophila taste. Current Opinion in Neurobiology, 2015, 34: 140–148

[19] Joseph RM, Carlson JR. Drosophila chemoreceptors: A molecular interface between the chemical world and the brain. Trends Genet, 2015, 31: 683–695

[20] Scott K. Gustatory processing in drosophila melanogaster. Annual Review of Entomology, 2018, 63: 15–30

[21] Sánchez-Alcañiz JA, Silbering AF, Croset V, et al. An expression atlas of variant ionotropic glutamate receptors identifies a molecular basis of carbonation sensing. Nature Communications, 2018, 9:

[22] Thistle R, Cameron P, Ghorayshi A, et al. Contact chemoreceptors mediate male-male repulsion and male-female attraction during drosophila courtship. Cell, 2012, 149: 1140–1151

[23] Zhang YLV, Ni JF, Montell C. The molecular basis for attractive salt-taste coding in. Science, 2013, 340: 1334–1338

[24] Miyamoto T, Slone J, Song X, et al. A fructose receptor functions as a nutrient sensor in the drosophila brain. Cell, 2012, 151: 1113–1125

[25] Baines RA, Uhler JP, Thompson A, et al. Altered electrical properties in drosophila neurons developing without synaptic transmission. J Neurosci, 2001, 21: 1523–1531

[26] Marella S, Fischler W, Kong P, et al. Imaging taste responses in the fly brain reveals a functional map of taste category and behavior. Neuron, 2006, 49: 285–295

[27] Moon SJ, Lee Y, Jiao Y, et al. A drosophila gustatory receptor essential for aversive taste and inhibiting male-to-male courtship. Current Biology, 2009, 19: 1623–1627

[28] Benton R, Vannice KS, Gomez-Diaz C, et al. Variant ionotropic glutamate receptors as chemosensory receptors in drosophila. Cell, 2009, 136: 149–162

[29] Ling F, Dahanukar A, Weiss LA, et al. The molecular and cellular basis of taste coding in the legs ofdrosophila. The Journal of Neuroscience, 2014, 34: 7148–7164

[30] Awasaki T, Kimura K-i. Pox-neuro is required for development of chemosensory bristles in drosophila. Journal of neurobiology, 1997, 32 7: 707–721

[31] Qi W, Yang Z, Lin Z, et al. A quantitative feeding assay in adult drosophila reveals rapid modulation of food ingestion by its nutritional value. Mol Brain, 2015, 8: 87

[32] Jin L, Kim CH, Seo JT, et al. Dietary salt induces taste desensitization via receptor internalization in drosophila in a sexually dimorphic manner. Molecules and Cells, 2025, 48:

[33] Kaushik S, Singh K, Kumar R, et al. Activity and state-dependent modulation of salt taste behavior via pharyngeal neurons in drosophila melanogaster. Neurosci Insights, 2026, 21: 26331055261440786

[34] Puri S, Sang J, Pandey P, et al. Insulin and leucokinin pathways coordinate adaptive salt appetite in drosophila. Proc Natl Acad Sci U S A, 2026, 123: e2530544123

[35] Sang J, Dhakal S, Shrestha B, et al. A single pair of pharyngeal neurons functions as a commander to reject high salt in drosophila melanogaster. eLife, 2023,

[36] Kim H, Jeong YT, Choi MS, et al. Involvement of a gr2a-expressing drosophila pharyngeal gustatory receptor neuron in regulation of aversion to high-salt foods. Molecules and Cells, 2017, 40: 331–338

[37] Heigwer F, Kerr G, Boutros M. E-crisp: Fast crispr target site identification. Nature Methods, 2014, 11: 122–123

[38] Port F, Chen H-M, Lee T, et al. Optimized crispr/cas tools for efficient germline and somatic genome engineering in drosophila. Proceedings of the National Academy of Sciences, 2014, 111:

[39] Chen Y, Amrein H. Enhancing perception of contaminated food through acid-mediated modulation of taste neuron responses. Current Biology, 2014, 24: 1969–1977

[40] Chen Y, Amrein H. Ionotropic receptors mediate drosophila oviposition preference through sour gustatory receptor neurons. Current Biology, 2017, 27: 2741–2750.e2744

[41] Miyamoto T, Chen Y, Slone J, et al. Identification of a drosophila glucose receptor using ca2+ imaging of single chemosensory neurons. PLoS ONE, 2013, 8:

[42] Livak KJ, Schmittgen TD. Analysis of relative gene expression data using real-time quantitative pcr and the 2(-delta delta c(t)) method. Methods, 2001, 25: 402–408

[43] Liu C, Zhang B, Zhang L, et al. A neural circuit encoding mating states tunes defensive behavior in drosophila. Nature Communications, 2020, 11: 3962

