## Supplemental dataset for statistics and genotype detail for "A Na^+^-selective receptor enables state-dependent plasticity in high-salt taste"

**Data S1-Genotypes and statistical details for Fig.1**

| Figures | Foods | Genotypes | Genotypes labeled in figure | N value | Compare groups | Statistical methods | P value summary | P value |
|---|---|---|---|---|---|---|---|---|
| Fig1-A | 5 mM Sucrose + 400 mM NaCl vs. 1 mM Sucrose | nSyb-GAL4,Tub-GAL80$^{ts}$ / + 21℃ | Ctl. | 23 | 21℃ vs. 31℃ | Mann-Whitney | | |
| | | nSyb-GAL4,Tub-GAL80$^{ts}$ / + 31℃ | | 24 | | | ns | 0.3636 |
| | | nSyb-GAL4,Tub-GAL80$^{ts}$ >IR25a-RNAi 21℃ | IR25a | 20 | | | | |
| | | nSyb-GAL4,Tub-GAL80$^{ts}$ >IR25a-RNAi 31℃ | | 10 | | | ** | 0.006 |
| | | nSyb-GAL4,Tub-GAL80ts>IR7c-RNAi 21℃ | IR7c | 22 | | | | |
| | | nSyb-GAL4,Tub-GAL80ts>IR7c-RNAi 31℃ | | 23 | | | *** | <0.001 |
| | | nSyb-GAL4,Tub-GAL80ts>IR11a-RNAi 21℃ | IR11a | 32 | | | | |
| | | nSyb-GAL4,Tub-GAL80ts>IR11a-RNAi 31℃ | | 29 | | | *** | <0.001 |
| | | nSyb-GAL4,Tub-GAL80ts>IR10a-RNAi 21℃ | IR10a | 22 | | | | |
| | | nSyb-GAL4,Tub-GAL80ts>IR10a-RNAi 31℃ | | 23 | | | * | 0.0418 |
| Fig1-C | 5 mM Sucrose + 400 mM NaCl vs. 1 mM Sucrose | IR11a$^{4}$/+ | IR11a$^{4}$/+ | 16 | IR11a$^{4}$/+ vs. IR11a$^{4}$ | Mann-Whitney | *** | <0.0001 |
| | | IR11a$^{4}$/IR11a$^{4}$ | IR11a$^{4}$ | 16 | | | | |
| Fig1-D | 50 mM NaCl vs. $H_2O$ | IR11a$^{4}$/+ | IR11a$^{4}$/+ | 31 | IR11a$^{4}$/+ vs. IR11a$^{4}$ | Mann-Whitney | * | 0.0364 |
| | | IR11a$^{4}$/IR11a$^{4}$ | IR11a$^{4}$ | 24 | | | | |
| | 250 mM NaCl vs. $H_2O$ | IR11a$^{4}$/+ | IR11a$^{4}$/+ | 26 | IR11a$^{4}$/+ vs. IR11a$^{4}$ | | *** | <0.001 |
| | | IR11a$^{4}$/IR11a$^{4}$ | IR11a$^{4}$ | 28 | | | | |
| Fig1-E | 5 mM Sucrose + 250 mM NaCl vs. 1 mM Sucrose | w$^{1118}$ | w$^{1118}$ | 12 | with Suc vs. without Suc | Mann-Whitney | ns | 0.4513 |
| | 250 mM NaCl vs. $H_2O$ | | | 10 | | | | |
| | 5 mM Sucrose + 250 mM LiCl vs. 1 mM Sucrose | | | 12 | | | *** | <0.0001 |
| | 250 mM LiCl vs. $H_2O$ | | | 12 | | | | |
| | 5 mM Sucrose + 250 mM KCl vs. 1 mM Sucrose | | | 11 | | | *** | 0.0001 |
| | 250 mM KCl vs. $H_2O$ | | | 11 | | | | |
| | 5 mM Sucrose + 50 mM $CaCl_2$ vs. 1 mM Sucrose | | | 11 | | | *** | <0.0001 |
| | 50 mM $CaCl_2$ vs. $H_2O$ | | | 11 | | | | |
| Fig1-F | 5 mM Sucrose + 50 mM LiCl vs. 1 mM Sucrose | IR11a$^{4}$/+ | IR11a$^{4}$/+ | 39 | IR11a$^{4}$/+ vs. IR11a$^{4}$ | Mann-Whitney | ns | 0.7367 |
| | | IR11a$^{4}$/IR11a$^{4}$ | IR11a$^{4}$ | 23 | | | | |
| | 5 mM Sucrose + 250 mM LiCl vs. 1 mM Sucrose | IR11a$^{4}$/+ | IR11a$^{4}$/+ | 30 | IR11a$^{4}$/+ vs. IR11a$^{4}$ | | *** | 0.0001 |
| | | IR11a$^{4}$/IR11a$^{4}$ | IR11a$^{4}$ | 30 | | | | |
| | 5 mM Sucrose + 50 mM KCl vs. 1 mM Sucrose | IR11a$^{4}$/+ | IR11a$^{4}$/+ | 33 | IR11a$^{4}$/+ vs. IR11a$^{4}$ | | ns | 0.219 |
| | | IR11a$^{4}$/IR11a$^{4}$ | IR11a$^{4}$ | 40 | | | | |
| | 5 mM Sucrose + 250 mM KCl vs. 1 mM Sucrose | IR11a$^{4}$/+ | IR11a$^{4}$/+ | 28 | IR11a$^{4}$/+ vs. IR11a$^{4}$ | | ns | 0.6609 |
| | | IR11a$^{4}$/IR11a$^{4}$ | IR11a$^{4}$ | 28 | | | | |
| | 5 mM Sucrose + 50 mM $MgCl_2$ vs. 1 mM Sucrose | IR11a$^{4}$/+ | IR11a$^{4}$/+ | 26 | IR11a$^{4}$/+ vs. IR11a$^{4}$ | | ns | 0.2887 |
| | | IR11a$^{4}$/IR11a$^{4}$ | IR11a$^{4}$ | 29 | | | | |
| | 5 mM Sucrose + 250 mM $CaCl_2$ vs. 1 mM Sucrose | IR11a$^{4}$/+ | IR11a$^{4}$/+ | 28 | IR11a$^{4}$/+ vs. IR11a$^{4}$ | | ns | 0.1228 |
| | | IR11a$^{4}$/IR11a$^{4}$ | IR11a$^{4}$ | 28 | | | | |
| | 5 mM Sucrose + 0.5 mM Quinine vs. 1 mM Sucrose | IR11a$^{4}$/+ | IR11a$^{4}$/+ | 28 | IR11a$^{4}$/+ vs. IR11a$^{4}$ | | ns | 0.4238 |
| | | IR11a$^{4}$/IR11a$^{4}$ | IR11a$^{4}$ | 28 | | | | |
| | 5 mM Sucrose + 0.1 mM Denatonium vs. 1 mM Sucrose | IR11a$^{4}$/+ | IR11a$^{4}$/+ | 30 | IR11a$^{4}$/+ vs. IR11a$^{4}$/IR11a$^{4}$ | | ns | 0.4425 |
| | | IR11a$^{4}$/IR11a$^{4}$ | IR11a$^{4}$ | 30 | | | | |
| Fig1-G | 250 mM NaCl vs. $H_2O$ | wcs | wcs | 35 | wcs vs. UAS-IR11a;IR11a4 | Kruskal-Wallis, with Dunn's multiple comparisons test | *** | <0.0001 |
| | | IR11a$^{4}$ / IR11a$^{4}$;UAS-IR11a/+ | UAS-IR11a;IR11a$^{4}$ | 45 | wcs vs. IR11a-GAL4;IR11a4 | | *** | <0.0001 |
| | | IR11a$^{4}$ / IR11a$^{4}$;;IR11a-GAL4/+ | IR11a-GAL4;IR11a$^{4}$ | 36 | wcs vs.IR11a-GAL4$^{rescue}$ | | ns | >0.9999 |
| | | IR11a$^{4}$ / IR11a$^{4}$;UAS-IR11a/+;IR11a-GAL4/+ | IR11a-GAL4$^{rescue}$ | 40 | IR11a$^{4}$;UAS-IR11a/+ vs. IR11a$^{4}$;IR11a-GAL4/+ | | ns | >0.9999 |
| | | | | | IR11a$^{4}$;UAS-IR11a/+ vs. IR11a-GAL4$^{rescue}$ | | *** | <0.0001 |
| | | | | | IR11a$^{4}$;;IR11a-GAL4/+ vs. IR11a-GAL4$^{rescue}$ | | *** | 0.0002 |
| Fig1-H | 5 mM Sucrose + 250 mM LiCl vs. 1 mM Sucrose | wcs | wcs | 35 | wcs vs. UAS-IR11a;IR11a4 | Kruskal-Wallis, with Dunn's multiple comparisons test | * | 0.0164 |
| | | IR11a$^{4}$ / IR11a$^{4}$;UAS-IR11a/+ | UAS-IR11a;IR11a$^{4}$ | 45 | wcs vs. IR11a-GAL4;IR11a4 | | *** | 0.0001 |
| | | IR11a$^{4}$ / IR11a$^{4}$;;IR11a-GAL4/+ | IR11a-GAL4;IR11a$^{4}$ | 36 | wcs vs.IR11a-GAL4$^{rescue}$ | | ns | >0.9999 |
| | | IR11a$^{4}$ / IR11a$^{4}$;UAS-IR11a/+;IR11a-GAL4/+ | IR11a-GAL4$^{rescue}$ | 40 | IR11a$^{4}$;UAS-IR11a/+ vs. IR11a$^{4}$;IR11a-GAL4/+ | | ns | 0.8054 |
| | | | | | IR11a$^{4}$;UAS-IR11a/+ vs. IR11a-GAL4$^{rescue}$ | | ns | 0.0864 |
| | | | | | IR11a$^{4}$;;IR11a-GAL4/+ vs. IR11a-GAL4$^{rescue}$ | | *** | 0.001 |
| Fig1-I | 150 mM NaCl vs. $H_2O$ | w$^{1118}$ | w$^{1118}$ | 30 | w$^{1118}$ vs. IR7c$^{3}$ | | ns | 0.6475 |
| | | IR7c$^{3}$/IR7c$^{3}$ | IR7c$^{3}$/IR7c$^{3}$ | 30 | w$^{1118}$ vs. IR11a$^{4}$ | Kruskal-Wallis, with Dunn's multiple comparisons test | ns | >0.9999 |
| | | IR11a$^{4}$/IR11a$^{4}$ | IR11a$^{4}$/IR11a$^{4}$ | 30 | w$^{1118}$ vs. IR7c$^{3}$,IR11a$^{4}$ | | **** | <0.0001 |
| | | IR7c$^{3}$,IR11a$^{4}$ / IR7c$^{3}$,IR11a$^{4}$ | IR7c$^{3}$,IR11a$^{4}$ / IR7c$^{3}$,IR11a$^{4}$ | 29 | IR7c$^{3}$ vs. IR11a$^{4}$ | | ns | >0.9999 |
| | | | | | IR7c$^{3}$ vs. IR7c$^{3}$,IR11a$^{4}$ | | ** | 0.0067 |
| | | | | | IR11a$^{4}$ vs. IR7c$^{3}$,IR11a$^{4}$ | | ** | 0.0019 |
| | 250 mM NaCl vs. $H_2O$ | w$^{1118}$ | w$^{1118}$ | 30 | w$^{1118}$ vs. IR7c$^{3}$ | Kruskal-Wallis, with Dunn's multiple comparisons test | **** | <0.0001 |
| | | IR7c$^{3}$/IR7c$^{3}$ | IR7c$^{3}$/IR7c$^{3}$ | 30 | w$^{1118}$ vs. IR11a$^{4}$ | | ** | 0.0016 |
| | | IR11a$^{4}$/IR11a$^{4}$ | IR11a$^{4}$/IR11a$^{4}$ | 30 | w$^{1118}$ vs. IR7c$^{3}$,IR11a$^{4}$ | | **** | <0.0001 |
| | | IR7c$^{3}$,IR11a$^{4}$ / IR7c$^{3}$,IR11a$^{4}$ | IR7c$^{3}$,IR11a$^{4}$ / | 30 | IR7c$^{3}$ vs. IR11a$^{4}$ | | ns | 0.0541 |
| | | | | | IR7c$^{3}$ vs. IR7c$^{3}$,IR11a$^{4}$ | | ns | 0.1209 |
| | | | | | IR11a$^{4}$ vs. IR7c$^{3}$,IR11a$^{4}$ | | **** | <0.0001 |
| | 400 mM NaCl vs. $H_2O$ | w$^{1118}$ | w$^{1118}$ | 30 | w$^{1118}$ vs. IR7c$^{3}$ | Kruskal-Wallis, with Dunn's multiple comparisons test | *** | 0.0006 |
| | | IR7c$^{3}$/IR7c$^{3}$ | IR7c$^{3}$/IR7c$^{3}$ | 29 | w$^{1118}$ vs. IR11a$^{4}$ | | ** | 0.004 |
| | | IR11a$^{4}$/IR11a$^{4}$ | IR11a$^{4}$/IR11a$^{4}$ | 30 | w$^{1118}$ vs. IR7c$^{3}$,IR11a$^{4}$ | | **** | <0.0001 |
| | | IR7c$^{3}$,IR11a$^{4}$ / IR7c$^{3}$,IR11a$^{4}$ | IR7c$^{3}$,IR11a$^{4}$ / IR7c$^{3}$,IR11a$^{4}$ | 28 | IR7c$^{3}$ vs. IR11a$^{4}$ | | ns | >0.9999 |
| | | | | | IR7c$^{3}$ vs. IR7c$^{3}$,IR11a$^{4}$ | | ns | >0.9999 |
| | | | | | IR11a$^{4}$ vs. IR7c$^{3}$,IR11a$^{4}$ | | ns | 0.6658 |
| Fig1-J | 5 mM Sucrose + 150 mM NaCl vs. 1 mM Sucrose | w$^{1118}$ | w$^{1118}$ | 26 | w$^{1118}$ vs. IR7c$^{3}$ | Kruskal-Wallis, with Dunn's multiple comparisons test | ns | 0.3363 |
| | | IR7c$^{3}$/IR7c$^{3}$ | IR7c$^{3}$/IR7c$^{3}$ | 30 | w$^{1118}$ vs. IR11a$^{4}$ | | ns | >0.9999 |
| | | IR11a$^{4}$/IR11a$^{4}$ | IR11a$^{4}$/IR11a$^{4}$ | 30 | w$^{1118}$ vs. IR7c$^{3}$,IR11a$^{4}$ | | ** | 0.0098 |
| | | IR7c$^{3}$,IR11a$^{4}$ / IR7c$^{3}$,IR11a$^{4}$ | IR7c$^{3}$,IR11a$^{4}$ / IR7c$^{3}$,IR11a$^{4}$ | 30 | IR7c$^{3}$ vs. IR11a$^{4}$ | | ns | 0.5533 |
| | | | | | IR7c$^{3}$ vs. IR7c$^{3}$,IR11a$^{4}$ | | ns | >0.9999 |
| | | | | | IR11a$^{4}$ vs. IR7c$^{3}$,IR11a$^{4}$ | | * | 0.0179 |
| | 5 mM Sucrose + 250 mM NaCl vs. 1 mM Sucrose | w$^{1118}$ | w$^{1118}$ | 30 | w$^{1118}$ vs. IR7c$^{3}$ | Kruskal-Wallis, with Dunn's multiple comparisons test | **** | <0.0001 |
| | | IR7c$^{3}$/IR7c$^{3}$ | IR7c$^{3}$/IR7c$^{3}$ | 30 | w$^{1118}$ vs. IR11a$^{4}$ | | ** | 0.0094 |
| | | IR11a$^{4}$/IR11a$^{4}$ | IR11a$^{4}$/IR11a$^{4}$ | 30 | w$^{1118}$ vs. IR7c$^{3}$,IR11a$^{4}$ | | **** | <0.0001 |
| | | IR7c$^{3}$,IR11a$^{4}$ / IR7c$^{3}$,IR11a$^{4}$ | IR7c$^{3}$,IR11a$^{4}$ / IR7c$^{3}$,IR11a$^{4}$ | 30 | IR7c$^{3}$ vs. IR11a$^{4}$ | | ns | 0.0854 |
| | | | | | IR7c$^{3}$ vs. IR7c$^{3}$,IR11a$^{4}$ | | ns | >0.9999 |
| | | | | | IR11a$^{4}$ vs. IR7c$^{3}$,IR11a$^{4}$ | | *** | 0.001 |
| | 5 mM Sucrose + 400 mM NaCl vs. 1 mM Sucrose | w$^{1118}$ | w$^{1118}$ | 30 | w$^{1118}$ vs. IR7c$^{3}$ | Kruskal-Wallis, with Dunn's multiple comparisons test | **** | <0.0001 |
| | | IR7c$^{3}$/IR7c$^{3}$ | IR7c$^{3}$/IR7c$^{3}$ | 29 | w$^{1118}$ vs. IR11a$^{4}$ | | * | 0.0103 |
| | | IR11a$^{4}$/IR11a$^{4}$ | IR11a$^{4}$/IR11a$^{4}$ | 30 | w$^{1118}$ vs. IR7c$^{3}$,IR11a$^{4}$ | | **** | <0.0001 |
| | | IR7c$^{3}$,IR11a$^{4}$ / IR7c$^{3}$,IR11a$^{4}$ | IR7c$^{3}$,IR11a$^{4}$ / IR7c$^{3}$,IR11a$^{4}$ | 28 | IR7c$^{3}$ vs. IR11a$^{4}$ | | ns | 0.1914 |
| | | | | | IR7c$^{3}$ vs. IR7c$^{3}$,IR11a$^{4}$ | | * | 0.0283 |
| | | | | | IR11a$^{4}$ vs. IR7c$^{3}$,IR11a$^{4}$ | | **** | <0.0001 |

**Data S2-Genotypes and statistical details for Fig.2**

| Figures | Foods | Genotypes | Genotype labeled in figures | N value | Compare groups | Statistical methods | P value summary | P value |
|---|---|---|---|---|---|---|---|---|
| Fig. 2-A1-A3 | standard food | *$w^{1118}$/$w^{1118}$;UAS-mcd8:RFP,LexAop-mcd8:GFP;GR66a-LexA/+;IR11a-GAL4/+* | GR66a/IR11a | 3 | NA | NA | NA | NA |
| Fig. 2-B1-B3 | standard food | *$w^{1118}$/$w^{1118}$;UAS-mcd8:RFP,LexAop-mcd8:GFP:+/+:$GR64f^{LexA}$/IR11a-GAL4* | GR64f/IR11a | 3 | NA | NA | NA | NA |
| Fig. 2-C1-C3 | standard food | *$w^{1118}$/$w^{1118}$; UAS-mcd8:GFP,IR76b-QF/+;QUAS-Tdtomato-3x HA/IR11a-GAL4* | IR76b/IR11a | 3 | NA | NA | NA | NA |
| Fig. 2-D1-D3 | standard food | *$w^{1118}$/$w^{1118}$;IR11a-GAL4/UAS-mcd8:RFP* | IR25a/IR11a | 3 | NA | NA | NA | NA |
| Fig. 2-E | 250 mM NaCl vs. $H_2O$ | *$w^{1118}$/$w^{1118}$;UAS-Kir2.1/+* | *UAS-Kir2.1 / +* | 39 | *UAS-Kir2.1/+ vs. IR11a-GAL4/+* | Kruskal-Wallis, Dunn's multiple comparisons test | * | 0.0115 |
| | | *$w^{1118}$/$w^{1118}$;;IR11a-GAL4/+* | *IR11a-GAL4 / +* | 28 | *UAS-Kir2.1/+ vs. IR11a > Kir2.1* | | ** | 0.0069 |
| | | *$w^{1118}$/$w^{1118}$;UAS-Kir2.1/+;IR11a-GAL4/+* | *IR11a > Kir2.1* | 30 | *IR11a-GAL4/+ vs. IR11a > Kir2.1* | | *** | <0.0001 |
| Fig. 2-F | 5 mM Sucrose + 250 mM LiCl vs. 1 mM Sucrose | *$w^{1118}$/$w^{1118}$;UAS-Kir2.1/+* | *UAS-Kir2.1 / +* | 22 | *UAS-Kir2.1/+ vs. IR11a-GAL4/+* | Kruskal-Wallis, Dunn's multiple comparisons test | ns | >0.9999 |
| | | *$w^{1118}$/$w^{1118}$;;IR11a-GAL4/+* | *IR11a-GAL4 / +* | 13 | *UAS-Kir2.1/+ vs. IR11a > Kir2.1* | | *** | 0.0002 |
| | | *$w^{1118}$/$w^{1118}$;UAS-Kir2.1/+;IR11a-GAL4/+* | *IR11a > Kir2.1* | 18 | *IR11a-GAL4/+ vs. IR11a > Kir2.1* | | *** | <0.0001 |
| Fig. 2-G | 5 mM Sucrose + 0.5 mM Quinine vs. 1 mM Sucrose | *$w^{1118}$/$w^{1118}$;UAS-Kir2.1/+* | *UAS-Kir2.1 / +* | 23 | *UAS-Kir2.1/+ vs. IR11a-GAL4/+* | Kruskal-Wallis, Dunn's multiple comparisons test | ns | >0.9999 |
| | | *$w^{1118}$/$w^{1118}$;;IR11a-GAL4/+* | *IR11a-GAL4 / +* | 24 | *UAS-Kir2.1/+ vs. IR11a > Kir2.1* | | ns | >0.9999 |
| | | *$w^{1118}$/$w^{1118}$;UAS-Kir2.1/+;IR11a-GAL4/+* | *IR11a > Kir2.1* | 23 | *IR11a-GAL4/+ vs. IR11a > Kir2.1* | | ns | >0.9999 |
| Fig. 2-H | 5 mM Sucrose + 0.1 mM Denatonium vs. 1 mM Sucrose | *$w^{1118}$/$w^{1118}$;UAS-Kir2.1/+* | *UAS-Kir2.1 / +* | 24 | *UAS-Kir2.1/+ vs. IR11a-GAL4/+* | Kruskal-Wallis, Dunn's multiple comparisons test | ns | >0.9999 |
| | | *$w^{1118}$/$w^{1118}$;;IR11a-GAL4/+* | *IR11a-GAL4 / +* | 25 | *UAS-Kir2.1/+ vs. IR11a > Kir2.1* | | ns | 0.8752 |
| | | *$w^{1118}$/$w^{1118}$;UAS-Kir2.1/+;IR11a-GAL4/+* | *IR11a > Kir2.1* | 24 | *IR11a-GAL4/+ vs. IR11a > Kir2.1* | | ns | >0.9999 |
| Fig. 2-I | 0.1 mM Capsaicin + 1 % Ethanol vs.1 % Ethanol | *$w^{1118}$/$w^{1118}$;UAS-VR1/+* | *UAS-VR1 / +* | 20 | *UAS-Kir2.1/+ vs. IR11a-GAL4/+* | Kruskal-Wallis, Dunn's multiple comparisons test | ns | >0.9999 |
| | | *$w^{1118}$/$w^{1118}$;;IR11a-GAL4/+* | *IR11a-GAL4 / +* | 18 | *UAS-Kir2.1/+ vs. IR11a > Kir2.1* | | *** | <0.001 |
| | | *$w^{1118}$/$w^{1118}$;UAS-VR1/+;IR11a-GAL4/+* | *IR11a > VR1* | 28 | *IR11a-GAL4/+ vs. IR11a > Kir2.1* | | *** | <0.001 |

**Data S3-Genotypes and statistical details for Fig.3**

| Figures | Test chemicals | Genotypes | Genotypes labeled in figures | N value | Compare groups | Statistical methods | P value summary | P value |
|---|---|---|---|---|---|---|---|---|
| Fig. 3-A, B | $H_2O$ | GR33a$^{GAL4}$,UAS-GCaMP6m/+ | GR33a > GCaMP6M | 8 | | Mann-Whitney | | |
| | 50 mM NaCl | | | 8 | 50 mM NaCl vs. H2O | | **ns** | 0.7984 |
| | 150 mM NaCl | | | 9 | 150 mM NaCl vs. H2O | | ** | 0.0037 |
| | 250 mM NaCl | | | 24 | 250 mM NaCl vs. H2O | | *** | <0.0001 |
| | 500 mM NaCl | | | 16 | 500 mM NaCl vs. H2O | | *** | <0.0001 |
| Fig. 3-C | H2O | IR11a$^{4}$/+;GR33a$^{GAL4}$,UAS-GCaMP6m/+ | IR11a$^{4}$ / + | 9 | | Mann-Whitney | | |
| | 50 mM LiCl | | | 12 | 50 mM LiCl vs. H2O | | ns | 0.7283 |
| | 50 mM KCl | | | 9 | 50 mM KCl vs. H2O | | ns | 0.1641 |
| | 250 mM NMDG-Cl | | | 9 | 250 mM NMDG-Cl vs. H2O | | ns | 0.1858 |
| | 250 mM LiCl | | | 10 | 250 mM LiCl vs. H2O | | *** | <0.0001 |
| | 250 mM NaCl | | | 10 | 250 mM NaCl vs. H2O | | *** | 0.0002 |
| | 250 mM NaBr | | | 12 | 250 mM NaBr vs. H2O | | *** | <0.0001 |
| | 250 mM KCl | | | 8 | 250 mM KCl vs. H2O | | ** | 0.0077 |
| | 50 mM $CaCl_2$ | | | 9 | 50 mM $CaCl_2$ vs. H2O | | *** | <0.0001 |
| | 1 mM Denatonium | | | 11 | 1 mM Denatonium vs. H2O | | *** | <0.0001 |
| | $H_2O$ | IR11a$^{4}$/+;GR33a$^{GAL4}$,UAS-GCaMP6m/+ | IR11a$^{4}$ / + | 9 | IR11a$^{4}$/+ vs. IR11a$^{4}$ mutants | Kruskal-Wallis, with Dunn's multiple comparisons test | ns | >0.9999 |
| | | IR11a$^{4}$ / IR11a$^{4}$;GR33a$^{GAL4}$,UAS-GCaMP6m/+ | IR11a$^{4}$ | 8 | IR11a$^{4}$/+ vs. IR11a$^{4}$ rescue | | ns | 0.1457 |
| | | IR11a$^{4}$ / IR11a$^{4}$;GR33a$^{GAL4}$,UAS-GCaMP6m/UAS-IR11a | IR11a$^{4}$ rescue | 11 | IR11a$^{4}$ mutants vs. IR11a$^{4}$ rescue | | ns | 0.6619 |
| | 250 mM LiCl | IR11a$^{4}$/+;GR33a$^{GAL4}$,UAS-GCaMP6m/+ | IR11a$^{4}$ / + | 10 | IR11a$^{4}$/+ vs. IR11a$^{4}$ mutants | Kruskal-Wallis, with Dunn's multiple comparisons test | **** | <0.0001 |
| | | IR11a$^{4}$ / IR11a$^{4}$;GR33a$^{GAL4}$,UAS-GCaMP6m/+ | IR11a$^{4}$ | 9 | IR11a$^{4}$/+ vs. IR11a$^{4}$ rescue | | ns | 0.3706 |
| | | IR11a$^{4}$ / IR11a$^{4}$;GR33a$^{GAL4}$,UAS-GCaMP6m/UAS-IR11a | IR11a$^{4}$ rescue | 14 | IR11a$^{4}$ mutants vs. IR11a$^{4}$ rescue | | ** | 0.0017 |
| | 250 mM NaCl | IR11a$^{4}$/+;GR33a$^{GAL4}$,UAS-GCaMP6m/+ | IR11a$^{4}$ / + | 10 | IR11a$^{4}$/+ vs. IR11a$^{4}$ mutants | Kruskal-Wallis, with Dunn's multiple comparisons test | ** | 0.0027 |
| | | IR11a$^{4}$ / IR11a$^{4}$;GR33a$^{GAL4}$,UAS-GCaMP6m/+ | IR11a$^{4}$ | 8 | IR11a$^{4}$/+ vs. IR11a$^{4}$ rescue | | ns | >0.9999 |
| | | IR11a$^{4}$ / IR11a$^{4}$;GR33a$^{GAL4}$,UAS-GCaMP6m/UAS-IR11a | IR11a$^{4}$ rescue | 16 | IR11a$^{4}$ mutants vs. IR11a$^{4}$ rescue | | * | 0.0135 |
| | 250 mM NaBr | IR11a$^{4}$/+;GR33a$^{GAL4}$,UAS-GCaMP6m/+ | IR11a$^{4}$ / + | 12 | IR11a$^{4}$/+ vs. IR11a$^{4}$ mutants | Kruskal-Wallis, with Dunn's multiple comparisons test | * | 0.0416 |
| | | IR11a$^{4}$ / IR11a$^{4}$;GR33a$^{GAL4}$,UAS-GCaMP6m/+ | IR11a$^{4}$ | 10 | IR11a$^{4}$/+ vs. IR11a$^{4}$ rescue | | ns | 0.4865 |
| | | IR11a$^{4}$ / IR11a$^{4}$;GR33a$^{GAL4}$,UAS-GCaMP6m/UAS-IR11a | IR11a$^{4}$ rescue | 15 | IR11a$^{4}$ mutants vs. IR11a$^{4}$ rescue | | ** | 0.0015 |
| | 250 mM KCl | IR11a$^{4}$/+;GR33a$^{GAL4}$,UAS-GCaMP6m/+ | IR11a$^{4}$ / + | 8 | IR11a$^{4}$/+ vs. IR11a$^{4}$ mutants | Mann-Whitney | ns | 0.9039 |
| | | IR11a$^{4}$ / IR11a$^{4}$;GR33a$^{GAL4}$,UAS-GCaMP6m/+ | IR11a$^{4}$ | 11 | | | | |
| | 50 mM $CaCl_2$ | IR11a$^{4}$/+;GR33a$^{GAL4}$,UAS-GCaMP6m/+ | IR11a$^{4}$ / + | 9 | IR11a$^{4}$/+ vs. IR11a$^{4}$ mutants | | ns | 0.7304 |
| | | IR11a$^{4}$ / IR11a$^{4}$;GR33a$^{GAL4}$,UAS-GCaMP6m/+ | IR11a$^{4}$ | 9 | | | | |
| | 1 mM Denatonium | IR11a$^{4}$/+;GR33a$^{GAL4}$,UAS-GCaMP6m/+ | IR11a$^{4}$ / + | 11 | IR11a$^{4}$/+ vs. IR11a$^{4}$ mutants | | ns | 0.3144 |
| | | IR11a$^{4}$ / IR11a$^{4}$;GR33a$^{GAL4}$,UAS-GCaMP6m/+ | IR11a$^{4}$ | 10 | | | | |
| Fig. 3-D | $H_2O$ | w$^{1118}$/w$^{1118}$;IR25a$^{2}$,GR33a$^{GAL4}$/UAS-GCaMP6m | IR25a$^{2}$ / + | 9 | IR25a$^{2}$/+ vs. IR25a$^{2}$ mutants | Kruskal-Wallis, with Dunn's multiple comparisons test | ns | >0.9999 |
| | | w$^{1118}$/w$^{1118}$;IR25a$^{2}$,GR33a$^{GAL4}$/IR25a$^{2}$;UAS-GCaMP6m/+ | IR25a$^{2}$ | 8 | IR25a$^{2}$/+ vs. IR25a$^{2}$ rescue | | ns | 0.4045 |
| | | w$^{1118}$/w$^{1118}$;IR25a$^{2}$,GR33a$^{GAL4}$/IR25a$^{2}$,UAS-IR25a;UAS-GCaMP6m/+ | IR25a$^{2}$ rescue | 12 | IR25a$^{2}$ mutants vs. IR25a$^{2}$ rescue | | ns | 0.1689 |
| | 250 mM LiCl | w$^{1118}$/w$^{1118}$;IR25a$^{2}$,GR33a$^{GAL4}$/UAS-GCaMP6m | IR25a$^{2}$ / + | 8 | IR25a$^{2}$/+ vs. IR25a$^{2}$ | Kruskal-Wallis, with Dunn's multiple comparisons test | * | 0.0441 |
| | | w$^{1118}$/w$^{1118}$;IR25a$^{2}$,GR33a$^{GAL4}$/IR25a$^{2}$;UAS-GCaMP6m/+ | IR25a$^{2}$ | 8 | IR25a$^{2}$/+ vs. IR25a$^{2}$ rescue | | ns | >0.9999 |
| | | w$^{1118}$/w$^{1118}$;IR25a$^{2}$,GR33a$^{GAL4}$/IR25a$^{2}$,UAS-IR25a;UAS-GCaMP6m/+ | IR25a$^{2}$ rescue | 8 | IR25a$^{2}$ mutants vs. IR25a$^{2}$ rescue | | ** | 0.0021 |
| | 250 mM NaCl | w$^{1118}$/w$^{1118}$;IR25a$^{2}$,GR33a$^{GAL4}$/UAS-GCaMP6m | IR25a$^{2}$ / + | 21 | IR25a$^{2}$/+ vs. IR25a$^{2}$ | Kruskal-Wallis, with Dunn's multiple comparisons test | ** | 0.0089 |
| | | w$^{1118}$/w$^{1118}$;IR25a$^{2}$,GR33a$^{GAL4}$/IR25a$^{2}$;UAS-GCaMP6m/+ | IR25a$^{2}$ | 13 | IR25a$^{2}$/+ vs. IR25a$^{2}$ rescue | | ns | >0.9999 |
| | | w$^{1118}$/w$^{1118}$;IR25a$^{2}$,GR33a$^{GAL4}$/IR25a$^{2}$,UAS-IR25a;UAS-GCaMP6m/+ | IR25a$^{2}$ rescue | 15 | IR25a$^{2}$ mutants vs. IR25a$^{2}$ rescue | | * | 0.0149 |
| | 250 mM KCl | w$^{1118}$/w$^{1118}$;IR25a$^{2}$,GR33a$^{GAL4}$/UAS-GCaMP6m | IR25a$^{2}$ / + | 10 | IR25a$^{2}$/+ vs. IR25a$^{2}$ | Mann-Whitney | ns | 0.0952 |
| | | w1118/w1118;IR25a2,GR33aGAL4/IR25a2;UAS-GCaMP6m/+ | IR25a$^{2}$ | 11 | | | | |
| | 50 mM $CaCl_2$ | w$^{1118}$/w$^{1118}$;IR25a$^{2}$,GR33a$^{GAL4}$/UAS-GCaMP6m | IR25a$^{2}$ / + | 10 | IR25a$^{2}$/+ vs. IR25a$^{2}$ | | ns | 0.9682 |
| | | w$^{1118}$/w$^{1118}$;IR25a$^{2}$,GR33a$^{GAL4}$/IR25a$^{2}$;UAS-GCaMP6m/+ | IR25a$^{2}$ | 9 | | | | |
| | 1 mM Denatonium | w$^{1118}$/w$^{1118}$;IR25a$^{2}$,GR33a$^{GAL4}$/UAS-GCaMP6m | IR25a$^{2}$ / + | 10 | IR25a$^{2}$/+ vs. IR25a$^{2}$ | | ns | 0.2475 |
| | | w$^{1118}$/w$^{1118}$;IR25a$^{2}$,GR33a$^{GAL4}$/IR25a$^{2}$;UAS-GCaMP6m/+ | IR25a$^{2}$ | 10 | | | | |
| Fig. 3-E | $H_2O$ | w$^{1118}$/w$^{1118}$;GR33a$^{GAL4}$,UAS-GCaMP6m/+;IR76b$^{1}$/+ | IR76b$^{1}$ / + | 13 | IR76b$^{1}$ + vs. IR76b$^{1}$ | Kruskal-Wallis, with Dunn's multiple comparisons test | ns | 0.4156 |
| | | w$^{1118}$/w$^{1118}$;GR33a$^{GAL4}$,UAS-GCaMP6m/+ ;IR76b$^{1}$/ IR76b$^{1}$ | IR76b$^{1}$ | 9 | IR76b$^{1}$/+ vs. IR76b$^{1}$ rescue | | ns | 0.1284 |
| | | w$^{1118}$/w$^{1118}$;GR33a$^{GAL4}$,UAS-GCaMP6m/UAS-IR76b ;IR76b$^{1}$/ IR76b$^{1}$ | IR76b$^{1}$ rescue | 10 | IR76b$^{1}$ mutants vs. IR76b$^{1}$ rescue | | ns | >0.9999 |
| | 250 mM LiCl | w$^{1118}$/w$^{1118}$;GR33a$^{GAL4}$,UAS-GCaMP6m/+;IR76b$^{1}$/+ | IR76b$^{1}$ / + | 14 | IR76b$^{1}$ + vs. IR76b$^{1}$ | Kruskal-Wallis, with Dunn's multiple comparisons test | ** | 0.0082 |
| | | w$^{1118}$/w$^{1118}$;GR33a$^{GAL4}$,UAS-GCaMP6m/+ ;IR76b$^{1}$/ IR76b$^{1}$ | IR76b$^{1}$ | 13 | IR76b$^{1}$/+ vs. IR76b$^{1}$ rescue | | ns | >0.9999 |
| | | w$^{1118}$/w$^{1118}$;GR33a$^{GAL4}$,UAS-GCaMP6m/UAS-IR76b ;IR76b$^{1}$/ IR76b$^{1}$ | IR76b$^{1}$ rescue | 18 | IR76b$^{1}$ mutants vs. IR76b$^{1}$ rescue | | * | 0.0491 |
| | 250 mM NaCl | w$^{1118}$/w$^{1118}$;GR33a$^{GAL4}$,UAS-GCaMP6m/+;IR76b$^{1}$/+ | IR76b$^{1}$ / + | 13 | IR76b$^{1}$ + vs. IR76b$^{1}$ | Kruskal-Wallis, with Dunn's multiple comparisons test | ns | 0.7151 |
| | | w$^{1118}$/w$^{1118}$;GR33a$^{GAL4}$,UAS-GCaMP6m/+ ;IR76b$^{1}$/ IR76b$^{1}$ | IR76b$^{1}$ | 8 | IR76b$^{1}$/+ vs. IR76b$^{1}$ rescue | | ns | 0.1413 |
| | | w$^{1118}$/w$^{1118}$;GR33a$^{GAL4}$,UAS-GCaMP6m/UAS-IR76b ;IR76b$^{1}$/ IR76b$^{1}$ | IR76b$^{1}$ rescue | 10 | IR76b$^{1}$ mutants vs. IR76b$^{1}$ rescue | | ns | >0.9999 |
| | 250 mM KCl | w$^{1118}$/w$^{1118}$;GR33a$^{GAL4}$,UAS-GCaMP6m/+;IR76b$^{1}$/+ | IR76b$^{1}$ / + | 10 | IR76b$^{1}$/+ vs. IR76b$^{1}$ | Mann-Whitney | ns | 0.6305 |
| | | w$^{1118}$/w$^{1118}$;GR33a$^{GAL4}$,UAS-GCaMP6m/+ ;IR76b$^{1}$/ IR76b$^{1}$ | IR76b$^{1}$ | 10 | | | | |
| | 50 mM $CaCl_2$ | w$^{1118}$/w$^{1118}$;GR33a$^{GAL4}$,UAS-GCaMP6m/+;IR76b$^{1}$/+ | IR76b$^{1}$ / + | 9 | IR76b$^{1}$ + vs. IR76b$^{1}$ | | ns | 0.6665 |
| | | w$^{1118}$/w$^{1118}$;GR33a$^{GAL4}$,UAS-GCaMP6m/+ ;IR76b$^{1}$/ IR76b$^{1}$ | IR76b$^{1}$ | 9 | | | | |
| | 1 mM Denatonium | w$^{1118}$/w$^{1118}$;GR33a$^{GAL4}$,UAS-GCaMP6m/+;IR76b$^{1}$/+ | IR76b$^{1}$ / + | 10 | IR76b$^{1}$ + vs. IR76b$^{1}$ | | ns | 0.4598 |
| | | w$^{1118}$/w$^{1118}$;GR33a$^{GAL4}$,UAS-GCaMP6m/+ ;IR76b$^{1}$/ IR76b$^{1}$ | IR76b$^{1}$ | 8 | | | | |
| Fig. 3-G | $H_2O$ | w$^{1118}$/w$^{1118}$;GR33a$^{GAL4}$,UAS-GCaMP6m/+ | GR33a > GCaMP6M | 8 | | Mann-Whitney | | |
| | 250 mM LiCl | | | 9 | 250 mM LiCl vs. H2O | | *** | 0.0006 |
| | 250 mM NaCl | | | 11 | 250 mM NaCl vs. H2O | | *** | <0.0001 |
| | 250 mM KCl | | | 9 | 250 mM KCl vs. H2O | | ** | 0.0037 |
| | 1 mM Denatonium | | | 9 | 1 mM Denatonium vs. H2O | | *** | 0.0002 |
| | $H_2O$ | w$^{1118}$/w$^{1118}$;GR33a$^{GAL4}$,UAS-GCaMP6m/+ | GR33a > GCaMP6M | 8 | $H_2O$ + 5mM Amiloride vs. H2O | Mann-Whitney | ns | 0.3823 |
| | $H_2O$ + 5mM Amiloride | | | 8 | | | | |
| | 250 mM LiCl | | | 9 | 250 mM LiCl + 5mM Amiloride vs. 250 mM Li | | ns | 0.482 |
| | 250 mM LiCl + 5mM Amiloride | | | 15 | | | | |
| | 250 mM NaCl | | | 11 | 250 mM NaCl + 5mM Amiloride vs. 250 mM NaCl | | **** | <0.0001 |
| | 250 mM NaCl + 5mM Amiloride | | | 13 | | | | |
| | 250 mM KCl | | | 9 | 250 mM KCl + 5mM Amiloride vs. 250 mM KCl | | ns | 0.1359 |
| | 250 mM KCl + 5mM Amiloride | | | 9 | | | | |
| | 1 mM Denatonium | | | 9 | 1 mM Denatonium + 5mM Amiloride vs. 1 mM Denatonium | | ns | 0.4807 |
| | 1 mM Denatonium + 5mM Amiloride | | | 8 | | | | |
| Fig. 3-H | $H_2O$ | w$^{1118}$/w$^{1118}$;UAS-GCaMP6m/+;IR7c-GAL4/+ | IR7c > GCaMP6M | 11 | | Mann-Whitney | | |
| | 250 mM LiCl | | | 9 | 250 mM LiCl vs. H2O | | ns | 0.1754 |
| | 250 mM NaCl | | | 9 | 250 mM NaCl vs. H2O | | ** | 0.0012 |
| | 250 mM KCl | | | 8 | 250 mM KCl vs. H2O | | * | 0.0259 |
| | $H_2O$ | w$^{1118}$/w$^{1118}$;UAS-GCaMP6m/+;IR7c-GAL4/+ | IR7c > GCaMP6M | 11 | $H_2O$ + 5mM Amiloride vs. H2O | Mann-Whitney | ns | 0.5027 |
| | $H_2O$ + 5mM Amiloride | | | 9 | | | | |
| | 250 mM LiCl | | | 9 | 250 mM LiCl + 5mM Amiloride vs. 250 mM Li | | ns | 0.3401 |
| | 250 mM LiCl + 5mM Amiloride | | | 9 | | | | |
| | 250 mM NaCl | | | 9 | 250 mM NaCl + 5mM Amiloride vs. 250 mM NaCl | | ns | 0.211 |
| | 250 mM NaCl + 5mM Amiloride | | | 10 | | | | |
| | 250 mM KCl | | | 8 | 250 mM KCl + 5mM Amiloride vs. 250 mM KCl | | ns | 0.4807 |
| | 250 mM KCl + 5mM Amiloride | | | 9 | | | | |

| Data S4-Genotypes and statistical details for Fig.4 | | | | | | | | |
|---|---|---|---|---|---|---|---|---|
| **Figures** | **Test Chemicals** | **Genotypes** | **Genotypes labeled in figures** | **N value** | **Compare groups** | **Statistical methods** | **P value summary** | **P value** |
| Fig. 4-A1-A3 | | *$w^{1118}$/$w^{1118}$; UAS-mcd8:GFP,IR76b-QF/+;QUAS-Tdtomato-3x HA/Ppk23-GAL4* | Ppk23/IR76b | 4 | NA | NA | NA | NA |
| Fig. 4-B1-B3 | | *$w^{1118}$/$w^{1118}$; UAS-mcd8:GFP/+;Ppk23-GAL4/+* | Ppk23/IR25a | 4 | NA | NA | NA | NA |
| Fig. 4-C | $H_2O$ | *$w^{1118}$/$w^{1118}$;;Ppk23-GAL4,UAS-GCaMP6m/+* | *Ppk23-GAL4 / +* | 8 | | | | |
| | 50 mM LiCl | | | 9 | $H_2O$ vs. 50 mM LiCl | Mann-Whitney | ns | 0.6058 |
| | 250 mM LiCl | | | 9 | $H_2O$ vs. 250 mM LiCl | | ns | 0.2359 |
| | 50 mM NaCl | | | 12 | $H_2O$ vs. 50 mM NaCl | | ns | 0.9699 |
| | 250 mM NaCl | | | 16 | $H_2O$ vs. 250 mM NaCl | | ns | 0.4523 |
| | 50 mM KCl | | | 10 | $H_2O$ vs. 50 mM KCl | | ns | 0.5726 |
| | 250 mM KCl | | | 12 | $H_2O$ vs. 250 mM KCl | | ns | 0.1813 |
| | $H_2O$ | *$w^{1118}$/$w^{1118}$;;Ppk23-GAL4,UAS-GCaMP6m/+* | *Ppk23-GAL4 / +* | 8 | *Ppk23-GAL4/+ vs. Ppk23 > IR11a* | Mann-Whitney | ns | 0.9626 |
| | | *$w^{1118}$/$w^{1118}$;UAS-IR11a/+;Ppk23-GAL4,UAS-GCaMP6m/+* | *Ppk23 > IR11a* | 9 | | | | |
| | 50 mM LiCl | *$w^{1118}$/$w^{1118}$;;Ppk23-GAL4,UAS-GCaMP6m/+* | *Ppk23-GAL4 / +* | 9 | *Ppk23-GAL4/+ vs. Ppk23 > IR11a* | | ns | 0.7962 |
| | | *$w^{1118}$/$w^{1118}$;UAS-IR11a/+;Ppk23-GAL4,UAS-GCaMP6m/+* | *Ppk23 > IR11a* | 9 | | | | |
| | 250 mM LiCl | *$w^{1118}$/$w^{1118}$;;Ppk23-GAL4,UAS-GCaMP6m/+* | *Ppk23-GAL4 / +* | 9 | *Ppk23-GAL4/+ vs. Ppk23 > IR11a* | | ns | 0.2071 |
| | | *$w^{1118}$/$w^{1118}$;UAS-IR11a/+;Ppk23-GAL4,UAS-GCaMP6m/+* | *Ppk23 > IR11a* | 16 | | | | |
| | 50 mM NaCl | *$w^{1118}$/$w^{1118}$;;Ppk23-GAL4,UAS-GCaMP6m/+* | *Ppk23-GAL4 / +* | 12 | *Ppk23-GAL4/+ vs. Ppk23 > IR11a* | | ns | 0.3451 |
| | | *$w^{1118}$/$w^{1118}$;UAS-IR11a/+;Ppk23-GAL4,UAS-GCaMP6m/+* | *Ppk23 > IR11a* | 9 | | | | |
| | 250 mM NaCl | *$w^{1118}$/$w^{1118}$;;Ppk23-GAL4,UAS-GCaMP6m/+* | *Ppk23-GAL4 / +* | 16 | *Ppk23-GAL4/+ vs. Ppk23 > IR11a* | | *** | 0.0003 |
| | | *$w^{1118}$/$w^{1118}$;UAS-IR11a/+;Ppk23-GAL4,UAS-GCaMP6m/+* | *Ppk23 > IR11a* | 13 | | | | |
| | 50 mM KCl | *$w^{1118}$/$w^{1118}$;;Ppk23-GAL4,UAS-GCaMP6m/+* | *Ppk23-GAL4 / +* | 10 | *Ppk23-GAL4/+ vs. Ppk23 > IR11a* | | ns | 0.6842 |
| | | *$w^{1118}$/$w^{1118}$;UAS-IR11a/+;Ppk23-GAL4,UAS-GCaMP6m/+* | *Ppk23 > IR11a* | 10 | | | | |
| | 250 mM KCl | *$w^{1118}$/$w^{1118}$;;Ppk23-GAL4,UAS-GCaMP6m/+* | *Ppk23-GAL4 / +* | 12 | *Ppk23-GAL4/+ vs. Ppk23 > IR11a* | | ns | 0.5208 |
| | | *$w^{1118}$/$w^{1118}$;UAS-IR11a/+;Ppk23-GAL4,UAS-GCaMP6m/+* | *Ppk23 > IR11a* | 8 | | | | |
| Fig. 4-D | $H_2O$ | *$w^{1118}$/$w^{1118}$;IR25a$^{2}$/UAS-IR11a;UAS-GCaMP6m,Ppk23-GAL4/+* | *IR25a$^{2}$/+ Ppk23 > IR11a* | 8 | IR25a$^{2}$/+ *Ppk23 > IR11a vs. IR25a$^{2}$ Ppk23 > IR11a* | Mann-Whitney | ns | 0.1139 |
| | | *$w^{1118}$/$w^{1118}$;IR25a$^{2}$/IR25a$^{2}$,UAS-IR11a;UAS-GCaMP6m,Ppk23-GAL4/+* | *IR25a$^{2}$* Ppk23 > IR11a | 9 | | | | |
| | 250 mM NaCl | *$w^{1118}$/$w^{1118}$;IR25a$^{2}$/UAS-IR11a;UAS-GCaMP6m,Ppk23-GAL4/+* | *IR25a$^{2}$/+ Ppk23 > IR11a* | 9 | *IR25a$^{2}$/+ Ppk23 > IR11a vs. IR25a$^{2}$ Ppk23 > IR11a* | Mann-Whitney | * | 0.0155 |
| | | *$w^{1118}$/$w^{1118}$;IR25a$^{2}$/IR25a$^{2}$,UAS-IR11a;UAS-GCaMP6m,Ppk23-GAL4/+* | *IR25a$^{2}$* Ppk23 > IR11a | 10 | | | | |
| | | | | | *IR25a$^{2}$/+ Ppk23 > IR11a* $H_2O$ *vs. IR25a$^{2}$/+ Ppk23 > IR11a* 250 mM NaCl | Mann-Whitney | ** | 0.003 |
| | | | | | *IR25a$^{2}$ Ppk23 > IR11a* $H_2O$ *vs. IR25a$^{2}$ Ppk23 > IR11a* 250 mM NaCl | Mann-Whitney | ns | 0.1823 |
| Fig. 4-E | $H_2O$ | *$w^{1118}$/$w^{1118}$;UAS-GCaMP6m/UAS-IR11a;Ppk23-GAL4,IR76b$^{1}$/+* | *IR76b$^{1}$/+ Ppk23 > IR11a* | 8 | *IR76b$^{1}$/+ Ppk23 > IR11a vs. IR76b$^{1}$ Ppk23 > IR11a* | Mann-Whitney | ns | 0.7345 |
| | | *$w^{1118}$/$w^{1118}$;UAS-GCaMP6m/UAS-IR11a;Ppk23-GAL4,IR76b$^{1}$/IR76b$^{1}$* | *IR76b$^{1}$* Ppk23 > IR11a | 12 | | | | |
| | 250 mM NaCl | *$w^{1118}$/$w^{1118}$;UAS-GCaMP6m/UAS-IR11a;Ppk23-GAL4,IR76b$^{1}$/+* | *IR76b$^{1}$/+ Ppk23 > IR11a* | 11 | IR76b$^{1}$/+ *Ppk23 > IR11a vs. IR76b$^{1}$ Ppk23 > IR11a* | Mann-Whitney | ns | 0.8801 |
| | | *$w^{1118}$/$w^{1118}$;UAS-GCaMP6m/UAS-IR11a;Ppk23-GAL4,IR76b$^{1}$/IR76b$^{1}$* | *IR76b$^{1}$* Ppk23 > IR11a | 12 | | | | |
| | | | | | *IR76b$^{1}$/+ Ppk23 > IR11a H2O vs. IR76b$^{1}$ Ppk23 > IR11a 250 mM NaCl* | Mann-Whitney | * | 0.0409 |
| | | | | | *IR76b$^{1}$* mutants *Ppk23 H2O > IR11a vs. IR76b$^{1}$ Ppk23 > IR11a 250 mM NaCl* | Mann-Whitney | ** | 0.0014 |

**Data S5-Genotypes and statistical details for Fig.5**

| Figures | Test solutions | Genotypes | Genotypes labeled in figures | N value | Compare groups | Statistical methods | P value summary | P value |
|---|---|---|---|---|---|---|---|---|
| Fig. 5-B | 10 mM Glucrose + 10 mM HEPES + 250 mM NMDG-Cl (Ctl.) | *pAc5.1-IR11a, pAc5.1-GCaMP6M, pAc5.1-GFP* | IR11a | 19 | | Mann-Whitney | | |
| | 10 mM Glucrose + 10 mM HEPES + 250 mM LiCl (250 LiCl) | | | 18 | Ctl. vs. 250 LiCl | | ns | 0.5578 |
| | 10 mM Glucrose + 10 mM HEPES + 50 mM NaCl (50 NaCl) | | | 18 | Ctl. vs. 50 NaCl | | ns | 0.2983 |
| | 10 mM Glucrose + 10 mM HEPES + 150 mM NaCl (150 NaCl) | | | 16 | Ctl. vs. 150 NaCl | | ns | 0.4611 |
| | 10 mM Glucrose + 10 mM HEPES + 250 mM NaCl (250 NaCl) | | | 20 | Ctl. vs. 250 NaCl | | ns | 0.5133 |
| | 10 mM Glucrose + 10 mM HEPES + 250 mM KCl (250 KCl) | | | 17 | Ctl. vs. 250 KCl | | ns | 0.0933 |
| Fig. 5-C | 10 mM Glucrose + 10 mM HEPES + 250 mM NMDG-Cl (Ctl.) | *pAc5.1-IR25a, pAc5.1-GCaMP6M, pAc5.1-GFP* | IR25a | 17 | | | | |
| | 10 mM Glucrose + 10 mM HEPES + 250 mM LiCl (250 LiCl) | | | 16 | Ctl. vs. 250 LiCl | | ns | 0.2604 |
| | 10 mM Glucrose + 10 mM HEPES + 50 mM NaCl (50 NaCl) | | | 16 | Ctl. vs. 50 NaCl | | ns | 0.2604 |
| | 10 mM Glucrose + 10 mM HEPES + 150 mM NaCl (150 NaCl) | | | 16 | Ctl. vs. 150 NaCl | | ns | 0.276 |
| | 10 mM Glucrose + 10 mM HEPES + 250 mM NaCl (250 NaCl) | | | 18 | Ctl. vs. 250 NaCl | | ns | 0.1434 |
| | 10 mM Glucrose + 10 mM HEPES + 250 mM KCl (250 KCl) | | | 15 | Ctl. vs. 250 KCl | | ns | 0.189 |
| Fig. 5-E | 10 mM Glucrose + 10 mM HEPES + 250 mM NMDG-Cl (Ctl.) | *pAc5.1-IR11a, pAc5.1-IR25a, pAc5.1-GCaMP6M, pAc5.1-GFP* | IR11a + IR25a | 16 | | | | |
| | 10 mM Glucrose + 10 mM HEPES + 250 mM LiCl (250 LiCl) | | | 18 | Ctl. vs. 250 LiCl | | ns | 0.3581 |
| | 10 mM Glucrose + 10 mM HEPES + 50 mM NaCl (50 NaCl) | | | 15 | Ctl. vs. 50 NaCl | | ns | 0.2811 |
| | 10 mM Glucrose + 10 mM HEPES + 150 mM NaCl (150 NaCl) | | | 20 | Ctl. vs. 150 NaCl | | ns | 0.2358 |
| | 10 mM Glucrose + 10 mM HEPES + 250 mM NaCl (250 NaCl) | | | 19 | Ctl. vs. 250 NaCl | | *** | <0.0001 |
| | 10 mM Glucrose + 10 mM HEPES + 250 mM KCl (250 KCl) | | | 17 | Ctl. vs. 250 KCl | | ns | 0.5814 |
| Fig. 5-F | 10 mM Glucrose + 10 mM HEPES + 250 mM NMDG-Cl (Ctl.) | *pAc5.1-IR76b, pAc5.1-GCaMP6M, pAc5.1-GFP* | IR76b | 20 | | | | |
| | 10 mM Glucrose + 10 mM HEPES + 250 mM LiCl (250 LiCl) | | | 14 | Ctl. vs. 250 LiCl | | ns | 0.2901 |
| | 10 mM Glucrose + 10 mM HEPES + 50 mM NaCl (50 NaCl) | | | 18 | Ctl. vs. 50 NaCl | | ** | 0.0071 |
| | 10 mM Glucrose + 10 mM HEPES + 150 mM NaCl (150 NaCl) | | | 17 | Ctl. vs. 150 NaCl | | *** | 0.0001 |
| | 10 mM Glucrose + 10 mM HEPES + 250 mM NaCl (250 NaCl) | | | 20 | Ctl. vs. 250 NaCl | | * | 0.0304 |
| | 10 mM Glucrose + 10 mM HEPES + 250 mM KCl (250 KCl) | | | 16 | Ctl. vs. 250 KCl | | ns | 0.1086 |
| Fig. 5-G | 10 mM Glucrose + 10 mM HEPES + 250 mM NMDG-Cl (Ctl.) | *pAc5.1-IR25a, pAc5.1-IR76b, pAc5.1-GCaMP6M, pAc5.1-GFP* | IR25a + IR76b | 19 | | | | |
| | 10 mM Glucrose + 10 mM HEPES + 250 mM LiCl (250 LiCl) | | | 19 | Ctl. vs. 250 LiCl | | ns | 0.8786 |
| | 10 mM Glucrose + 10 mM HEPES + 50 mM NaCl (50 NaCl) | | | 20 | Ctl. vs. 50 NaCl | | ns | 0.84 |
| | 10 mM Glucrose + 10 mM HEPES + 150 mM NaCl (150 NaCl) | | | 18 | Ctl. vs. 150 NaCl | | * | 0.048 |
| | 10 mM Glucrose + 10 mM HEPES + 250 mM NaCl (250 NaCl) | | | 17 | Ctl. vs. 250 NaCl | | ns | 0.1067 |
| | 10 mM Glucrose + 10 mM HEPES + 250 mM KCl (250 KCl) | | | 19 | Ctl. vs. 250 KCl | | ns | 0.1458 |
| Fig. 5-H | 10 mM Glucrose + 10 mM HEPES + 250 mM NMDG-Cl (Ctl.) | *pAc5.1-IR11a, pAc5.1-IR76b, pAc5.1-GCaMP6M, pAc5.1-GFP* | IR11a + IR76b | 18 | | | | |
| | 10 mM Glucrose + 10 mM HEPES + 250 mM LiCl (250 LiCl) | | | 16 | Ctl. vs. 250 LiCl | | ns | 0.1434 |
| | 10 mM Glucrose + 10 mM HEPES + 50 mM NaCl (50 NaCl) | | | 18 | Ctl. vs. 50 NaCl | | ns | 0.1913 |
| | 10 mM Glucrose + 10 mM HEPES + 150 mM NaCl (150 NaCl) | | | 18 | Ctl. vs. 150 NaCl | | ** | 0.0038 |
| | 10 mM Glucrose + 10 mM HEPES + 250 mM NaCl (250 NaCl) | | | 18 | Ctl. vs. 250 NaCl | | ** | 0.0075 |
| | 10 mM Glucrose + 10 mM HEPES + 250 mM KCl (250 KCl) | | | 16 | Ctl. vs. 250 KCl | | ns | 0.5334 |
| Fig. 5-I | 10 mM Glucrose + 10 mM HEPES + 250 mM NMDG-Cl (Ctl.) | *pAc5.1-IR11a, pAc5.1-IR25a, pAc5.1-IR76b, pAc5.1-GCaMP6M, pAc5.1-GFP* | IR11a + 2 IR25a + IR76b | 11 | | | | |
| | 10 mM Glucrose + 10 mM HEPES + 50 mM LiCl (50 LiCl) | | | 16 | Ctl. vs. 50 LiCl | | ns | 0.3565 |
| | 10 mM Glucrose + 10 mM HEPES + 250 mM LiCl (250 LiCl) | | | 17 | Ctl. vs. 250 LiCl | | ** | 0.0029 |
| | 10 mM Glucrose + 10 mM HEPES + 50 mM NaCl (50 NaCl) | | | 19 | Ctl. vs. 50 NaCl | | ns | 0.1945 |
| | 10 mM Glucrose + 10 mM HEPES + 150 mM NaCl (150 NaCl) | | | 15 | Ctl. vs. 150 NaCl | | * | 0.0369 |
| | 10 mM Glucrose + 10 mM HEPES + 250 mM NaCl (250 NaCl) | | | 17 | Ctl. vs. 250 NaCl | | *** | 0.0005 |
| | 10 mM Glucrose + 10 mM HEPES + 250 mM KCl (250 KCl) | | | 14 | Ctl. vs. 250 KCl | | ns | 0.1201 |

**Data S6-Genotypes and statistical details for Fig.6**

| Figures | Foods or Test chemicals | Genotypes | Genotypes labeled in figures | N value | Culture conditions | Compare groups | Statistical methods | P value summary | P value |
|---|---|---|---|---|---|---|---|---|---|
| Fig. 6-A | 250 mM NaCl vs. $H_2O$ | w$^{1118}$ | w$^{1118}$ | 27 | salt deprived | salt deprived vs. salt fed | Mann-Whitney | *** | <0.0001 |
| | | w$^{1118}$ | w$^{1118}$ | 26 | salt fed | | | | |
| | | Poxn$^{70}$/+ | Poxn$^{70}$/+ | 20 | salt deprived | salt deprived vs. salt fed | | *** | <0.0001 |
| | | Poxn$^{70}$/+ | Poxn$^{70}$/+ | 20 | salt fed | | | | |
| | | Poxn$^{84}$/+ | Poxn$^{84}$/+ | 20 | salt deprived | salt deprived vs. salt fed | | *** | 0.0003 |
| | | Poxn$^{84}$/+ | Poxn$^{84}$/+ | 20 | salt fed | | | | |
| | | Poxn$^{70}$/Poxn$^{84}$ | Poxn$^{70}$/Poxn$^{84}$ | 21 | salt deprived | salt deprived vs. salt fed | | ns | 0.4587 |
| | | Poxn$^{70}$/Poxn$^{84}$ | Poxn$^{70}$/Poxn$^{84}$ | 22 | salt fed | | | | |
| Fig. 6-B | | wcs | wcs | 22 | salt deprived | salt deprived vs. salt fed | | *** | <0.0001 |
| | | wcs | wcs | 19 | salt fed | | | | |
| | | IR25a$^{2}$/+ | IR25a$^{2}$/+ | 23 | salt deprived | salt deprived vs. salt fed | | *** | <0.0001 |
| | | IR25a$^{2}$/+ | IR25a$^{2}$/+ | 24 | salt fed | | | | |
| | | IR25a$^{2}$ | IR25a$^{2}$ | 18 | salt deprived | salt deprived vs. salt fed | | ns | 0.2683 |
| | | IR25a$^{2}$ | IR25a$^{2}$ | 18 | salt fed | | | | |
| | | w$^{1118}$/w$^{1118}$;IR25a$^{2}$/IR25a$^{2}$,UAS-IR25a;+/+ | UAS-IR25a IR25a$^{2}$ | 13 | salt deprived | salt deprived vs. salt fed | | ns | 0.2877 |
| | | w$^{1118}$/w$^{1118}$;IR25a$^{2}$/IR25a$^{2}$,UAS-IR25a;+/+ | UAS-IR25a IR25a$^{2}$ | 13 | salt fed | | | | |
| | | w$^{1118}$/w$^{1118}$;IR25a$^{2}$/IR25a$^{2}$,IR25a-GAL4;+/+ | IR25a-GAL4 IR25a$^{2}$ | 14 | salt deprived | salt deprived vs. salt fed | | ns | 0.0959 |
| | | w$^{1118}$/w$^{1118}$;IR25a$^{2}$/IR25a$^{2}$,IR25a-GAL4;+/+ | IR25a-GAL4 IR25a$^{2}$ | 19 | salt fed | | | | |
| | | w$^{1118}$/w$^{1118}$;IR25a$^{2}$,UAS-IR25a/IR25a$^{2}$,IR25a-GAL4;+/+ | IR25a-GAL4$^{rescue}$ | 29 | salt deprived | salt deprived vs. salt fed | | *** | <0.0001 |
| | | w$^{1118}$/w$^{1118}$;IR25a$^{2}$,UAS-IR25a/IR25a$^{2}$,IR25a-GAL4;+/+ | IR25a-GAL4$^{rescue}$ | 25 | salt fed | | | | |
| Fig. 6-C | | w$^{1118}$/w$^{1118}$;IR25a$^{2}$/IR25a$^{2}$;IR11a-GAL4/+ | IR11a-GAL4 IR25a$^{2}$ | 17 | salt deprived | salt deprived vs. salt fed | | ns | 0.7613 |
| | | w$^{1118}$/w$^{1118}$;IR25a$^{2}$/IR25a$^{2}$;IR11a-GAL4/+ | IR11a-GAL4 IR25a$^{2}$ | 14 | salt fed | | | | |
| | | w$^{1118}$/w$^{1118}$;IR25a$^{2}$,UAS-IR25a/IR25a$^{2}$;IR11a-GAL4/+ | IR11a-GAL4$^{rescue}$ | 23 | salt deprived | salt deprived vs. salt fed | | *** | 0.0001 |
| | | w$^{1118}$/w$^{1118}$;IR25a$^{2}$,UAS-IR25a/IR25a$^{2}$;IR11a-GAL4/+ | IR11a-GAL4$^{rescue}$ | 23 | salt fed | | | | |
| | | w$^{1118}$/w$^{1118}$;IR25a$^{2}$/IR25a$^{2}$;IR7c-GAL4/+ | IR7c-GAL4 IR25a$^{2}$ | 11 | salt deprived | salt deprived vs. salt fed | | ns | 0.8328 |
| | | w$^{1118}$/w$^{1118}$;IR25a$^{2}$/IR25a$^{2}$;IR7c-GAL4/+ | IR7c-GAL4 IR25a$^{2}$ | 12 | salt fed | | | | |
| | | w$^{1118}$/w$^{1118}$;IR25a$^{2}$,UAS-IR25a/IR25a$^{2}$;IR7c-GAL4/+ | IR7c-GAL4$^{rescue}$ | 19 | salt deprived | salt deprived vs. salt fed | | ns | 0.1062 |
| | | w$^{1118}$/w$^{1118}$;IR25a$^{2}$,UAS-IR25a/IR25a$^{2}$;IR7c-GAL4/+ | IR7c-GAL4$^{rescue}$ | 21 | salt fed | | | | |
| Fig. 6-D | 250 mM NaCl | IR11a$^{4}$/+ | IR11a$^{4}$/+ | 5 | salt deprived | salt deprived vs. salt fed | | ** | 0.01 |
| | | | | 9 | salt fed | | | | |
| | | IR11a$^{4}$/IR11a$^{4}$ | IR11a$^{4}$ | 6 | salt deprived | | | ns | 0.1698 |
| | | | | 8 | salt fed | | | | |
| | | IR11a$^{4}$/ IR11a$^{4}$;UAS-IR11a/+ | UAS-IR11a IR11a$^{4}$ | 5 | salt deprived | | | ns | 0.514 |
| | | | | 6 | salt fed | | | | |
| | | IR11a$^{4}$/ IR11a$^{4}$;+/+;IR11a-GAL4/+ | IR11a-GAL4 IR11a$^{4}$ | 6 | salt deprived | | | ns | 0.9719 |
| | | | | 7 | salt fed | | | | |
| | | IR11a$^{4}$/ IR11a$^{4}$;UAS-IR11a/+;IR11a-GAL4/+ | IR11a-GAL4$^{rescue}$ | 11 | salt deprived | | | *** | <0.0001 |
| | | | | 15 | salt fed | | | | |
| Fig. 6-E | 250 mM NaCl | IR11a$^{4}$/+ | IR11a$^{4}$/+ | 26 | salt deprived | salt deprived vs. salt fed | | *** | <0.0001 |
| | | | | 26 | salt fed | | | | |
| | | IR11a$^{4}$/IR11a$^{4}$ | IR11a$^{4}$ | 25 | salt deprived | | | ns | 0.292 |
| | | | | 29 | salt fed | | | | |
| | | IR11a$^{4}$/ IR11a$^{4}$;UAS-IR11a/+ | UAS-IR11a IR11a$^{4}$ | 24 | salt deprived | | | ns | 0.7683 |
| | | | | 16 | salt fed | | | | |
| | | IR11a$^{4}$/ IR11a$^{4}$;+/+;IR11a-GAL4/+ | IR11a-GAL4 IR11a$^{4}$ | 21 | salt deprived | | | ns | 0.3233 |
| | | | | 20 | salt fed | | | | |
| | | IR11a$^{4}$/ IR11a$^{4}$;UAS-IR11a/+;IR11a-GAL4/+ | IR11a-GAL4$^{rescue}$ | 24 | salt deprived | | | *** | <0.0001 |
| | | | | 18 | salt fed | | | | |
| Fig. 6-F | 20 mM Sucrose | IR11a$^{4}$/+ | IR11a$^{4}$/+ | 26 | salt deprived | salt deprived vs. salt fed | | * | 0.0152 |
| | | | | 26 | salt fed | | | | |
| | | IR11a$^{4}$/IR11a$^{4}$ | IR11a$^{4}$ | 25 | salt deprived | | | ns | 0.4554 |
| | | | | 29 | salt fed | | | | |
| | | IR11a$^{4}$/ IR11a$^{4}$;UAS-IR11a/+ | UAS-IR11a IR11a$^{4}$ | 24 | salt deprived | | | ns | 0.0824 |
| | | | | 16 | salt fed | | | | |
| | | IR11a$^{4}$/ IR11a$^{4}$;+/+;IR11a-GAL4/+ | IR11a-GAL4 IR11a$^{4}$ | 21 | salt deprived | | | * | 0.0173 |
| | | | | 20 | salt fed | | | | |
| | | IR11a$^{4}$/ IR11a$^{4}$;UAS-IR11a/+;IR11a-GAL4/+ | IR11a-GAL4$^{rescue}$ | 24 | salt deprived | | | ns | 0.8849 |
| | | | | 18 | salt fed | | | | |
| Fig. 6-G | $H_2O$ | w$^{1118}$/w$^{1118}$;GR33a$^{GAL4}$,UAS-GCaMP6m/+ | GR33a > GCaMP6M | 8 | salt deprived | salt deprived vs. salt fed | | ns | 0.673 |
| | | | | 9 | salt fed | | | | |
| | 250 mM NaCl | | | 13 | salt deprived | | | ** | 0.001 |
| | | | | 15 | salt fed | | | | |
| | 250 mM KCl | | | 8 | salt deprived | | | ns | 0.7921 |
| | | | | 12 | salt fed | | | | |
| | 1 mM Denatonium | | | 11 | salt deprived | | | ns | 0.7564 |
| | | | | 10 | salt fed | | | | |
| Fig. 6-H | $H_2O$ | w$^{1118}$/w$^{1118}$;UAS-GCaMP6m/+ ; IR7c-GAL4/+ | IR7c > GCaMP6m | 11 | salt deprived | salt deprived vs. salt fed | | ns | 0.5027 |
| | | | | 9 | salt fed | | | | |
| | 250 mM NaCl | | | 9 | salt deprived | | | ns | 0.211 |
| | | | | 10 | salt fed | | | | |
| | 250 mM KCl | | | 8 | salt deprived | | | ns | 0.4807 |
| | | | | 9 | salt fed | | | | |
| Fig. 6-I | 250 mM NaCl | w$^{1118}$/w$^{1118}$;IR25a$^{2}$,GR33a$^{GAL4}$/UAS-GCaMP6m | R25a control | 9 | salt deprived | salt deprived vs. salt fed | | *** | 0.0002 |
| | | | | 9 | salt fed | | | | |
| | | w$^{1118}$/w$^{1118}$;IR25a$^{2}$,GR33a$^{GAL4}$/IR25a$^{2}$;UAS-GCaMP6m/+ | R25a mutants | 8 | salt deprived | | | ns | 0.743 |
| | | | | 9 | salt fed | | | | |
| | | w$^{1118}$/w$^{1118}$;IR25a$^{2}$,GR33a$^{GAL4}$/IR25a$^{2}$,UAS-IR25a;UAS-GCaMP6m/+ | R25a rescue | 8 | salt deprived | | | *** | <0.0001 |
| | | | | 9 | salt fed | | | | |
| Fig. 6-J | 250 mM NaCl | IR11a$^{4}$/+;GR33a$^{GAL4}$,UAS-GCaMP6m/+ | R11a control | 11 | salt deprived | salt deprived vs. salt fed | | ** | 0.0086 |
| | | | | 12 | salt fed | | | | |
| | | IR11a$^{4}$/ IR11a$^{4}$;GR33a$^{GAL4}$,UAS-GCaMP6m/+ | R11a mutants | 11 | salt deprived | | | ns | 0.3494 |
| | | | | 10 | salt fed | | | | |
| | | IR11a$^{4}$/ IR11a$^{4}$;GR33a$^{GAL4}$,UAS-GCaMP6m/UAS-IR11a | R11a rescue | 12 | salt deprived | | | *** | 0.0001 |
| | | | | 13 | salt fed | | | | |

| Data S7-Genotypes and statistical details for Fig.S1 | | | | | | | |
|---|---|---|---|---|---|---|---|
| **Figures** | **Genotypes** | **Genotypes labeled in figures** | **N value** | **Compare groups** | **Statistical methods** | **P value summary** | **P value** |
| Fig. S1-A | *nSyb-GAL4,Tub-GAL80ts / + 21℃* | *Ctl.* | 23 | 21℃ vs. 31℃ | Mann-Whitney | ns | 0.3636 |
| | *nSyb-GAL4,Tub-GAL80ts / + 31℃* | | 24 | | | | |
| | *nSyb-GAL4,Tub-GAL80ts>IR7b-RNAi 21℃* | *IR7b* | 22 | | | ns | 0.8403 |
| | *nSyb-GAL4,Tub-GAL80ts>IR7b-RNAi 31℃* | | 24 | | | | |
| | *nSyb-GAL4,Tub-GAL80ts>IR8a-RNAi 21℃* | *IR8a* | 20 | | | ns | 0.8565 |
| | *nSyb-GAL4,Tub-GAL80ts>IR8a-RNAi 31℃* | | 20 | | | | |
| | *nSyb-GAL4,Tub-GAL80ts>IR21a-RNAi 21℃* | *IR21a* | 15 | | | ns | 0.267 |
| | *nSyb-GAL4,Tub-GAL80ts>IR21a-RNAi 31℃* | | 19 | | | | |
| | *nSyb-GAL4,Tub-GAL80ts>IR48c-RNAi 21℃* | *IR48c* | 27 | | | ns | 0.1288 |
| | *nSyb-GAL4,Tub-GAL80ts>IR48c-RNAi 31℃* | | 20 | | | | |
| | *nSyb-GAL4,Tub-GAL80ts>IR52d-RNAi 21℃* | *IR52d* | 20 | | | ns | 0.6346 |
| | *nSyb-GAL4,Tub-GAL80ts>IR52d-RNAi 31℃* | | 20 | | | | |
| | *nSyb-GAL4,Tub-GAL80ts>IR56b-RNAi 21℃* | *IR56b* | 22 | | | *** | <0.001 |
| | *nSyb-GAL4,Tub-GAL80ts>IR56b-RNAi 31℃* | | 22 | | | | |
| | *nSyb-GAL4,Tub-GAL80ts>IR56d-RNAi 21℃* | *IR56d* | 19 | | | ns | 0.5734 |
| | *nSyb-GAL4,Tub-GAL80ts>IR56d-RNAi 31℃* | | 20 | | | | |
| | *nSyb-GAL4,Tub-GAL80ts>IR62a-RNAi 21℃* | *IR62a* | 10 | | | ns | 0.7663 |
| | *nSyb-GAL4,Tub-GAL80ts>IR62a-RNAi 31℃* | | 22 | | | | |
| | *nSyb-GAL4,Tub-GAL80ts>IR68a-RNAi 21℃* | *IR68a* | 19 | | | *** | 0.0004 |
| | *nSyb-GAL4,Tub-GAL80ts>IR68a-RNAi 31℃* | | 20 | | | | |
| | *nSyb-GAL4,Tub-GAL80ts>IR75a-RNAi 21℃* | *IR75a* | 20 | | | ns | 0.3247 |
| | *nSyb-GAL4,Tub-GAL80ts>IR75a-RNAi 31℃* | | 22 | | | | |
| | *nSyb-GAL4,Tub-GAL80ts>IR75c-RNAi 21℃* | *IR75c* | 20 | | | ns | 0.5541 |
| | *nSyb-GAL4,Tub-GAL80ts>IR75c-RNAi 31℃* | | 24 | | | | |
| | *nSyb-GAL4,Tub-GAL80ts>IR76a-RNAi 21℃* | *IR76a* | 20 | | | * | 0.284 |
| | *nSyb-GAL4,Tub-GAL80ts>IR76a-RNAi 31℃* | | 21 | | | | |
| | *nSyb-GAL4,Tub-GAL80ts>IR94a-RNAi 21℃* | *IR94a* | 16 | | | ns | 0.2405 |
| | *nSyb-GAL4,Tub-GAL80ts>IR94a-RNAi 31℃* | | 18 | | | | |
| Fig. S1-B | *nSyb-GAL4,Tub-GAL80ts / + 21℃* | *Ctl.* | 23 | 21℃ vs. 31℃ | Mann-Whitney | ns | 0.3636 |
| | *nSyb-GAL4,Tub-GAL80ts / + 31℃* | | 24 | | | | |
| | *nSyb-GAL4,Tub-GAL80ts>Trp-RNAi 21℃* | *Trp* | 20 | | | ns | 0.2218 |
| | *nSyb-GAL4,Tub-GAL80ts>Trp-RNAi 31℃* | | 17 | | | | |
| | *nSyb-GAL4,Tub-GAL80ts>Trpa1-RNAi 21℃* | *Trpa1* | 25 | | | ns | 0.5894 |
| | *nSyb-GAL4,Tub-GAL80ts>Trpa1-RNAi 31℃* | | 19 | | | | |
| | *nSyb-GAL4,Tub-GAL80ts>Trpm1-RNAi 21℃* | *Trpm1* | 20 | | | ns | 0.634 |
| | *nSyb-GAL4,Tub-GAL80ts>Trpm1-RNAi 31℃* | | 18 | | | | |
| | *nSyb-GAL4,Tub-GAL80ts>Trpγ-RNAi 21℃* | *Trpγ* | 22 | | | ns | 0.7215 |
| | *nSyb-GAL4,Tub-GAL80ts>Trpγ-RNAi 31℃* | | 18 | | | | |
| | *nSyb-GAL4,Tub-GAL80ts / + 21℃* | *Ctl.* | 23 | | | ns | 0.3636 |
| | *nSyb-GAL4,Tub-GAL80ts / + 31℃* | | 24 | | | | |
| | *nSyb-GAL4,Tub-GAL80ts>Ppk5-RNAi 21℃* | *Ppk5* | 28 | | | ns | 0.8554 |
| | *nSyb-GAL4,Tub-GAL80ts>Ppk5-RNAi 31℃* | | 30 | | | | |
| | *nSyb-GAL4,Tub-GAL80ts>Ppk6-RNAi 21℃* | *Ppk6* | 19 | | | ns | 0.1271 |
| | *nSyb-GAL4,Tub-GAL80ts>Ppk6-RNAi 31℃* | | 19 | | | | |
| | *nSyb-GAL4,Tub-GAL80ts>Ppk7-RNAi 21℃* | *Ppk7* | 20 | | | ns | 0.176 |
| | *nSyb-GAL4,Tub-GAL80ts>Ppk7-RNAi 31℃* | | 20 | | | | |
| | *nSyb-GAL4,Tub-GAL80ts>Ppk8-RNAi 21℃* | *Ppk8* | 20 | | | ns | 0.3427 |
| | *nSyb-GAL4,Tub-GAL80ts>Ppk8-RNAi 31℃* | | 20 | | | | |

| | | | | | | | |
|---|---|---|---|---|---|---|---|
| Fig. S1-C | *nSyb-GAL4,Tub-GAL80ts>Ppk9-RNAi 21℃* | *Ppk9* | 22 | 21℃ vs. 31℃ | Mann-Whitney | ns | 0.1948 |
| | *nSyb-GAL4,Tub-GAL80ts>Ppk9-RNAi 31℃* | | 20 | | | | |
| | *nSyb-GAL4,Tub-GAL80ts>Ppk11-RNAi 21℃* | *Ppk11* | 28 | | | * | 0.0293 |
| | *nSyb-GAL4,Tub-GAL80ts>Ppk11-RNAi 31℃* | | 22 | | | | |
| | *nSyb-GAL4,Tub-GAL80ts>Ppk13-RNAi 21℃* | *Ppk13* | 20 | | | ** | 0.0097 |
| | *nSyb-GAL4,Tub-GAL80ts>Ppk13-RNAi 31℃* | | 19 | | | | |
| | *nSyb-GAL4,Tub-GAL80ts>Ppk14-RNAi 21℃* | *Ppk14* | 20 | | | ns | 0.0827 |
| | *nSyb-GAL4,Tub-GAL80ts>Ppk14-RNAi 31℃* | | 29 | | | | |
| | *nSyb-GAL4,Tub-GAL80ts>Ppk15-RNAi 21℃* | *Ppk15* | 30 | | | ns | 0.7722 |
| | *nSyb-GAL4,Tub-GAL80ts>Ppk15-RNAi 31℃* | | 20 | | | | |
| | *nSyb-GAL4,Tub-GAL80ts>Ppk16-RNAi 21℃* | *Ppk16* | 20 | | | * | 0.0254 |
| | *nSyb-GAL4,Tub-GAL80ts>Ppk16-RNAi 31℃* | | 20 | | | | |
| | *nSyb-GAL4,Tub-GAL80ts>Ppk18-RNAi 21℃* | *Ppk18* | 20 | | | * | 0.0314 |
| | *nSyb-GAL4,Tub-GAL80ts>Ppk18-RNAi 31℃* | | 20 | | | | |
| | *nSyb-GAL4,Tub-GAL80ts>Ppk19-RNAi 21℃* | *Ppk19* | 19 | | | ns | 0.0581 |
| | *nSyb-GAL4,Tub-GAL80ts>Ppk19-RNAi 31℃* | | 17 | | | | |
| | *nSyb-GAL4,Tub-GAL80ts>Ppk20-RNAi 21℃* | *Ppk20* | 19 | | | ns | 0.9278 |
| | *nSyb-GAL4,Tub-GAL80ts>Ppk20-RNAi 31℃* | | 20 | | | | |
| | *nSyb-GAL4,Tub-GAL80ts>Ppk21-RNAi 21℃* | *Ppk21* | 20 | | | ns | 0.9941 |
| | *nSyb-GAL4,Tub-GAL80ts>Ppk21-RNAi 31℃* | | 20 | | | | |
| | *nSyb-GAL4,Tub-GAL80ts>Ppk22-RNAi 21℃* | *Ppk22* | 20 | | | ns | 0.157 |
| | *nSyb-GAL4,Tub-GAL80ts>Ppk22-RNAi 31℃* | | 20 | | | | |
| | *nSyb-GAL4,Tub-GAL80ts>Ppk23-RNAi 21℃* | *Ppk23* | 20 | | | ns | 0.928 |
| | *nSyb-GAL4,Tub-GAL80ts>Ppk23-RNAi 31℃* | | 19 | | | | |
| | *nSyb-GAL4,Tub-GAL80ts>Ppk24-RNAi 21℃* | *Ppk24* | 19 | | | ns | 0.1756 |
| | *nSyb-GAL4,Tub-GAL80ts>Ppk24-RNAi 31℃* | | 20 | | | | |
| | *nSyb-GAL4,Tub-GAL80ts>Ppk25-RNAi 21℃* | *Ppk25* | 27 | | | ns | 0.1873 |
| | *nSyb-GAL4,Tub-GAL80ts>Ppk25-RNAi 31℃* | | 10 | | | | |
| | *nSyb-GAL4,Tub-GAL80ts>Ppk28-RNAi 21℃* | *Ppk28* | 30 | | | * | 0.0108 |
| | *nSyb-GAL4,Tub-GAL80ts>Ppk28-RNAi 31℃* | | 19 | | | | |
| | *nSyb-GAL4,Tub-GAL80ts>Ppk29-RNAi 21℃* | *Ppk29* | 20 | | | ns | 0.5487 |
| | *nSyb-GAL4,Tub-GAL80ts>Ppk29-RNAi 31℃* | | 19 | | | | |
| | *nSyb-GAL4,Tub-GAL80ts>Ppk30-RNAi 21℃* | *Ppk30* | 30 | | | ns | 0.4879 |
| | *nSyb-GAL4,Tub-GAL80ts>Ppk30-RNAi 31℃* | | 28 | | | | |
| | *nSyb-GAL4,Tub-GAL80ts>Ppk31-RNAi 21℃* | *Ppk31* | 29 | | | ns | 0.9049 |
| | *nSyb-GAL4,Tub-GAL80ts>Ppk31-RNAi 31℃* | | 20 | | | | |
| Fig. S1-D | *nSyb-GAL4,Tub-GAL80ts / + 21℃* | *Ctl.* | 23 | 21℃ vs. 31℃ | Mann-Whitney | ns | 0.3636 |
| | *nSyb-GAL4,Tub-GAL80ts / + 31℃* | | 24 | | | | |
| | *nSyb-GAL4,Tub-GAL80ts>GR22b-RNAi 21℃* | *GR22b* | 29 | | | ns | 0.4941 |
| | *nSyb-GAL4,Tub-GAL80ts>GR22b-RNAi 31℃* | | 24 | | | | |
| | *nSyb-GAL4,Tub-GAL80ts>GR22c-RNAi 21℃* | *GR22c* | 10 | | | ns | 0.5836 |
| | *nSyb-GAL4,Tub-GAL80ts>GR22c-RNAi 31℃* | | 12 | | | | |
| | *nSyb-GAL4,Tub-GAL80ts>GR22d-RNAi 21℃* | *GR22d* | 21 | | | ns | 0.9926 |
| | *nSyb-GAL4,Tub-GAL80ts>GR22d-RNAi 31℃* | | 12 | | | | |
| | *nSyb-GAL4,Tub-GAL80ts>GR22f-RNAi 21℃* | *GR22f* | 20 | | | ns | 0.2109 |
| | *nSyb-GAL4,Tub-GAL80ts>GR22f-RNAi 31℃* | | 9 | | | | |
| | *nSyb-GAL4,Tub-GAL80ts>GR32a-RNAi 21℃* | *GR32a* | 19 | | | ns | 0.9138 |
| | *nSyb-GAL4,Tub-GAL80ts>GR32a-RNAi 31℃* | | 19 | | | | |
| | *nSyb-GAL4,Tub-GAL80ts>GR36c-RNAi 21℃* | *GR36c* | 20 | | | ns | 0.5119 |
| | *nSyb-GAL4,Tub-GAL80ts>GR36c-RNAi 31℃* | | 17 | | | | |
| | *nSyb-GAL4,Tub-GAL80ts>GR43a-RNAi 21℃* | *GR43a* | 21 | | | ** | 0.0054 |
| | *nSyb-GAL4,Tub-GAL80ts>GR43a-RNAi 31℃* | | 13 | | | | |
| | *nSyb-GAL4,Tub-GAL80ts>GR47b-RNAi 21℃* | *GR47b* | 18 | | | ns | 0.071 |
| | *nSyb-GAL4,Tub-GAL80ts>GR47b-RNAi 31℃* | | 20 | | | | |
| | *nSyb-GAL4,Tub-GAL80ts>GR57a-RNAi 21℃* | *GR57a* | 20 | | | ns | 0.5258 |
| | *nSyb-GAL4,Tub-GAL80ts>GR57a-RNAi 31℃* | | 20 | | | | |
| | *nSyb-GAL4,Tub-GAL80ts>GR59d-RNAi 21℃* | *GR59d* | 16 | | | * | 0.028 |
| | *nSyb-GAL4,Tub-GAL80ts>GR59d-RNAi 31℃* | | 19 | | | | |
| | *nSyb-GAL4,Tub-GAL80ts>GR59e-RNAi 21℃* | *GR59e* | 16 | | | ns | 0.5377 |
| | *nSyb-GAL4,Tub-GAL80ts>GR59e-RNAi 31℃* | | 18 | | | | |
| | *nSyb-GAL4,Tub-GAL80ts>GR59f-RNAi 21℃* | *GR59f* | 33 | | | ns | 0.3042 |
| | *nSyb-GAL4,Tub-GAL80ts>GR59f-RNAi 31℃* | | 30 | | | | |
| | *nSyb-GAL4,Tub-GAL80ts>GR64b-RNAi 21℃* | *GR64b* | 21 | | | ns | 0.1941 |
| | *nSyb-GAL4,Tub-GAL80ts>GR64b-RNAi 31℃* | | 20 | | | | |
| | *nSyb-GAL4,Tub-GAL80ts>GR64c-RNAi 21℃* | *GR64c* | 20 | | | ** | 0.0071 |
| | *nSyb-GAL4,Tub-GAL80ts>GR64c-RNAi 31℃* | | 29 | | | | |
| | *nSyb-GAL4,Tub-GAL80ts>GR64d-RNAi 21℃* | *GR64d* | 19 | | | ns | 0.2111 |
| | *nSyb-GAL4,Tub-GAL80ts>GR64d-RNAi 31℃* | | 20 | | | | |
| | *nSyb-GAL4,Tub-GAL80ts>GR64f-RNAi 21℃* | *GR64f* | 28 | | | ns | 0.2734 |
| | *nSyb-GAL4,Tub-GAL80ts>GR64f-RNAi 31℃* | | 20 | | | | |

| | | | | | | | |
|---|---|---|---|---|---|---|---|
| | *nSyb-GAL4,Tub-GAL80ts>GR77a-RNAi 21℃* | *GR77a* | 18 | | | ns | 0.1005 |
| | *nSyb-GAL4,Tub-GAL80ts>GR77a-RNAi 31℃* | | 26 | | | | |
| | *nSyb-GAL4,Tub-GAL80ts>GR92a-RNAi 21℃* | *GR92a* | 30 | | | ns | 0.1777 |
| | *nSyb-GAL4,Tub-GAL80ts>GR92a-RNAi 31℃* | | 25 | | | | |
| | *nSyb-GAL4,Tub-GAL80ts>GR97a-RNAi 21℃* | *GR97a* | 30 | | | ns | 0.0516 |
| | *nSyb-GAL4,Tub-GAL80ts>GR97a-RNAi 31℃* | | 29 | | | | |
| | *nSyb-GAL4,Tub-GAL80ts>GR98a-RNAi 21℃* | *GR98a* | 14 | | | ns | 0.0818 |
| | *nSyb-GAL4,Tub-GAL80ts>GR98a-RNAi 31℃* | | 19 | | | | |
| | *nSyb-GAL4,Tub-GAL80ts>GR98c-RNAi 21℃* | *GR98c* | 18 | | | * | 0.0363 |
| | *nSyb-GAL4,Tub-GAL80ts>GR98c-RNAi 31℃* | | 15 | | | | |

| Data S8-Genotypes and statistical details for Fig.S2 | | | |
|---|---|---|---|
| **Figures** | **Foods** | **Genotypes** | **N value** |
| Fig. S2-A | **standard food** | *$w^{1118}$/$w^{1118}$;UAS-mcd8:GFP/+;IR11a-GAL4/+* | 20 |
| Fig. S2-B | **standard food** | *$w^{1118}$/$w^{1118}$;UAS-mcd8:GFP/+;IR11a-GAL4/+* | 2 |
| Fig. S2-C | **standard food** | *$w^{1118}$/$w^{1118}$;UAS-mcd8:GFP/+;IR11a-GAL4/+* | 4 |
| Fig. S2-D | **standard food** | *$w^{1118}$/$w^{1118}$;UAS-mcd8:GFP/+;+/+* | 2 |
| | | *$w^{1118}$/$w^{1118}$;UAS-mcd8:GFP/+;IR11a-GAL4/+* | 2 |

| Data S9-Genotypes and statistical details for Fig.S3 | | | | | | | | |
|---|---|---|---|---|---|---|---|---|
| **Figures** | **Test chemicals** | **Genotypes** | **Genotypes labeled in figures** | **N value** | **Compare groups** | **Statistical methods** | **P value summary** | **P value** |
| Fig. S3-A,B | $H_2O$ | *$w^{1118}$/$w^{1118}$;;IR11a-GAL4,UAS-GCaMP6m/+* | *IR11a > GCaMP6M* | 16 | | Mann-Whitney | | |
| | 50 mM NaCl | | | 15 | $H_2O$ vs. 50 mM NaCl | | ns | 0.3791 |
| | 150 mM NaCl | | | 15 | $H_2O$ vs. 150 mM NaCl | | ns | 0.1102 |
| | 250 mM NaCl | | | 15 | $H_2O$ vs. 250 mM NaCl | | ** | 0.0063 |
| | 500 mM NaCl | | | 18 | $H_2O$ vs. 500 mM NaCl | | *** | 0.0003 |
| Fig. S3-C | $H_2O$ | *$w^{1118}$/$w^{1118}$;;IR11a-GAL4,UAS-GCaMP6m/+* | *IR11a > GCaMP6M* | 16 | | Mann-Whitney | | |
| | 50 mM LiCl | | | 11 | $H_2O$ vs. 50 mM LiCl | | ns | 0.1759 |
| | 50 mM KCl | | | 14 | $H_2O$ vs. 50 mM KCl | | ns | 0.5253 |
| | 250 mM NMDG-Cl | | | 8 | $H_2O$ vs. 250 mM NMDG-Cl | | ns | 0.8128 |
| | 250 mM LiCl | | | 12 | $H_2O$ vs. 250 mM LiCl | | ns | 0.9331 |
| | 250 mM NaCl | | | 13 | $H_2O$ vs. 250 mM NaCl | | *** | 0.0003 |
| | 250 mM KCl | | | 16 | $H_2O$ vs. 250 mM KCl | | *** | 0.0009 |
| | 50 mM $CaCl_2$ | | | 8 | $H_2O$ vs. 50 mM $CaCl_2$ | | ns | 0.2861 |
| | 10 mM Denatonium | | | 14 | $H_2O$ vs. 10 mM Denatonium | | *** | <0.001 |
| | $H_2O$ | *$w^{1118}$/$w^{1118}$;;IR11a-GAL4,UAS-GCaMP6m/+* | *$IR11a^4$ / +* | 16 | *$IR11a^4$/+ vs. $IR11a^4$* | Kruskal-Wallis, with Dunn's multiple comparisons test | ns | >0.9999 |
| | | *$IR11a^4$ / $IR11a^4$;;IR11a-GAL4,UAS-GCaMP6m/+* | *$IR11a^4$* | 10 | *$IR11a^4$/+ vs. $IR11a^4$ rescue* | | ns | 0.3424 |
| | | *$IR11a^4$ / $IR11a^4$;UAS-IR11a/+;IR11a-GAL4,UAS-GCaMP6m/+* | *$IR11a^4$* rescue | 8 | *$IR11a^4$ vs. $IR11a^4$ rescue* | | ns | >0.9999 |
| | 250 mM NaCl | *$w^{1118}$/$w^{1118}$;;IR11a-GAL4,UAS-GCaMP6m/+* | *$IR11a^4$ / +* | 13 | *$IR11a^4$/+ vs. $IR11a^4$* | Kruskal-Wallis, with Dunn's multiple comparisons test | * | 0.024 |
| | | *$IR11a^4$ / $IR11a^4$;;IR11a-GAL4,UAS-GCaMP6m/+* | *$IR11a^4$* | 8 | *$IR11a^4$/+ vs. $IR11a^4$ rescue* | | ns | 0.9087 |
| | | *$IR11a^4$ / $IR11a^4$;UAS-IR11a/+;IR11a-GAL4,UAS-GCaMP6m/+* | *$IR11a^4$* rescue | 12 | *$IR11a^4$ vs. $IR11a^4$ rescue* | | ** | 0.0017 |
| | 250 mM KCl | *$w^{1118}$/$w^{1118}$;;IR11a-GAL4,UAS-GCaMP6m/+* | *$IR11a^4$ / +* | 16 | *$IR11a^4$/+ vs. $IR11a^4$* | Mann-Whitney | ns | 0.2907 |
| | | *$IR11a^4$ / $IR11a^4$;;IR11a-GAL4,UAS-GCaMP6m/+* | *$IR11a^4$* | 8 | | | | |
| | 50 mM $CaCl_2$ | *$w^{1118}$/$w^{1118}$;;IR11a-GAL4,UAS-GCaMP6m/+* | *$IR11a^4$ / +* | 8 | *$IR11a^4$/+ vs. $IR11a^4$* | | ns | 0.5737 |
| | | *$IR11a^4$ / $IR11a^4$;;IR11a-GAL4,UAS-GCaMP6m/+* | *$IR11a^4$* | 8 | | | | |
| | 10 mM Denatonium | *$w^{1118}$/$w^{1118}$;;IR11a-GAL4,UAS-GCaMP6m/+* | *$IR11a^4$ / +* | 14 | *$IR11a^4$/+ vs. $IR11a^4$* | | ns | 0.1936 |
| | | *$IR11a^4$ / $IR11a^4$;;IR11a-GAL4,UAS-GCaMP6m/+* | *$IR11a^4$* | 14 | | | | |

**Data S10-Genotypes and statistical details for Fig.S4**

| Figures | Genotypes | N value | Tissues | Genes | Culture conditions | Compare groups | Statistical methods | P value summary | P value |
|---|---|---|---|---|---|---|---|---|---|
| Fig. S4-A | $w^{1118}$ | 9 | labellum | IR11a | salt deprived | salt deprived vs. salt fed | Mann-Whitney | ns | 0.0625 |
| | | 9 | | | salt fed | | | | |
| | | 9 | | IR25a | salt deprived | | | | 0.7962 |
| | | 9 | | | salt fed | | | ns | |
| | | 3 | | IR76b | salt deprived | | | | 0.1 |
| | | 9 | | | salt fed | | | ns | |
| | | 9 | | IR7c | salt deprived | | | ns | 0.7304 |
| | | 3 | | | salt fed | | | | |
| Fig. S4-B1-B3 | $GR33a^{GAL4}/+;UAS\text{-}RFP/+$ | 15±3 | | | salt deprived | | | | |
| Fig. S4-C1-C3 | | 21±3 | | | salt fed | | | | |
| Fig. S4-D | | 12 | f5b | | salt deprived | salt deprived vs. salt fed | Mann-Whitney | ns | 0.8683 |
| | | 21 | | | salt fed | | | | |
| Fig. S4-E | | 15 | f5s | | salt deprived | | | ns | 0.216 |
| | | 18 | | | salt fed | | | | |
